# Loss of *RND3/RHOE* controls entosis through *LAMP1* expression in hepatocellular carcinoma

**DOI:** 10.1101/2023.09.01.555890

**Authors:** Sara Basbous, Lydia Dif, Camille Dantzer, Sylvaine Di-Tommaso, Jean-William Dupuy, Paulette Bioulac-Sage, Anne-Aurélie Raymond, Chantal Desdouets, Frédéric Saltel, Violaine Moreau

## Abstract

Entosis is a process that leads to the formation of cell-in-cell structures commonly found in cancers. Here, we identified entosis in hepatocellular carcinoma and the loss of Rnd3 as an efficient inducer of this mechanism. We characterized the different stages and the molecular regulators of entosis induced after Rnd3 silencing. We demonstrated that this process depends on RhoA/ROCK pathway, but not on E-cadherin. The proteomic profiling of entotic cells allowed us to identify LAMP1 as a protein upregulated by Rnd3 silencing and implicated not only in the degradation final stage of entosis, but also in the full mechanism. Moreover, we found a positive correlation between the presence of entotic cells and the metastatic potential of tumors in human patient samples. Altogether, these data suggest the involvement of entosis in liver tumor progression and highlight a new perspective for entosis analysis in medicine research as a novel therapeutic target.

## Introduction

The presence of cell-in-cell (CIC) structures has been observed in a wide range of human cancers and mainly associated with poor prognosis *(1–7)*. CIC formation involves a dynamic interaction between an outer and an inner cell. It can be the result of different types of mechanisms involving either homotypic or heterotypic interactions and initiated by either the outer (endocytic CIC) or the inner (invasive CIC) cell *(8)*. Entosis is defined as a homotypic invasion occurring between tumor cells of the same type where one living cell is internalized inside another one. After internalization by the outer cell, the inner cell is found in the entotic vacuole *(10)*, where it will be degraded generally by a non-apoptotic cell death, autophagy involving the lysosome fusion. Entosis was discovered *in vitro* in breast cancer cells (MCF7) cultured under different stress factors such as extracellular matrix detachment, glucose starvation or ultraviolet radiation *(11–13)*. Entosis was also described in cells with prolonged aberrant mitosis to eliminate by engulfment the aneuploid progenies, maintaining thus the genome integrity *(14)*. Mechanistically, entosis was found dependent on cell-cell adhesion molecules such as cadherins and driven by imbalances in actomyosin contractility which is mainly under the control of the RhoA/ROCK pathway *(11, 15)*.

Rnd3/RhoE is an atypical RhoGTPase protein and a negative regulator of the Rho/ROCK pathway. Others and we previously described Rnd3 as a tumor suppressor in liver cancer *(16, 17)*. Indeed, *RND3* expression is downregulated in human hepatocellular carcinoma (HCC) samples compared to non-tumoral liver tissues and correlated with poor survival rates *(18, 19)*. It is implicated in cancer development through the regulation of cell growth and invasion *(18, 20)*. While working on the role of Rnd3 in HCC, we observed CIC structures, prompting us to study this phenomenon in liver cancer cells. Here, we reported that HCC cells are susceptible to form entosis after Rnd3 down-regulation in cultured liver cancer cell lines and in xenografts in mice. Entosis is highly dependent on RhoA/ROCK pathway, but not on E-cadherin. We also found that Rnd3 loss leads to Lamp-1 up-regulation, required for the internalization and degradation of the entotic cell. Finally, we related that entosis is rare in human HCC tissues, but associated to poor prognosis. Our results suggest that entotic engulfment induced by the loss of Rnd3 in HCC promotes liver tumor progression.

## Results

### Liver cancer cells are prone to perform entosis

In order to investigate entosis in liver cancer, cultured cells (Hep3B, Huh7, Huh6 or HepG2) were subjected to stress conditions known to favor entosis in breast cancer cells such as nutrient deprivation *(12)* or matrix detachment *(11)*. MCF-7 human breast cancer cells were used as a positive control. Cell internalization was analyzed and quantified by confocal microscopy after immunostaining of nuclei, F-actin and β-catenin to visualize nucleus deformation, cell periphery and shape respectively. The outer cell shows a crescent shape nucleus, whereas the inner cell is rounded and retains its plasma membrane (**Figure 1A**). In normal conditions, entotic structures can be observed in about 1-3% of total adherent liver cancer cells. After nutrient deprivation, a significant increase in the percentage of entotic cells is observed in all four cell lines, with up to 10% of entotic cells in HepG2 and Huh6 cells, when compared to normal culture conditions (**Figure 1B**). Matrix detachment is also described as an inducer of entosis in MCF-7 cells *(11)*. In order to test this stimulus, cells were cultured in non-adherent conditions and deposited on a slide by cytospin for immunostaining. The results show an increase of entotic cells after matrix detachment in all liver cancer cells, even if the percentage of entotic cells remains lower in liver cancer cells (6-14%) when compared to MCF-7 cells (**Figure 1C**). As entosis is favored under stress conditions, we attempted to address whether hypoxia or drug treatment may trigger this mechanism in liver cancer cells. Hypoxic conditions were monitored by the increase in the mRNA expression of *GLUT1* and *VEGF*, two target genes of Hypoxia-Inducible Factor (Hif1α) *(21)* (**Supplemental Figure 1A**). The quantification of entotic Hep3B and Huh7 cells under hypoxia did not show any significant change compared to basal levels (**Supplemental Figure 1B**). We also evaluated whether HCC treatments like Sorafenib could be an inducer of entosis. While Sorafenib treatment altered pERK/ERK ratio as expected, no significant difference was observed in the percentage of entosis in treated versus untreated HCC cells (**Supplemental Figure 1C)**. Altogether, our results demonstrate that liver cancer cells are prone to perform entosis upon proper stimuli such as nutrient deprivation or matrix detachment.

**Figure 1:**
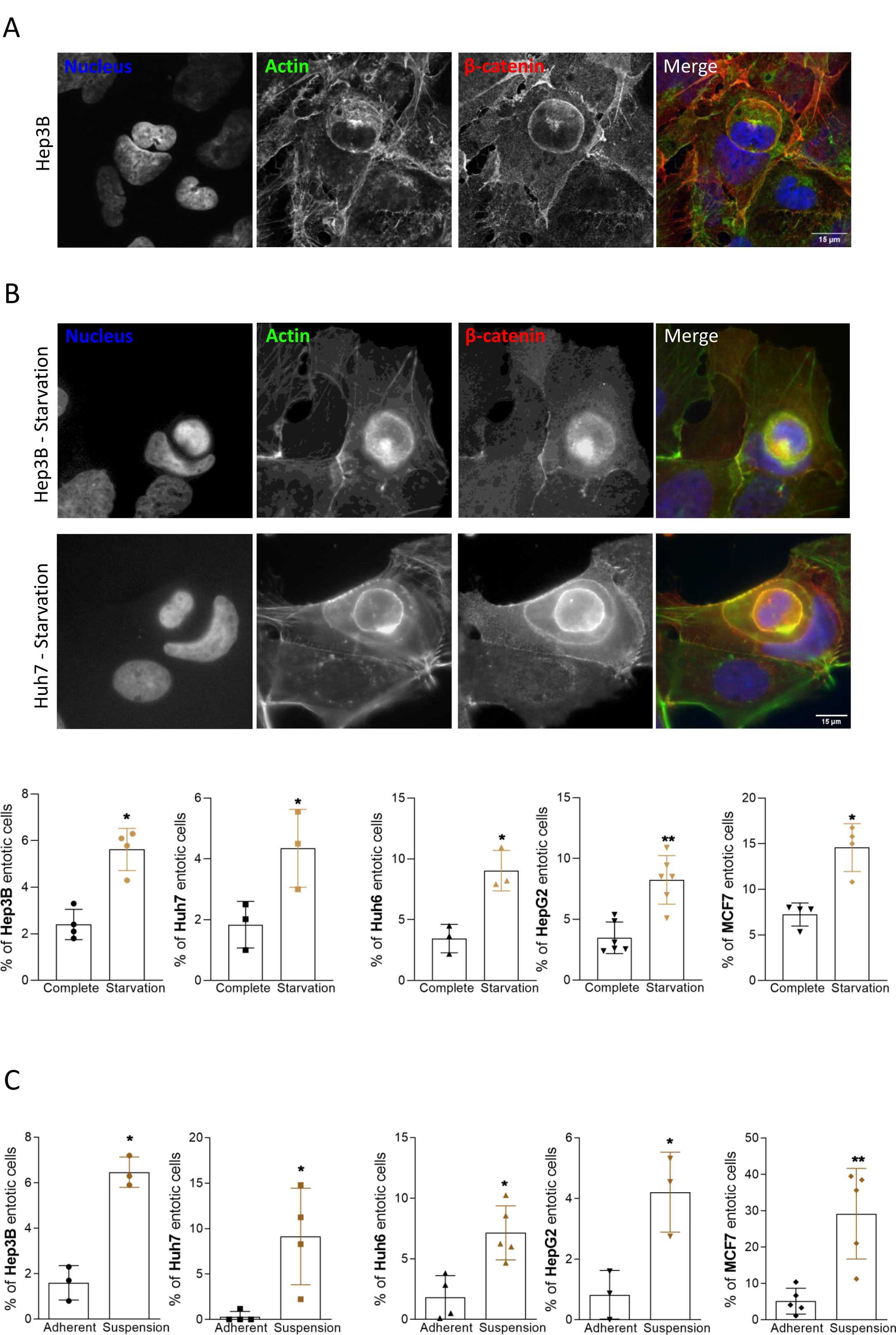
Liver cancer cells perform entosis upon starvation and matrix detachment. **(A)** Representative Hep3B entotic cells characterized using confocal microscopy by the nucleus deformation showed with the DAPI staining (blue) and high concentration of actin with the phalloidin (green). Beta-catenin staining (red) was used to confirm that the inner cell is inside the outer cell. Scale bar, 15µm. **(B)** Nutrient starvation induces entosis as determined by immunofluorescence in Hep3B and Huh7 cells stained for nucleus (blue), actin (green) and beta-catenin (red). Scale bar, 15µm. Graphs show the quantification of entosis induced by starvation in Hep3B, Huh7, Huh6 and HepG2 cells. **(C)** Matrix detachment favors entosis in liver cancer cells. Hep3B, Huh7, Huh6 and HepG2 cells were cultured in non-adherent conditions and analyzed for their ability to form entosis. Graphs show the quantification of entosis induced by suspension culture conditions. (**B-C**) MCF-7 breast cancer cells were used as positive control. Error bars: SD of three or more independent experiments. Significance was determined with the Mann Whitney U test.

### CIC formation is driven by loss of Rnd3 expression

While working on the role of Rnd3 in HCC progression *(18, 20)*, we noticed that, upon Rnd3 silencing, some cells appeared to be inside other cells, reminiscent of entosis. To examine whether the silencing of Rnd3 may trigger this mechanism, we first analyzed Rnd3 expression in liver cancer and MCF7 cells. While Hep3B and Huh7 cells strongly express Rnd3 and Huh6 cells slightly less, Rnd3 is not expressed in HepG2 or MCF-7 cells (**Supplemental Figure 2A**). We thus choose to silence Rnd3 in Hep3B and Huh7 cells using two different approaches: transient transfection of siRNAs using three different siRNAs targeting Rnd3 (SiRnd3#1, #2, #3), or inducible Rnd3 knockdown (KD) in cell lines stably expressing shRNA *(20)*. We found that Rnd3 KD led to a significant increase of CIC events in Hep3B and Huh7 cells regardless of the approach used (**Figure 2A**). Neighboring cells engulfed each other, with about 10% of adherent cells containing engulfed neighbors. CIC events induced by loss of Rnd3 show similar characteristics to stress-induced entotic cells, such as crescent shaped-nuclei and round engulfed cells (**Figure 2B**). More complex cell structures were observed, with three or more cells involved in sequential engulfments (**Figure 2C**). Thus, Rnd3 loss-mediated CIC events share common features with entosis observed in nutrient-depleted or matrix-detached conditions. Moreover, the combination of stimuli, i.e. silencing of Rnd3 and starvation did not increase the entosis mechanism in Hep3B and Huh7 cells, suggesting the involvement of the same pathways to mediate cell engulfment (**Supplemental Figure 2B-C**). The loss of Rnd3 expression, and therefore function, appears to be a trigger of cell engulfment in HCC cells. We next aimed at identifying entosis *in vivo* in tumor tissues using a xenograft mouse model described previously *(20)*. We noticed a significant increase of entosis percentage in tumors generated from Rnd3-KD cells when compared to control Hep3B cells (**Figure 2D**). Thus, Rnd3 loss favors HCC CIC events both *in vitro* and *in vivo*.

**Figure 2:**
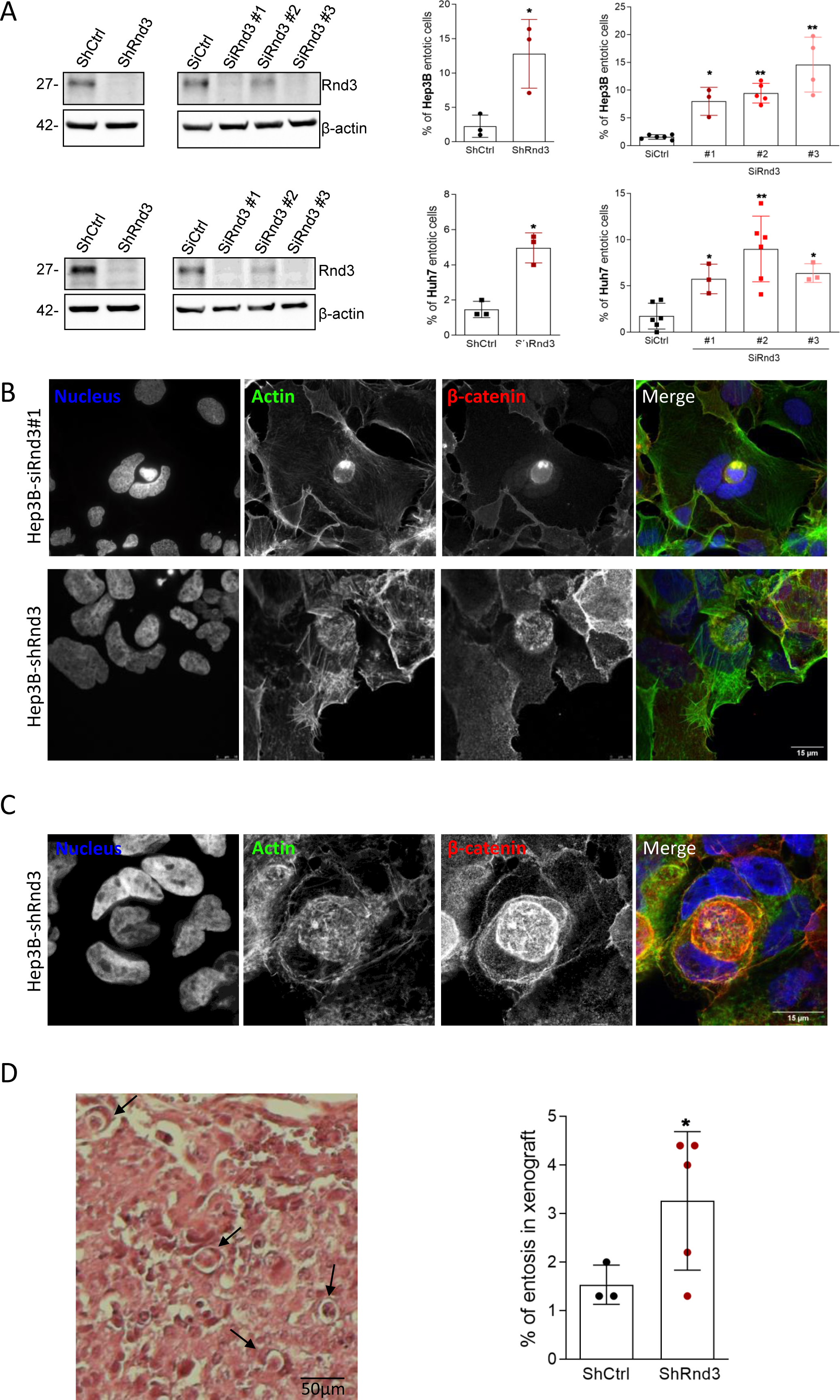
Rnd3 silencing promotes entosis in hepatocellular carcinoma cells. **(A)** The inhibition of Rnd3 expression was performed in Hep3B **(upper panel)** and in Huh7 **(lower panel)** cells using either a shRNA targeting Rnd3 (ShRnd3) inducible with doxycycline treatment and compared to a control shRNA (ShCtrl) or three independent siRNA targeting Rnd3 SiRnd3#1, SiRnd3#2, SiRnd3#3 and SiCtrl as control. Rnd3 knock-down was assessed by Western blot, β-actin is the loading control. Graphs show quantification of entotic cells upon Rnd3 KD. Error bars: SD of three or more independent experiments. Significance was determined with the Mann Whitney U test. **(B-C)** Representative of a single (B) or a double (C) event of entosis after silencing of Rnd3 expression in Hep3B cells with siRnd3 (B, **upper panel**) or with shRnd3 (B, **lower panel; C**). Cells were labelled with DAPI, phalloidin and beta-catenin antibodies to visualize nucleus, actin and membrane respectively. Scale bar, 15µm. **(D)** Entosis was visualized in fixed tumor tissues from mice subcutaneously inoculated with Hep3B-shCtrl (n=3) or Hep3B-shRnd3 (n=5) cells. Tumor sections were stained with hematoxylin-eosin (HE) and the percentage of entotic cells was quantified under the microscope. Note the numerous entotic nuclei, where one nucleus is within the confines of a single cell (arrow). Error bars: SD of three or more independent samples. Significance was determined with the Mann Whitney U test.

### Characteristics of entosis stages after silencing of Rnd3

To investigate the CIC events induced by loss of Rnd3, Hep3B cells expressing GFP-H2B and LifeAct-mRuby were treated with siRnd3 #1 and analyzed by time-lapse microscopy (**Video1**). Entosis initiated by the contact between the two cells, and then the internalization was marked by a high concentration of actin at the border between the two cells. The nucleus of the outer cell acquired a change in its shape due to the nucleus of the inner cell that pushes and deforms it. The final stage was characterized by the degradation of the inner cell, visible by the disappearance of the inner cell nuclei (**Figure 3A**). Using correlative light-scanning electron microscopy (CLEM) allowing analysis of the same entotic cell by fluorescence microscopy and scanning EM (**Figure 3B**), we found that the internalized cell has a rounded shape and seemed to be denser than the outer cell, with apparent cytoskeletal filaments. The inner cell appeared to be inside a large vacuole, the so-called entotic vacuole. The nucleus of the host cell acquired a crescent-like shape and was pushed to cell periphery. The analysis of CLEM demonstrated that CIC structures-mediated by the loss of Rnd3 are similar to entosis, showing a complete internalization of one cell inside another.

**Figure 3:**
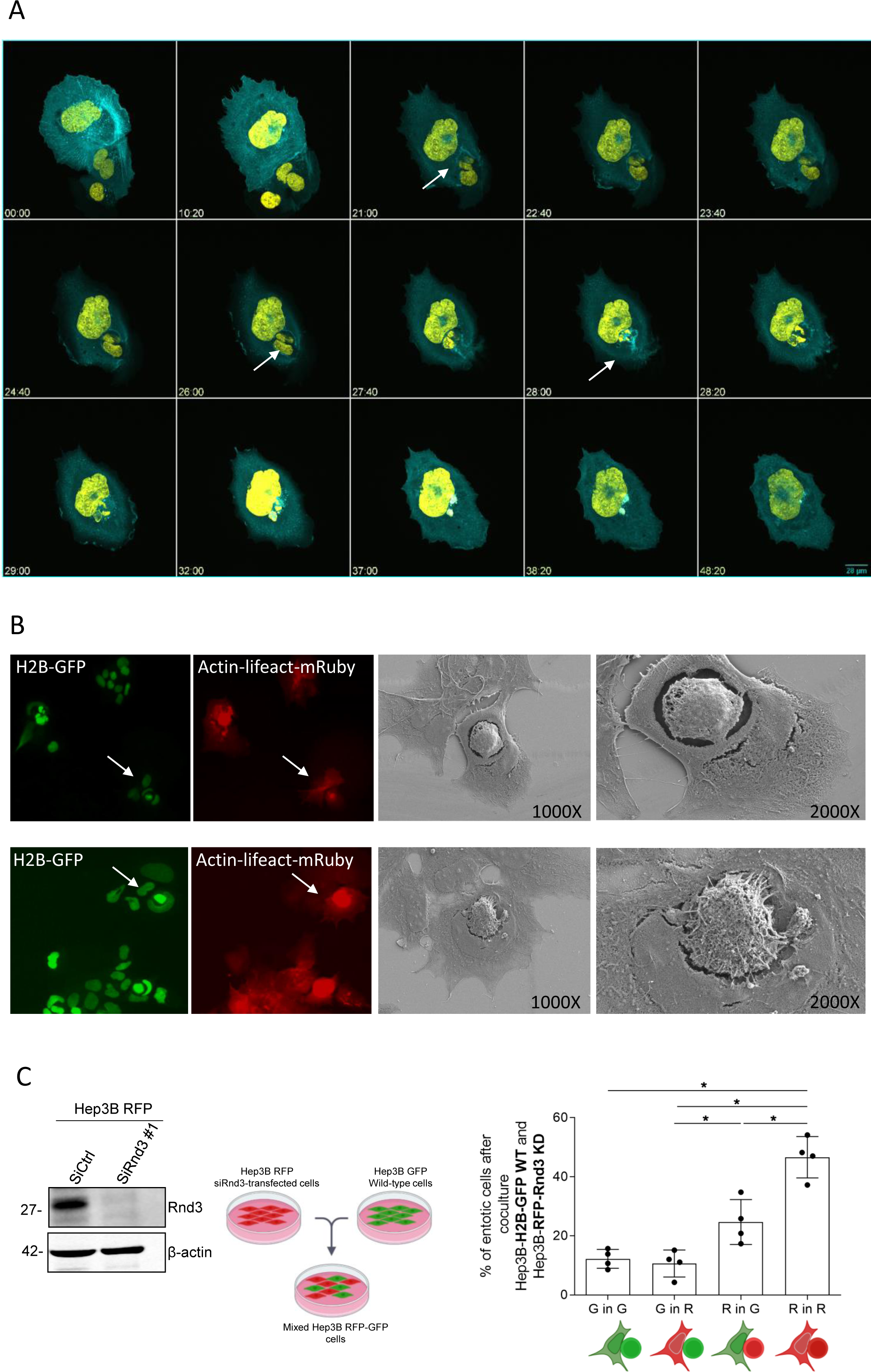
Characterization of entosis mediated by the loss of Rnd3 expression. **(A)** Dynamics of entosis upon Rnd3 silencing. Hep3B cell lines were transduced with H2B-GFP (yellow) and LifeAct-mRuby (cyan) to mark the nucleus and the actin respectively. Cells were transfected with siRNA targeting Rnd3 and spinning disk microscopy analysis was done over 48 hours. The gallery corresponds to Video 1. Time is in hours. Three stages were defined: i) cell contact, ii) internalization, arrowheads show the internalization of the inner binuclear cell 21 hours after contact between cells and iii) degradation of the inner cell; the inner cell degradation starts about 2 hours after internalization and the cell degradation takes almost 10 hours to be completed. Scale bar, 28 µm. **(B)** Analysis of entosis by correlative light-scanning electron microscopy. Entotic cells characterized by the nucleus (H2B-GFP, green) deformation and high concentration of actin (LifeAct-mRuby, red) in the inner cell were chosen using immunofluorescence microscopy and then the same event was analyzed by scanning electron microscopy. Entotic cells were taken with two magnification 1000X and 2000X. **(C)** Two populations of Hep3B cells were mixed, one population transduced with H2B-GFP and another one transduced with H2B-RFP and transfected with siRNA targeting Rnd3. The inhibition was confirmed by Western Blot. 48 hours after mixing, the quantification of entotic cells was performed by evaluating the percentage of different combinations, **G**reen in **G**reen (wild-type in wild-type cells), **G**reen in **R**ed (Wild-type in siRnd3-transfected cells), **R**ed in **G**reen (siRnd3-transfected in wild-type) and **R**ed in **R**ed (siRnd3-transfected in siRnd3-transfected cells). Error bars: SD of three or more independent experiments. Significance was determined with the Mann Whitney U test.

To further examine whether the loss of Rnd3 was required in inner cells, outer cells or both. For that, we generated Hep3B cells with green and red nuclei by expression of GFP-H2B (G) and mCherry-H2B (R), respectively (**Supplemental Figure 3A**). Hep3B-mcherry-H2B were then transfected with siRnd3#1 and co-cultured with untreated Hep3B-GFP-H2B. 48 hours after mixing, entotic cells were analyzed by fluorescence microscopy and counted for each combination (G in G, G in R, R in G, or R in R) (**Figure 3C, Supplemental Figure 3B**). Among all entotic cells, the percentage of events between wild-type cells (G in G) is about 10%, and it reaches 50% between those transfected with siRnd3#1 (R in R). This result demonstrated that the decrease of Rnd3 expression is important in both the inner and outer cells for internalization. We noted that the percentage of the combination R in G is higher than that G in R combination suggesting that the loss of Rnd3 in the inner cell could be sufficient to induce its internalization in the wild-type outer cell.

### Entosis mediated by loss of Rnd3 is dependent on an active Rho/ROCK pathway and a decrease of E-cadherin expression

Rho/ROCK pathway was implicated in the entosis induced after matrix detachment and starvation. As Rnd3 is a known antagonist of this signaling pathway, we analyzed whether the RhoA/ROCK pathway was required for cell engulfment after Rnd3 silencing. We first combined RhoA and Rnd3 knockdowns in HCC cells to assess RhoA involvement (**Supplemental Figure 4A**). We demonstrated that the inhibition of RhoA reverses the effect of Rnd3 silencing by decreasing the percentage of entosis in Hep3B cells (**Figure 4A, left panel**). We next used the Y-27632 compound to inhibit RhoA downstream effector, ROCK. We confirmed that Y-27632 treatment was able to inhibit MYPT1 phosphorylation induced upon Rnd3 KD in Hep3B cells (**Supplemental Figure 4B)**. Moreover, similar to RhoA KD, we found that this treatment decreases the percentage of entosis induced by the silencing of Rnd3 (**Figure 4B, right panel**). Like Rnd3, p190RhoGAP (p190A) is a negative regulator of the Rho/ROCK pathway. HCC cells were transfected with siRNA targeting p190A (**Supplemental Figure 4C**). Consistently the silencing of p190A increased the percentage of entotic cells in Hep3B and Huh7 cell lines, similar results to those obtained after Rnd3 inhibition (**Figure 4B**). The inhibition of p190A expression in Hep3B and Huh7 cells in the presence of siRNA targeting Rnd3 did not modify the percentage of entotic cells compared to the condition where Rnd3 expression was inhibited Rnd3, confirming that Rnd3 and p190A act in the same pathway for the induction of entosis (**Supplemental Figure 5**). Altogether, these data demonstrate that entosis mediated by the silencing of Rnd3 is dependent on the RhoA/ROCK pathway in HCC cells. Therefore, deregulation of the Rho/ROCK pathway alters the ability of HCC cells to perform entosis.

**Figure 4:**
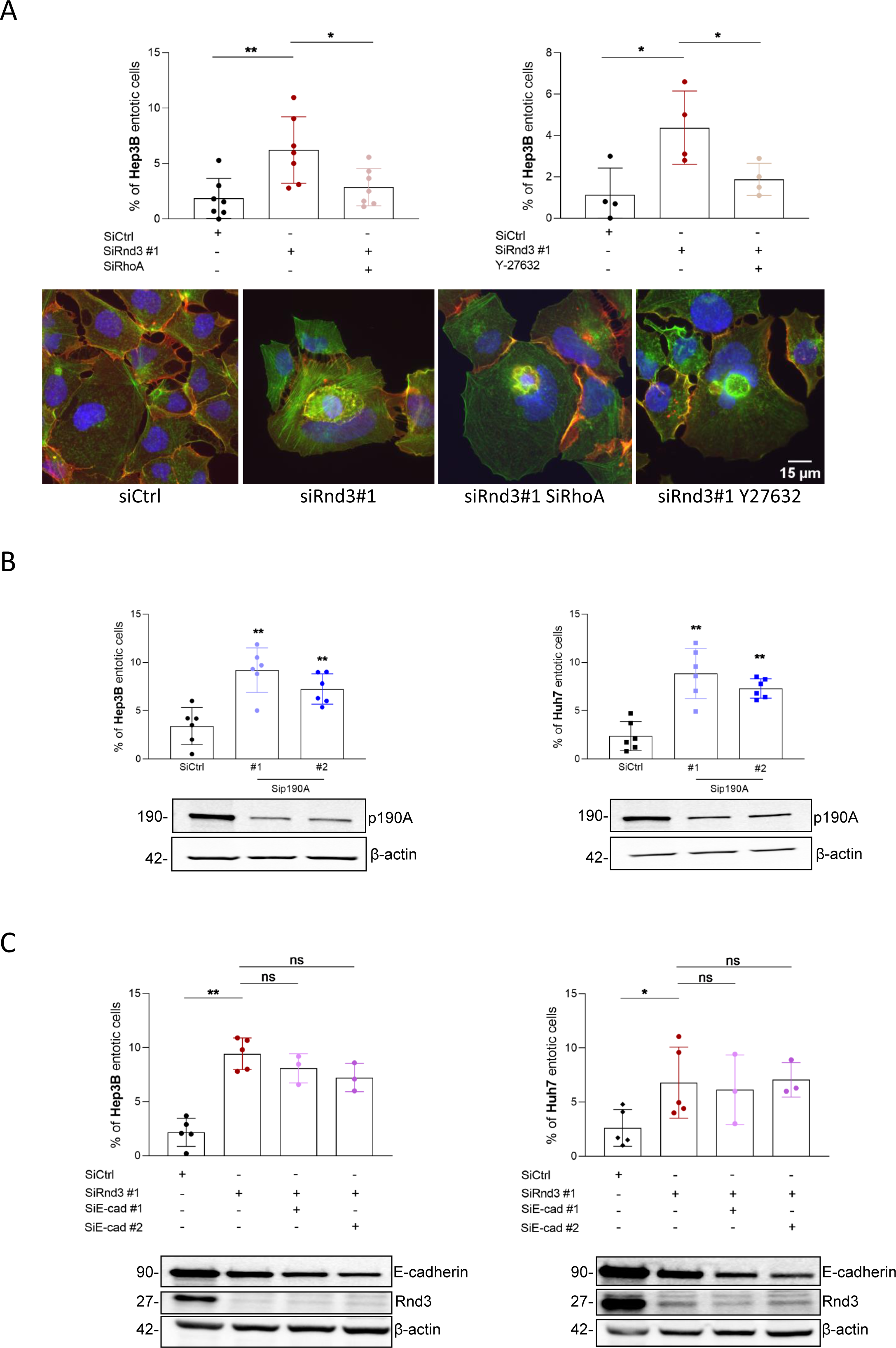
Entosis induced by Rnd3 silencing is dependent on Rho/ROCK pathway and occurs independently of E-cadherin expression. **(A)** Hep3B cells were transfected with siRNA targeting Rnd3 (siRnd3#1) in the presence or absence of siRNA targeting RhoA (siRhoA) (**left panel**), or after treatment or not with Y-27632 (**right panel**) and the percentage of entosis was evaluated. Immunofluorescence images for the different conditions SiCtrl, SiRnd3#1, SiRnd3#1+SiRhoA, SiRnd3#1+Y27632 showing entotic cells stained with DAPI (blue), Phalloidin (green), beta-catenin (red) to visualize nucleus, actin and membrane respectively. **(B)** Hep3B or Huh7 cell lines were treated with siRNA targeting p190RhoGAP-A (siP190A #1 and siP190A#2). The Western blots show the inhibition of p190A and the graphs indicate the percentage of entotic cells. **(C)** Hep3B or Huh7 cells were transfected with siRNA targeting Rnd3 in the presence or absence of siRNA targeting E-cadherin (siEcad#1 or siEcad#2). The inhibition of Rnd3 and E-cadherin was confirmed by Western blot and the percentage of entotic cells was evaluated. Error bars: SD of three or more independent experiments. Significance was determined with the Mann Whitney U test.

Entosis was described to be dependent on E-cadherin-based cell-cell junctions *(15)*. However, we have previously found a decrease in E-cadherin expression upon silencing of Rnd3 in HCC cells *(18)*, which raises the question of the involvement of E-cadherin in the entosis observed here. We thus used E-cadherin-targeting siRNAs and analyzed their impact on entosis induced by the silencing of Rnd3 in HCC cells (**Figure 4C, Supplemental Figure 6A**). As previously published, E-cadherin expression decreased upon Rnd3 silencing in both cell types (**Figure 4C**). Our results showed that inhibition of E-cadherin in Hep3B and Huh7 cells did not alter the percentage of entotic cells compared to Rnd3 knockdown alone (**Figure 4C**), demonstrating that entosis mediated by Rnd3 loss is independent on E-cadherin in HCC cells. These data prompted us to explore the impact of E-cadherin silencing alone on entosis in HCC cells. Surprisingly, in contrast to breast cancer cells *(22)*, we found that decreased E-cadherin expression promoted entosis in HCC cells (**Supplemental Figure 6B**).

### Identification of LAMP1 as an effector of Rnd3 loss-mediated entosis

In order to better characterize the entosis mediated by Rnd3 loss, we sought to apply a global approach. However, isolation of entotic cells is not an easy task, and we failed to do so using flow cytometry. We therefore applied a pipeline combining isolation of entotic cells by laser microdissection in a Hep3B-H2B-GFP cell population transfected with siRnd3#1 (**Figure 5A, left panel**) and mass spectrometry based proteomic analysis according to our published protocol *(23)*. We obtained 139 proteins that were underrepresented in the entotic proteome with an entosis/total proteome abundance ratio ≤ 0.5 and 55 enriched proteins with an entosis/total proteome abundance ratio ≥ 2 (**Supplemental Tables 1-2**). We used Gene Ontology (GO) to classify the identified proteins (**Supplemental Tables 3-4**). We highlighted that the underrepresented proteins were mainly involved in cellular metabolic processes with 63% (n=88) of proteins associated to these biological processes. Proteins associated to intracellular anatomical structures and organelles were also found significantly decreased in entotic samples, suggesting a disassembly of cellular elements. Notably we found a decrease of abundance of Lamin-B1 and Lamin-B2, proteins of the nuclear envelop (**Supplemental Table 2**). In parallel, we found an enrichment of RNA binding proteins (42%, 23 proteins) including ribosomal and/or translation proteins (**Supplemental Table 3**). Network analysis using the Ingenuity Pathways Analysis platform (IPA, Qiagen) of the upregulated proteins further showed an implication of these candidates mainly in cell death/survival (**Figure 5A, Right panel**). We indeed found an overrepresentation of ER-associated proteins involved in stress response and/or folding of proteins such as Prdx2, HSPA5 (Bip) or protein disulfide-isomerases (P4HB, PDIA6) and PPIA, suggesting the recognition of cellular intrusion as a stressful cellular event (**Supplemental Table 1**). Network analysis using IPA also revealed a significantly altered functional network linked to the LAMP1 protein in entotic cells, where proteasome, chaperone and ER stress proteins were also present (**Supplemental Figure 7A**). LAMP1 is known to be implicated in the degradation of the inner cells by autophagy during the entotic mechanism. We confirmed the high expression of LAMP1 in the inner cells in the early on stage of degradation (**Figure 5B**). To confirm the involvement of LAMP1 in entosis induced by Rnd3 silencing, we inhibited Rnd3 in the presence or absence of siRNA targeting LAMP1 in Hep3B and Huh7 cells (**Figure 5C**). Whereas, as previously shown, the silencing of Rnd3 increased the percentage of entosis in both cell lines, this percentage returned to control levels when Rnd3 and LAMP1 were co-silenced (**Figure 5D**). These data demonstrated the involvement of LAMP1 not only in the final stage of inner cell degradation but also in the overall mechanism of entosis induced by silencing of Rnd3. Interestingly, using RNAseq analysis of Hep3B cells (data not shown), *LAMP1* gene expression was found upregulated upon Rnd3 silencing (**Supplemental Figure 7B**). These data were confirmed at the protein level in both Hep3B and Huh7 cells (**Figure 5C, Bottom panel**). We further analyzed whether the overexpression of LAMP1 is sufficient to promote entosis. The ectopic expression of LAMP1 in Hep3B cells only slightly increased the percentage of entotic cells, which remains very low compared to that obtained with Rnd3-silencing alone. However, the combination of overexpression of LAMP1 and Rnd3 inhibition significantly increased the entosis percentage compared to Rnd3-silencing alone (**Figure 5E**). All these results suggest that Rnd3 favors entosis through LAMP1 expression and function. Thus, LAMP1 is an important factor to be considered all over the entotic mechanism.

**Figure 5:**
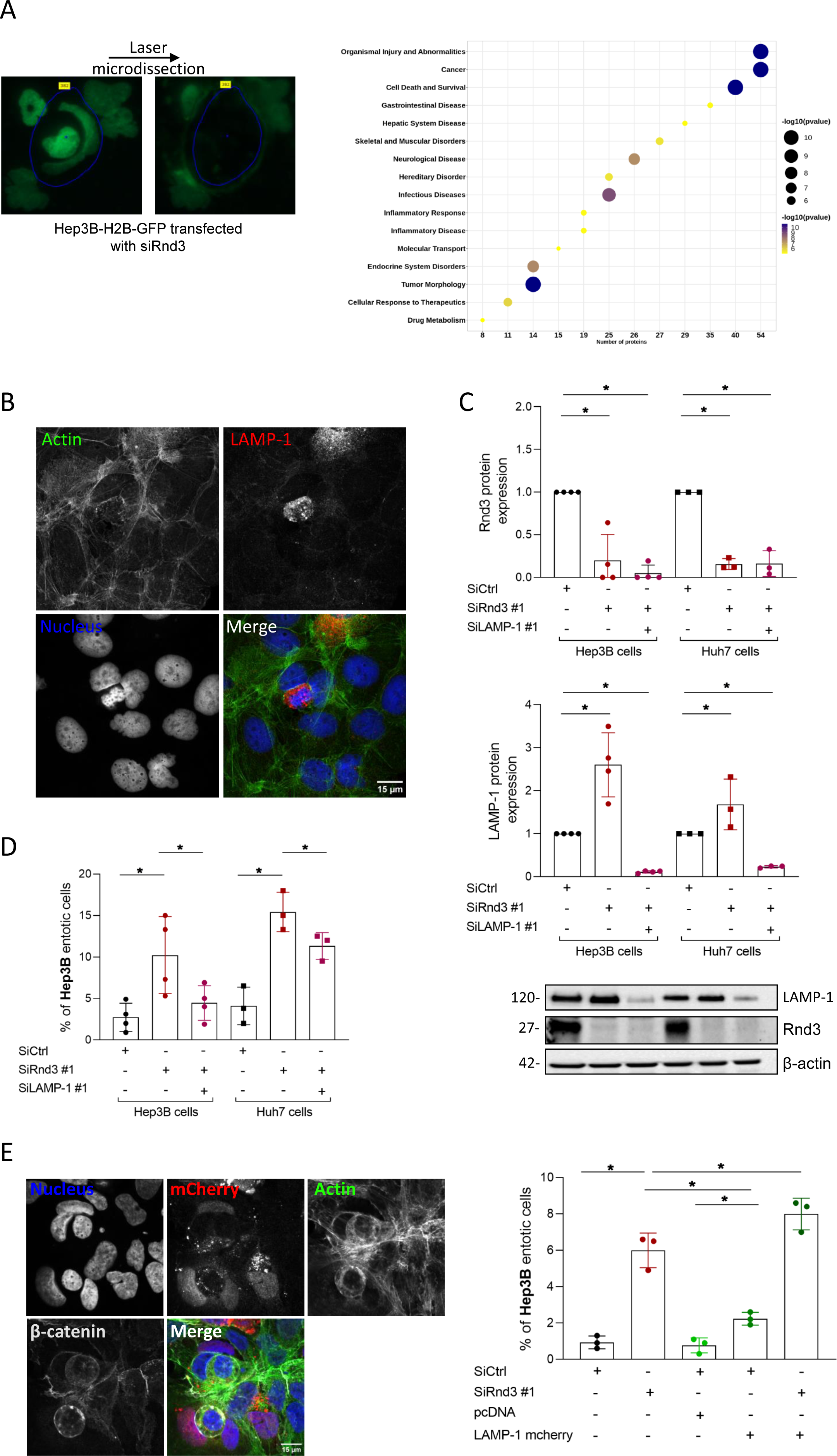
Proteomic analysis and identification of LAMP1 as an effector of Rnd3 loss-mediated entosis. **(A)** Hep3B-H2B-GFP cells were labeled with SILAC and transfected with siRNA targeting Rnd3 (siRnd3#1) and the entotic cells were micro-dissected and analyzed using mass spectrometry. Upregulated proteins when compared to proteins from lysate of Hep3B cells transfected with control siRNA were analyzed using ingenuity pathway analysis (IPA) and represented in a bubble plot showing pathways implicated and the number of proteins. The bubble size represents the number of upregulated proteins involved in a specific pathway. **(B)** Hep3B entotic cells stained to visualize nuclei (Blue), actin (Green) and LAMP1 (Red). **(C)** Hep3B or Huh7 cells were transfected with siRNA targeting Rnd3 in the presence or absence of siRNA targeting LAMP1 (siLAMP1#1). The inhibition was confirmed by Western blot. The graphs show the quantification of Rnd3 and LAMP1 knock-downs in Huh7 and Hep3B cells. (D) Evaluation of entotic cells in Hep3B and Huh7 cell lines after Rnd3 and LAMP1 silencing. Error bars: SD of three or more independent experiments. Significance was determined with the Mann Whitney U test. (E) Overexpression of LAMP1-mcherry in Hep3B cells after silencing of Rnd3. pcDNA is the control vector. Representative image of the overexpression showed by the mcherry staining and the quantification of entotic cells are showed in the right panel of (E).

### Entosis in patient tumors correlates with invasive features

We next investigated the presence of entotic cells in HCC patient tissues. Using membrane (pan-keratin/β-catenin) and nucleus staining, we found few entotic cells in tumor samples (**Figure 6A**). These events were rare, but visible in the tumor part, using immunofluorescence approach on tissues. We then performed Rnd3 staining on HCC sections and looked for entotic cells in negative and positive Rnd3 areas. We noticed that the entotic cells are present in tumor sections with low expression of Rnd3 (**Figure 6B**) supporting our *in vitro* data concerning the implication of Rnd3 downregulation in the entosis process. Using a cohort of 10 patient samples (**Table 1**), we then correlated the number of entotic cells with the characteristics of patient tumors. Similar to the low level of Rnd3 expression *(18)*, we found that the number of entotic cells was significantly higher in tumors with satellite nodules or vascular invasion, which are indicative of local invasion of HCC (**Figure 6C**). All these results suggest the association between the entotic events, loss of Rnd3 and tumor progression.

**Table 1.** Clinical and tumoral characteristics of patients.

| Table 1 Clinical and tumoral characteristics of patients |  |
| --- | --- |
| Clinical characteristics | Value |
| Age (mean ± standard deviation) | 69.5 ± 9.1 years |
| Gender (male) | 100% |
| HBV infection | 0% |
| HCV infection | 44.4% |
| Cirrhosis | 44.4% |
| Tumor characteristics | Value |
| Diameter (mean ± standard deviation) | 56.63 ± 39.76 |
| Satellite nodule | 50% |
| Vascular invasion | 70% |

**Figure 6:**
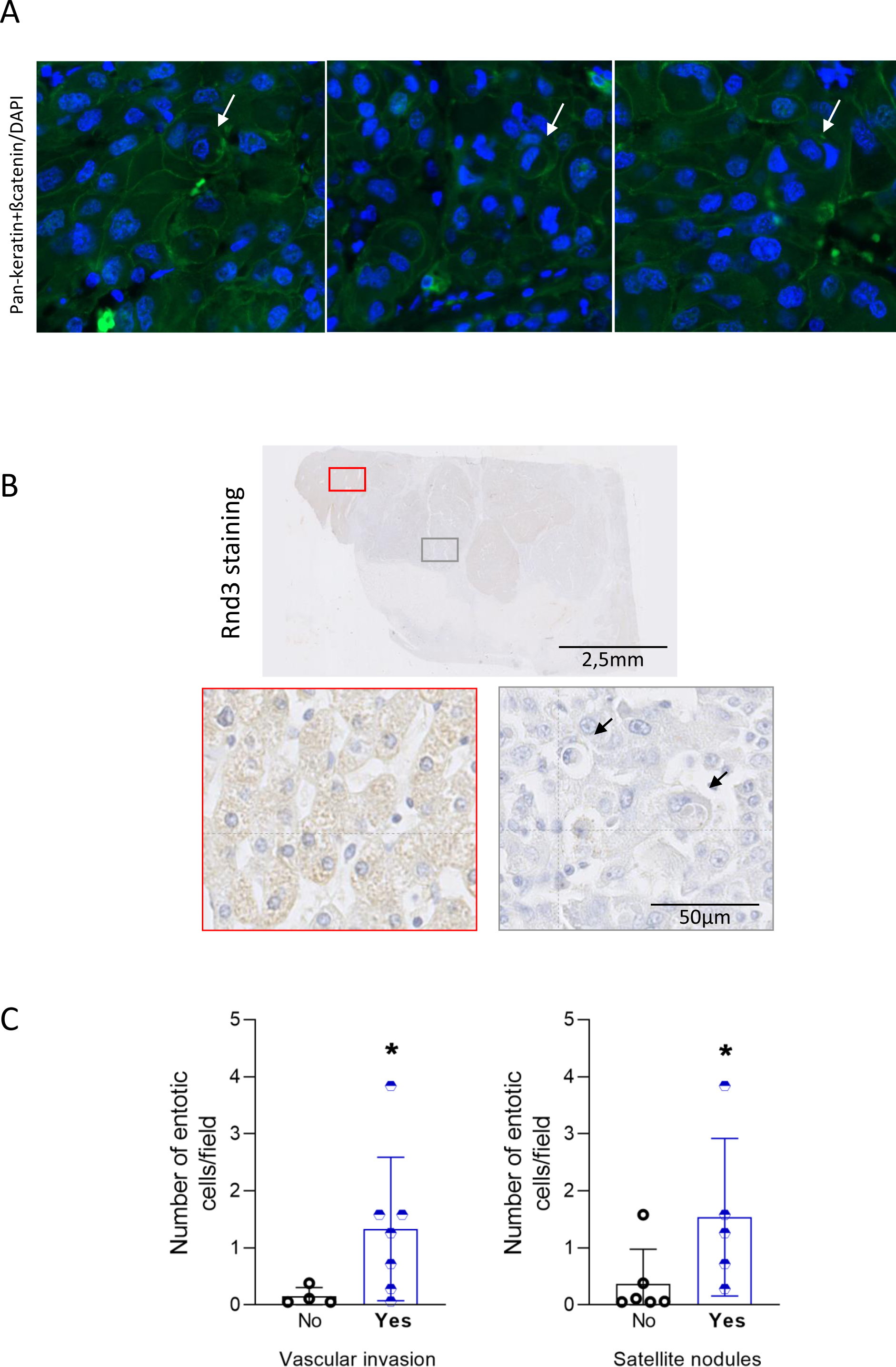
Entosis in patient tumors correlates with invasive features. (A) Representative entotic cells in HCC tumor tissues stained with DAPI (Blue) and pan-keratin/bate-catenin (Green). (B) HCC tissues stained using hematoxylin staining, and Rnd3 antibody. Scale bar: 2.5 mm. Entotic cells were evaluated in the positive (red) or negative (gray) Rnd3 areas. Scale bar: 50µm (C) Correlation between the number of entotic cells and the presence (Yes,) or not (No) of satellite nodules or vascular invasion. N=10 patient samples, Significance was determined with the Mann Whitney U test.

## Discussion

In this study, we characterized entosis in liver cancer cells, which resembles to that largely described in literature in breast cancer cells *(24)*. We demonstrated that entosis could be induced by matrix detachment and nutrient deprivation in HCC cells. In addition, we identified that entosis can be efficiently triggered in HCC cells through the loss of *RND3* expression. Others and we, previously described the down-regulation of this gene expression in human HCC *(18, 19, 25)*. Moreover, Rnd3 loss was associated with liver cancer cell proliferation, invasion, chemoresistance and senescence *(18–20, 26)*. We revealed here that Rnd3 loss also favors entosis in HCC tumors. Recently, Rnd3 has been involved in the p53-dependent entotic mechanism driven by transient mitotic arrest in breast cancer cells *(27)*. However, in contrast to our results, it was found that Rnd3 expression under the regulation of p53 was essential to promote entosis in breast cancer cells. Thus, Rnd3 role appears to be different in breast and liver cancers. In this line, Rnd3 expression does not seems to be regulated by p53 in HCC cells as no correlation between *RND3* expression and TP53 mutations could be established in HCC human samples *(18)*.

Rnd3 is a negative regulator of the RhoA/ROCK pathway. Herein, we confirmed the strong dependency of the entotic mechanism on this pathway. We found that entosis induced by the loss of *RND3* is reduced after the inhibition of Rho/ROCK pathway. Mechanically, entosis depends on actomyosin contractility regulated by the RhoA GTPase activity in the invading cell. It was described that outer cells have lower actomyosin contractility, consistent with a more deformable status compared to invading cells. In our co-culture experiments, the percentage of wild-type cells inside Rnd3-negative cells was very low, suggesting that the penetrating cells require an increase in contractility. However, in liver cancer cells, the loss of Rnd3 is important in both the inner and outer cells, as a maximum of entosis was found when Rnd3 expression is decreased in both the inner and the outer cell. This result is consistent with the existence of a required threshold for Rnd3 expression to favor entosis. Thus, a knockdown recreating an imbalance in contractility, in agreement with the literature, the more rigid cell could be internalized by a more deformable cell *(22)*. Consistently, we observed a high concentration of actin and the presence of fibers in the inner cells by electronic microscopy.

Several studies have shown the importance of E-cadherin in entosis induced by matrix detachment and starvation *(11, 12, 22)*. Interestingly, we showed that entosis induced by the silencing of Rnd3 occurs independently of E-cadherin. Consistent with our published results showing the decrease of E-cadherin expression upon silencing of Rnd3 in HCC cells *(18)*, we even found that loss of E-cadherin favors entosis in Hep3B and Huh7 cells. Thus, entosis induced by the loss of Rnd3 could be dependent on other transmembrane proteins that may trigger cell-cell interaction.

A proteomic analysis of laser-captured entotic cells allowed us to identify an overrepresentation of LAMP1 and ER-associated proteins involved in stress response and/or folding of proteins. Statistical analyses by Gene Set Enrichment Analysis did not reveal significant enrichment of canonical pathways of cellular degradation or stress. However, a functional environment of the LAMP1 protein is significantly modified with key proteins of these pathways. It is possible that entosis pathways involve only certain proteins known to be implicated in degradation and stress, and mobilize a specific protein set. Our data confirm that entotic cells express highly LAMP1, a protein described in literature as implicated in the final stage of entosis i.e. degradation of inner cells. Indeed, the membrane of the entotic vacuole surrounding the internalized cells recruits LAMP1 and LC3 independently of the autophagosome formation. The fusion between the entotic vacuole and the lysosome of outer cells allows to degrade the inner cells *(28)*. Herein, we highlighted that *LAMP1* expression is upregulated at the mRNA and protein levels upon Rnd3 silencing. How Rnd3 mediates *LAMP1* expression is not known and remains to be explored. Our results are consistent with an involvement of LAMP1 in inner cell degradation after internalization induced by the loss of Rnd3. However, we found that LAMP1 is involved not only in degradation but also in the whole mechanism of entosis. LAMP1 could be among the different signals necessary to the internalization. Although LAMP1 primarily resides at the lysosomal membrane, its localization to cell surface expression was described to mediate cell-cell adhesion and favor melanoma cell invasion *(29)*.

The degradation of entotic cells could serve to feed cells, supporting the survival of the outer tumor cells in stress conditions. Entosis may also contribute to tumorigenesis by inducing aneuploidy *(30)*. Others and we previously described Rnd3 as a potential metastasis suppressor as it down-expression was associated to poor prognosis in HCC patients. The results presented here support the hypothesis that loss of Rnd3 could participate to HCC progression though the promotion of entosis. Accordingly, entosis in HCC patient tissues correlates with the presence of satellite nodules and vascular invasion. Thus, targeting the entotic mechanism may be valuable as a novel therapeutic avenue to impair HCC progression.

## Acknowledgments

We thank V. Pitard and N. Dugot-Senant from the FACSility and Histopathology platform respectively (TBMCore, UMS005, Bordeaux). Electron and photonic microscopy was done in the Bordeaux Imaging Center, a service unit of the CNRS-INSERM and Bordeaux University, member of the national infrastructure France Bio Imaging, with the help of Dr Etienne Gontier and Isabelle Svahn. We express our gratitude to Paul Divet, Cyril Dourthe and Clotilde Billottet for their help, with, respectively, evaluation of entotic cells, bubble plot representation of proteomic data and collecting of patient tumor data.

## Author contributions

Study design: SB, VM. Generation of experimental data: SB, LD, CD, SDT, JWD. Analysis and interpretation of data: SB, LD, CD, AAR. Providing biological samples from HCC patients: PBS, ChD. Writing of the manuscript: SB, VM. Critical reading of the manuscript: LD, JWD, AAR, FS. Supervision of the project: VM.

## Materials/Subjects and Methods

### Cell culture

The liver cancer cell lines (Hep3B, Huh7, HepG2, Huh6) and the breast adenocarcinoma cell line MCF7 were cultured in Dulbecco modified Eagle’s medium (DMEM 1x glutamax, Fisher Scientific) supplemented with 10% heat-inactivated fetal bovine serum and incubated at 37°C in a humidified 5% CO2 atmosphere. Cell line authentication was performed using short tandem repeat analysis, and absence of mycoplasma contamination in cell culture media was tested every week. The nutrient deprivation was performed by culturing the cells in DMEM 1x glutamax with very low concentration of glucose (Fisher Scientific) and supplemented with 10% of dialyzed heat inactivated FBS for 10-13h. For induction of cell suspension, the adherent cancer cells were trypsinized and cultured in non-treated plastic Petri dishes for 13h. Stable cell lines with fluorescent nuclei or fluorescent F-actin were generated through transduction with lentiviruses expressing H2B-GFP (Addgene #25999), H2B-RFP (Addgene #26001) or LifeAct-mRuby. Transient knockdowns were done by transfection of small interfering RNAs (siRNAs) into cells using the lipofectamine RNAi max (Invitrogen) according to its manufacturer’s protocol. siRNAs targeting Rnd3 (SiRnd3#1, SiRnd3#2, SiRnd3#3), p190RhoGAP-A (Sip190RhoGAP-A #1, Sip190RhoGAP-A #2), RhoA and E-cadherin (siE-cadherin #1) were purchased from Eurofins Genomics and the sequences are presented in **Supplemental Table 5**. siE-cadherin #2 were purchased from Thermo Fisher Scientific (CDH1, cat#: 4427037, ID: s2768). Control siRNA corresponds to AllStars Negative control from Qiagen. To induce a stable suppression of endogenous Rnd3 expression, we used Hep3B-shRnd3 and Huh7-shRnd3 cell lines with conditional, doxycycline-dependent, expression of a Rnd3 shRNA as described previously *(31)*. Hep3B-shCtrl and Huh7-shCtrl cell lines, conditionally expressing the control shRNA targeting the firefly luciferase, were used as controls. Stable cell lines were cultured as described above and shRNA expressions were induced by a doxycycline treatment (50 ng/mL). For ROCK inhibition, cells were treated with Y-27632 (Sigma Aldrich) at 5 µM for 24 hours. Sorafenib (Bay 43-9006, Enzo Life Science) was used at two doses 8 or 10 µm to treat Huh7 cell lines.

### Antibodies, immunoblot analysis

Cells were lysed in RIPA lysis Buffer (Sigma) supplemented with protease inhibitor cocktail (Roche Diagnostics) and protein concentration was determined using Bio-Rad Protein Assay (Lowry). 60 µg of proteins from each sample was separated on 10% polyacrylamide gel (Bio-Rad) and blotted onto nitrocellulose membranes (0.2 µm nitrocellulose, Bio-Rad) using Trans-Blot Turbo Transfer System (Bio-Rad). The membranes were blocked in Odyssey blocking buffer for 30 minutes and then incubated with each of the following specific primary antibodies: Mouse anti-RhoE/Rnd3 (1:1000, clone 4, Cell Signaling, #3664S); Rabbit anti-actin (1:2000, Sigma Aldrich, #A2066), Mouse anti-HSP90 (1:1000, Santa Cruz, #sc-69703), Rabbit anti-Mypt-1 (1:500, Millipore, #07-672-I); Rabbit anti-phospho-Mypt-1 (Thr696) (1:500, Millipore, #ABS45); Mouse anti-RhoA (1:1000, Santa Cruz, #sc-418); Mouse anti-p190A (1:1000, BD Biosciences, #610149); Mouse anti-E-cadherin (1:1000; BD Biosciences, #610182); Mouse anti-HSP90 (1:1000, Santa Cruz, #sc-69703) at 4°C overnight. All blots were analyzed with the Bio-Rad Chemidoc system.

### Immunofluorescence staining

The following primary antibodies were used for immunofluorescence (IF): Mouse anti-beta-catenin (1:400; BD Biosciences, #610154), Mouse anti-E-cadherin (1:50; BD Biosciences, #610182). IF was performed on cells cultured on glass coverslips after adherence or on suspended cells putted on glass cover by cytospin using the Shandon Cytospin 2 centrifuge at 110 rpm for 5 minutes. Cells were fixed and permeabilized in 4% paraformaldehyde PFA (Electron Microscopy Science, #15710) and 1% Triton X-100 respectively for 10 minutes at RT, followed by three 5 minutes PBS washes and blocking in PBS-5%BSA for 20 minutes. Cells were then incubated with primary antibodies for 45 minutes followed, after PBS washing, by secondary antibodies (Interchim) specific for primary antibodies. F-actin was stained using fluorescent phalloidin (Molecular Probes). Finally, the cells were counterstained with DAPI (Sigma, #D9542) and coverslip mounting on glass slide was done using Fluoromount-G medium (Interchim #FP-483331).

### Time-lapse microscopy

Hep3B cells expressing H2B-GFP and Lifeact-mRuby were cultured on glass-bottom dishes 35 mm high (Ibidi) after Rnd3 silencing, and time-lapse microscopy was performed in 37°C and 5% CO_2_ live-cell incubation chambers. The fluorescence was acquired every 2h for 48 hours using spinning-disk LiveSR confocal microscope. The image analysis and the video reconstitution were done using imageJ program.

### Correlative light-electron microscopy (CLEM)

Hep3B cells expressing H2B-GFP were transfected with siRNA targeting Rnd3 (SiRnd3#1) and cultured on glass coverslips gridded and numbered (Delta microscopy, #72265-12). After selection of the area of interest containing the entotic events using fluorescent microscopy, cells were fixed with 2.5 % glutaraldehyde (Electron Microscopy Science, #15960) for 30 minutes at RT followed by an incubation at 4° for 1 hour in the dark. After washing with sorensen’s phosphate buffer 0.2M, pH 7.4 (Electron Microscopy Science, #11601-10), cells were dehydrated in graded series of ethanol solutions, including 50%, 70%, 90% for 5 minutes and 100% for 2 times, 5 minutes at RT in the dark. Finally, the cells were dried with CO_2_ and observed using Zeiss GeminiSEM300 microscope.

### Entosis quantification

The entotic event is defined by 1) the presence of cell inside another one with a deformed nucleus and 2) the high concentration of actin staining around the inner cell. The entotic cells were observed with epifluorescence microscopy (Zeiss). Quantification was done by counting 300-400 of total cells for each condition. The percentage of entotic events was calculated by dividing the number of entotic events on the total number of cells.

### SILAC labeling, laser capture micro-dissection and proteomic analysis

To discriminate laser captured proteins from undesirable exogenous contaminating proteins, Hep3B-H2B-GFP cells were first metabolically labeled using stable isotope labeling with amino acids in cell culture (SILAC) method, as previously published *(23)*. For that, Hep3B-H2B-GFP cells were cultured in DMEM medium (Dulbecco’s modified Eagle’s medium, Invitrogen) supplemented with 10% dialyzed fetal bovine serum, 200 mg/L L-proline, and 84 mg/L L-Arginine and Lysine. The incorporation of labeled amino acids was done after six cycles of cellular doubling. After transfection with siRnd3 #1, cells were seeded for 12-24 hours on LamPen coated with collagen matrix allowing cells to adhere. Cells were fixed with PFA and the laser capture micro-dissection was performed using PALM type 4 micro-beam (Zeiss). SILAC-labeled cells were transfected with Rnd3-targeting siRNAs and entotic cells with crescent-shaped nuclei were micro-dissected. To obtain enough material for proteomic analysis, we manually collected 2,000 cells in triplicate. With 2,000 isolated entotic cells, we have identified a mean of 2880 ^13^C peptides corresponding to 406 proteins with at least 2 specific peptides. Given the small quantity of material analyzed, the first step guarantees the specificity of identifications by distinguishing labeled proteins from microdissected cells from unlabeled environmental contaminants (keratins, skin and hair proteins, etc…). In parallel, a standard range (500 ng to 5 ng) of protein quantity was done on cells labeled with SILAC and transfected with control siRNA (siCtrl) in order to compare the same quantity of entotic cells micro-dissected and obtained after transfection with siRnd3 to cells transfected with siCtrl. Three independent biological replicates on total protein extracts from SILAC-labeled cells were compared by label-free protein quantification. Proteins were loaded on a 10% acrylamide SDS-PAGE gel and proteins were visualized by Colloidal Blue staining. Migration was stopped when samples had just entered the resolving gel and the unresolved region of the gel was cut into only one segment. The steps of sample preparation and protein digestion by the trypsin were performed as previously described *(32)*. NanoLC-MS/MS analysis were performed using an Ultimate 3000 RSLC Nano-UPHLC system (Thermo Scientific, USA) coupled to a nanospray Orbitrap Fusion™ Lumos™ Tribrid™ Mass Spectrometer (Thermo Fisher Scientific, California, USA). Each peptide extracts were loaded on a 300 µm ID x 5 mm PepMap C_18_ precolumn (Thermo Scientific, USA) at a flow rate of 10 µL/min. After a 3 min desalting step, peptides were separated on a 50 cm EasySpray column (75 µm ID, 2 µm C_18_ beads, 100 Å pore size, ES903, Thermo Fisher Scientific) with a 4-40% linear gradient of solvent B (0.1% formic acid in 80% ACN) in 57 min. The separation flow rate was set at 300 nL/min. The mass spectrometer operated in positive ion mode at a 2.0 kV needle voltage. Data was acquired using Xcalibur 4.4 software in a data-dependent mode. MS scans (m/z 375-1500) were recorded at a resolution of R = 120000 (@ m/z 200), a standard AGC target and an injection time in automatic mode, followed by a top speed duty cycle of up to 3 seconds for MS/MS acquisition. Precursor ions (2 to 7 charge states) were isolated in the quadrupole with a mass window of 1.6 Th and fragmented with HCD@28% normalized collision energy. MS/MS data was acquired in the ion trap with rapid scan mode, a 20% normalized AGC target and a maximum injection time in dynamic mode. Selected precursors were excluded for 60 seconds. Protein identification was done in Proteome Discoverer 2.5. Mascot 2.5 algorithm was used for protein identification in batch mode by searching against a UniProt *Homo sapiens* protein database (75796 entries, release September 3, 2020; https://www.uniprot.org/ website). Two missed enzyme cleavages were allowed for the trypsin. Mass tolerances in MS and MS/MS were set to 10 ppm and 0.6 Da. Oxidation (M), acetylation (K), SILAC modifications (K, R) were searched as dynamic modifications and carbamidomethylation (C) as static modification. Raw LC-MS/MS data were imported in Proline Studio for feature detection, alignment, and quantification. Protein identification was accepted only with at least 2 specific peptides with a pretty rank=1 and with a protein FDR value less than 1.0% calculated using the “decoy” option in Mascot. Label-free quantification of MS1 level by extracted ion chromatograms (XIC) was carried out with parameters indicated previously *(32)*. The normalization was carried out on median of ratios. The inference of missing values was applied with 5% of the background noise. The mass spectrometry proteomics data have been deposited to the ProteomeXchange Consortium via the PRIDE partner repository *(33)* with the dataset identifier PXD043640.

### Immunohistochemistry on HCC samples

Mouse HCC samples were from Hep3B xenografts previously described *(20)*. This model consisted of subcutaneous inoculation of Hep3B-shCtrl or Hep3B-shRnd3 cells in immunodeficient NOG mice treated with doxycycline to induce or not Rnd3 knockdown. The analysis of entosis was performed on tumor sections stained with hematoxylin to mark cytoplasm and nuclei of cells. Human samples came from resected or explanted livers with HCC of patients treated in Bordeaux from 1992-2005. The characteristics of HCCs used for the IHC analysis (10 HCCs) are indicated in Table 1. Formalin-fixed, paraffin-embedded sections of human or mouse HCC samples were used for Hematoxylin-Eosin or immunohistochemical staining. Staining of Rnd3 and pan-keratin/beta-catenin was performed as previously published *(18, 34)*.

### Statistical analysis

The frequency of entotic cells was represented as a percentage (mean±SD) of the total counted cells for at least three experiments. Data were analyzed using GraphPad Prism 10. For all experiments, significance was determined with the Mann Whitney U test, *\*p<0.05, **p<0.01, ***p<0.001*.

## Supplemental information

### Supplemental figure legends

**Supplemental Figure 1:**
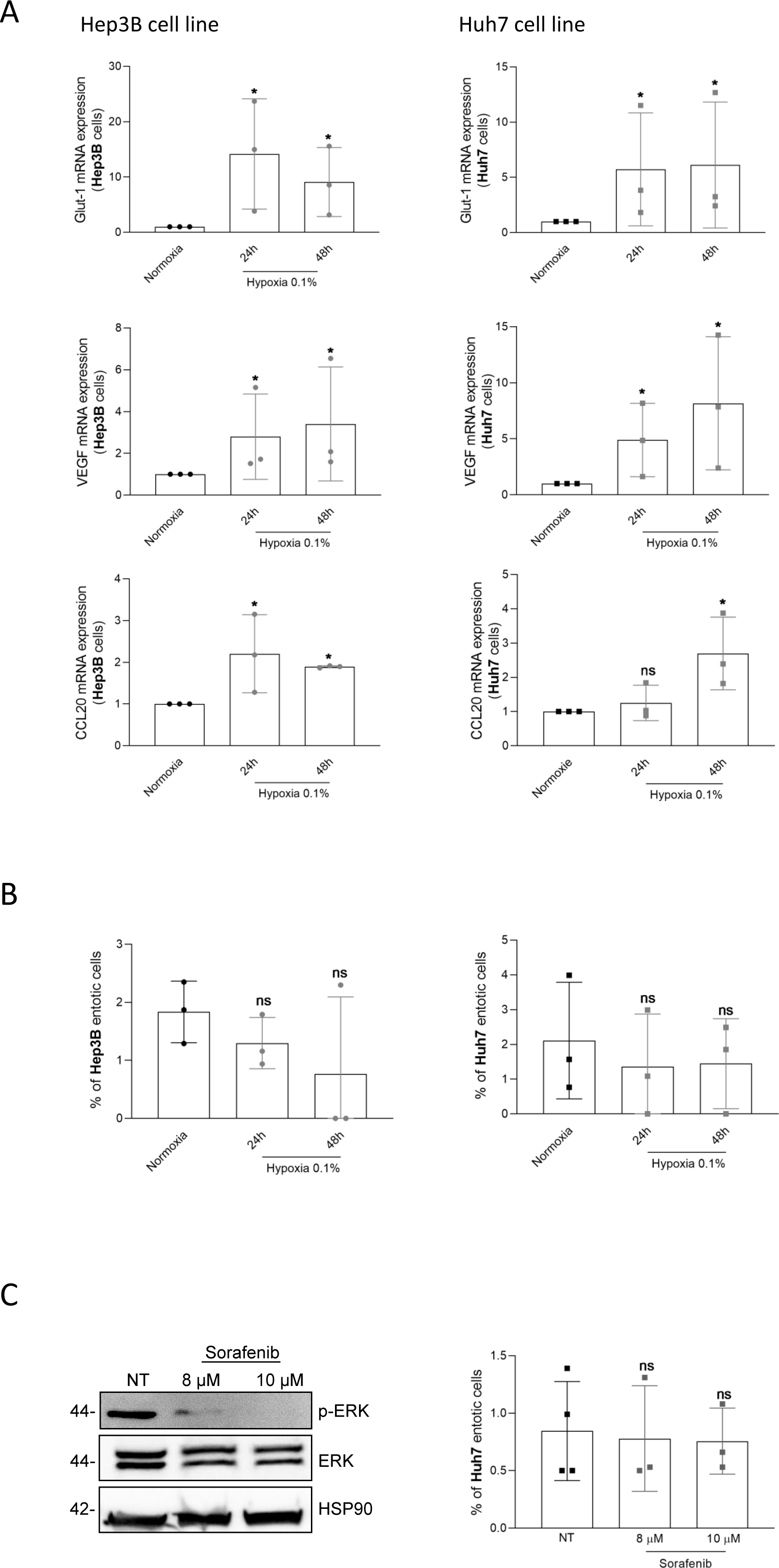
(A) Hypoxia and Sorafenib treatments do not induce entotic events in Hep3B and Huh7 cell lines. **(A)** Quantification of mRNA levels of GLUT1, VEGF, CCL20 in Hep3B (left panel) and Huh7 (right panel) cell lines 24h or 48h after hypoxia (0.1% O2) compared to cells grown in normoxia condition. **(B)** Evaluation of percentage of Hep3B and Huh7 entotic cells after hypoxia. **(C)** Huh7 cell line treated with two doses of Sorafenib (8, 10 µM) for 72 hours and confirmed by Western Blot showing the inhibition of p-ERK compared to the total ERK. The right panel presented the quantification of entotic events in these conditions. Error bars: SD of three independent experiments. Significance was determined with the Mann Whitney U test.

**Supplemental Figure 2:**
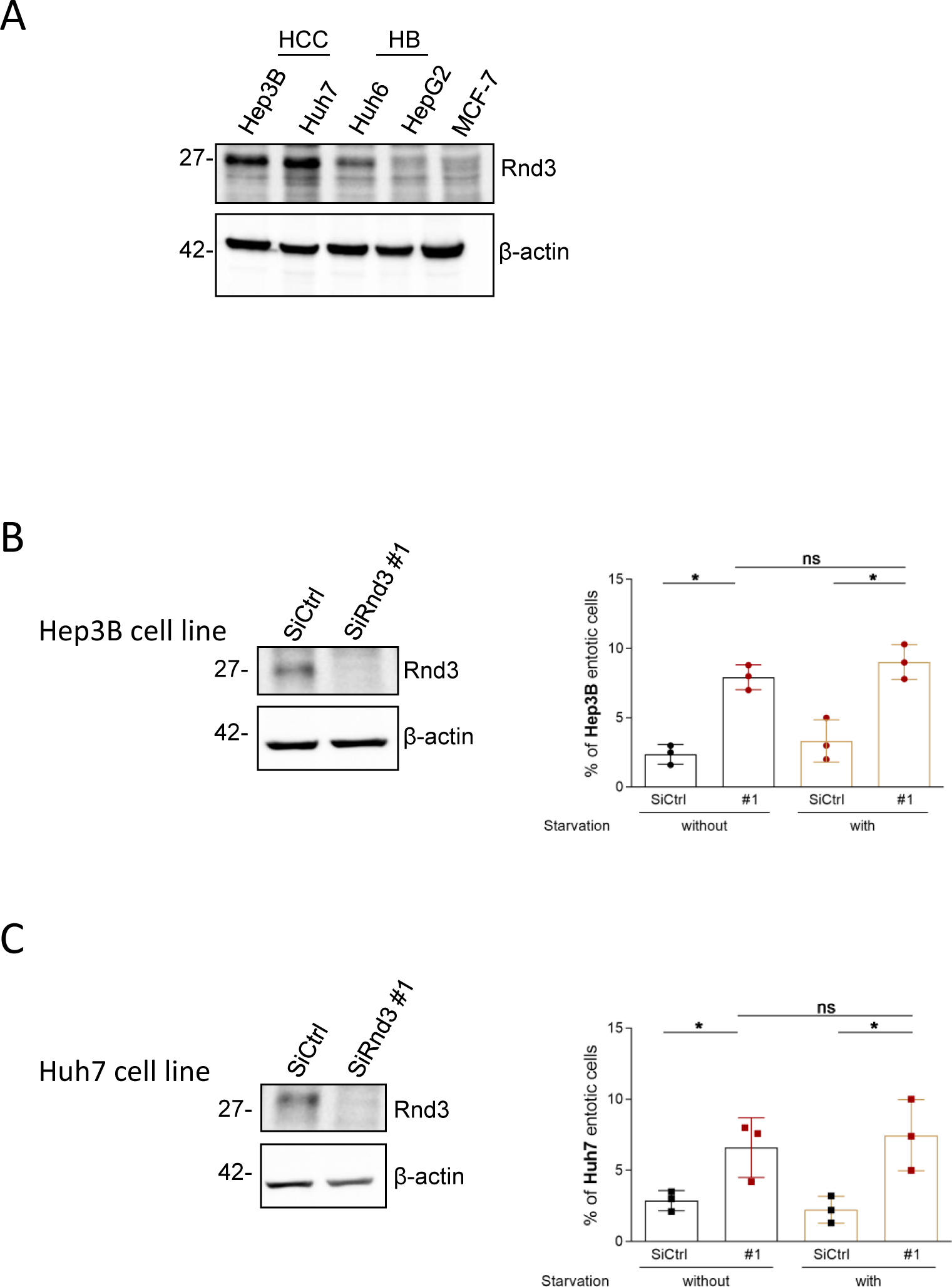
The combination of Rnd3 silencing and starvation does not increase the entosis mechanism in Hep3B and Huh7 cells compared to that induced by Rnd3 silencing alone. **(A)** Comparison of the Rnd3 protein expression between HCC (Hep3B, Huh7), HB (Huh6, HepG2) and MCF7 breast cancer cell lines. **(B-C)** Evaluation of entotic events in Hep3B **(B)** and Huh7 **(C)** after silencing of Rnd3 combined or not with starvation. Error bars: SD of three independent experiments. Significance was determined with the Mann Whitney U test.

**Supplemental Figure 3:**
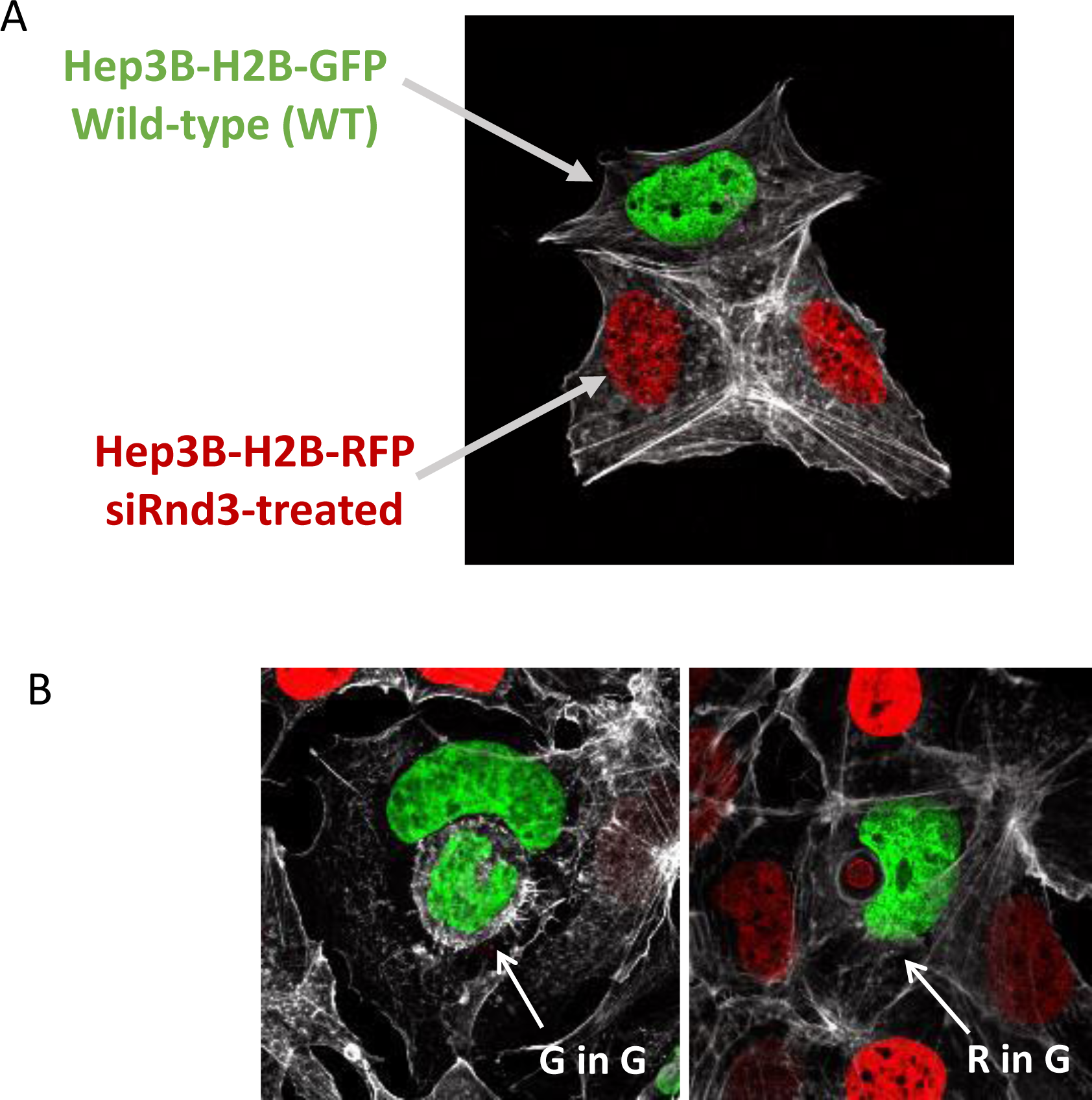
Role of Rnd3 silencing in the inner or outer cells. **(A)** Image of Rnd3-silencing RFP cells and wild-type GFP cells co-culture. (B) Examples of events counted after mixing of Rnd3-silencing RFP cells and wild-type GFP cells; G in G = **G**reen in **G**reen (wild-type in wild-type cell), R in G= **R**ed in **G**reen (siRnd3-transfected in wild-type cell).

**Supplemental Figure 4:**
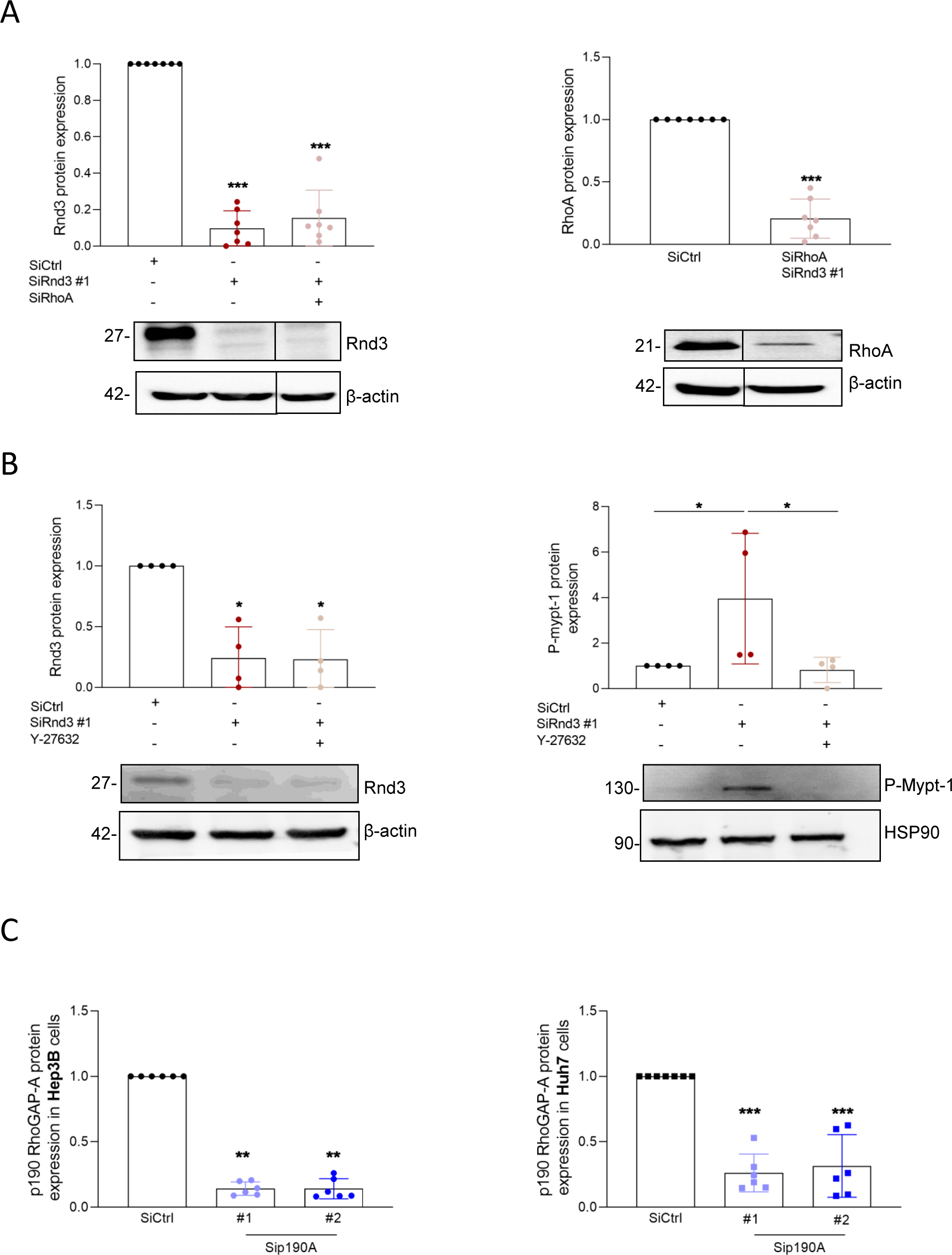
Validation of Rnd3, RhoA, p190RhoGAP-A inhibition by western blot. **(A-B)** Western blot and quantification of Rnd3 and RhoA knock-downs in Hep3B cells in conditions corresponding to Figure 4A. ß-actin was used as loading control (A and B, left-hand graph). Inhibition of ROCK using Y-27632 was validated by the expression of P-Mypt-1. HSP90 was used as loading control (B, right-hand graph). **(C)** Protein expression of p190RhoGAP-A after its silencing in Hep3B (left-hand graph) and Huh7 (right-hand graph) cells, in conditions corresponding to Figure 4B. Error bars: SD of three or more independent experiments. Significance was determined with the Mann Whitney U test.

**Supplemental Figure 5:**
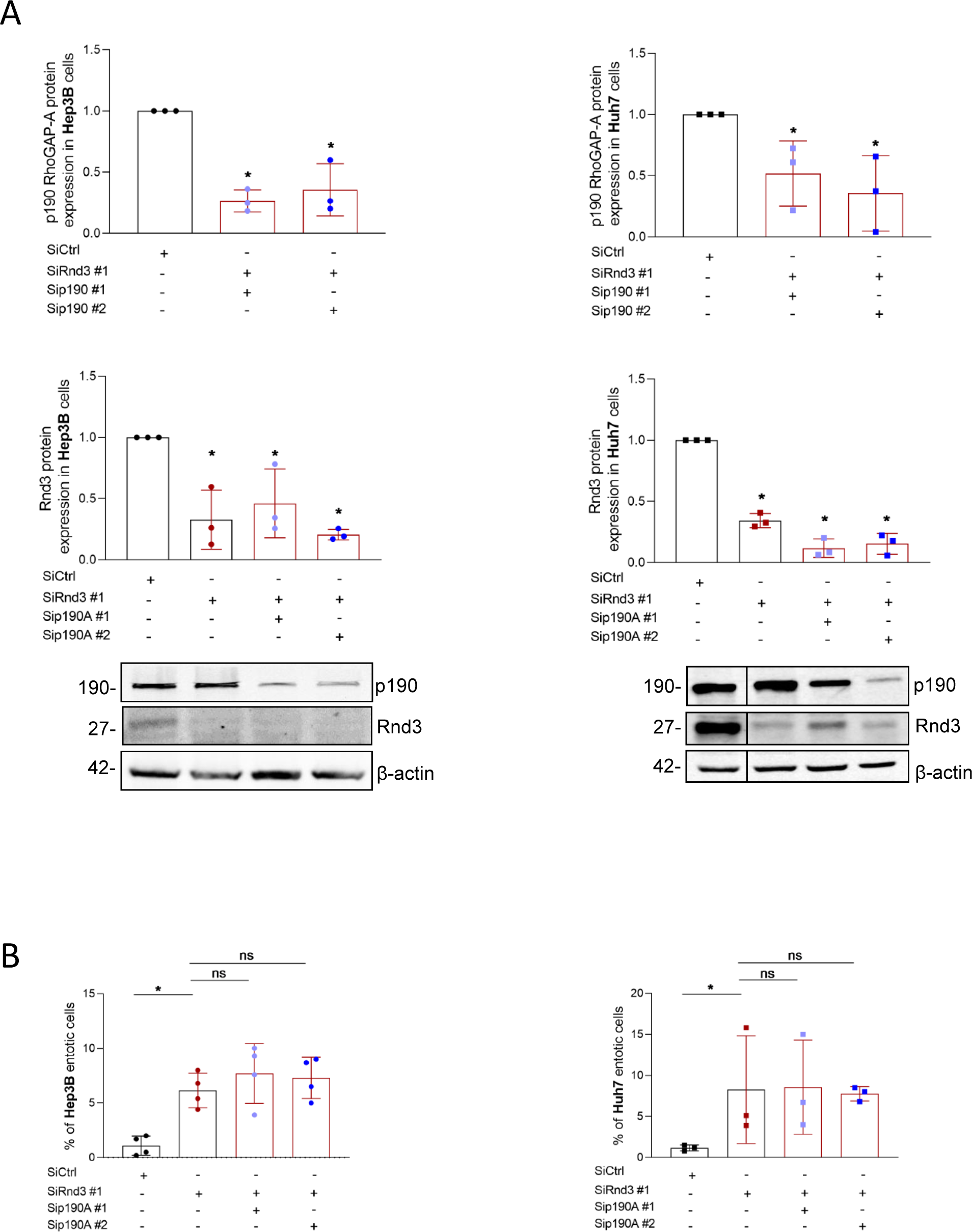
The combination of p190RhoGAP-A and Rnd3 silencing does not increase the entotic events in Hep3B and Huh7 cells. **(A)** Evaluation of the p190RhoGAP-A and Rnd3 protein expression after using siRNA by Western blot in Hep3B (Left panel) and Huh7 (Right panel) cells and the quantification of entotic events are represented in **(B)**. Error bars: SD of three or more independent experiments. Significance was determined with the Mann Whitney U test.

**Supplemental Figure 6:**
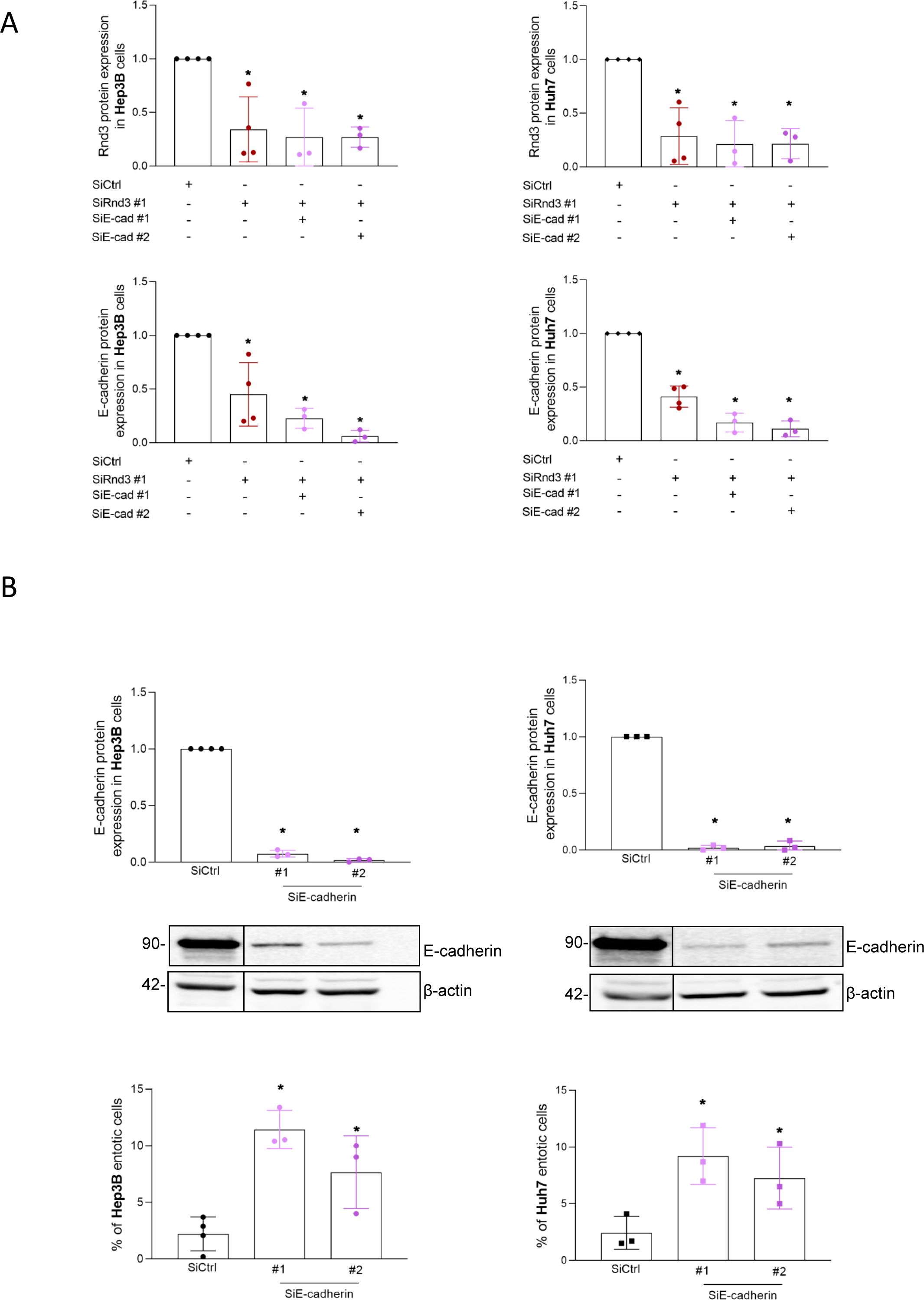
The silencing of E-cadherin alone increases the percentage of entotic events in Hep3B and Huh7 cells. **(A)** Validation of the inhibition of Rnd3 in the presence or absence of two siRNA targeting E-cadherin (#1 and #2) in Hep3B and Huh7 cell lines. **(B)** Evaluation of the entotic cells after inhibition of E-cadherin alone. Error bars: SD of three or more independent experiments. Significance was determined with the Mann Whitney U test.

**Supplemental Figure 7:**
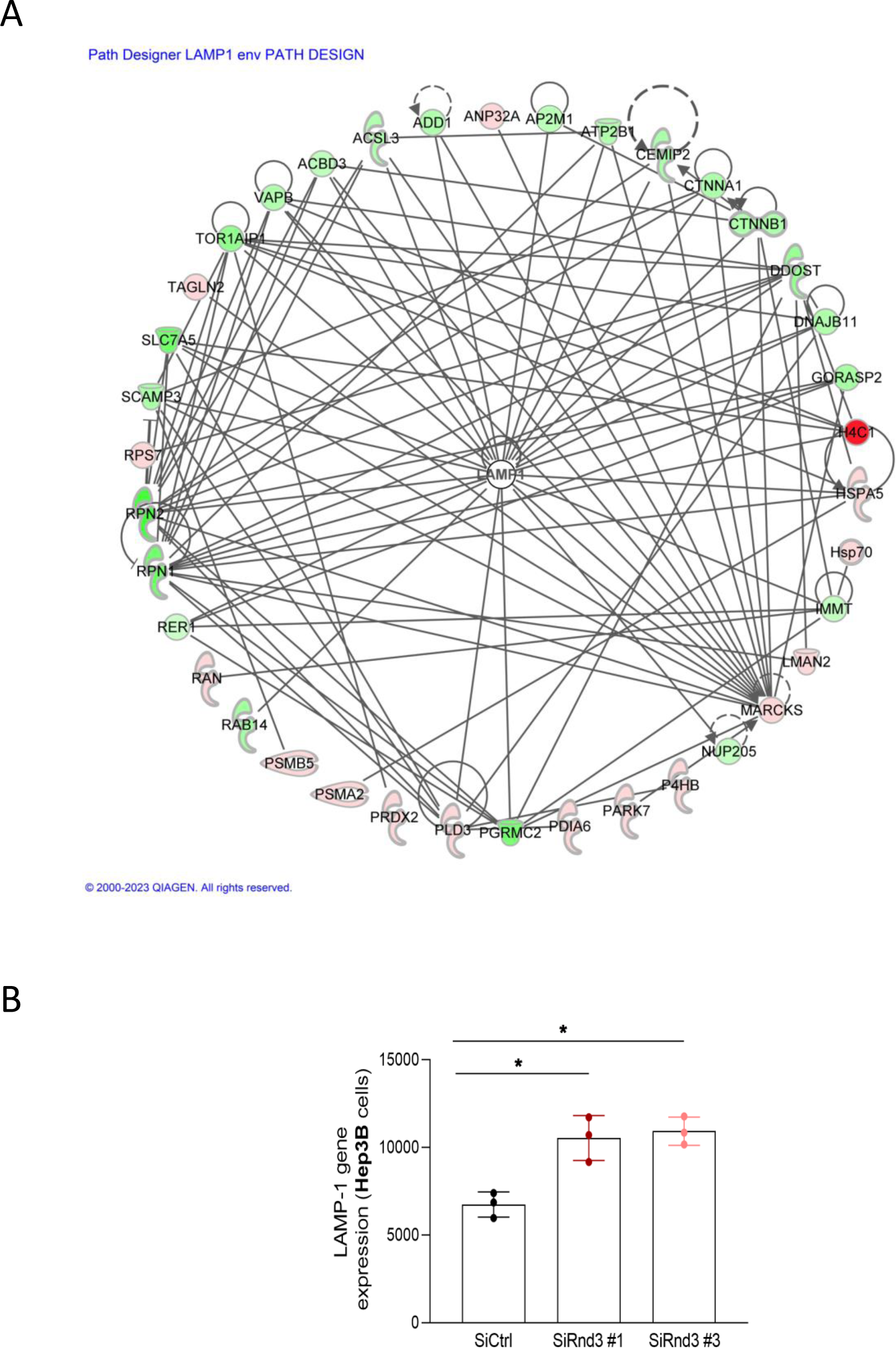
**(A)** Functional interactome of the LAMP1 protein identified in the proteomic profile of entotic cells. Networks produced with the Ingenuity Pathways platform (IPA, QIAGEN). Proteins over-represented or under-represented in entotic cells were colored in red or green, respectively. **(B)** mRNA expression of *LAMP1* upon silencing of Rnd3. Hep3B cells were transfected by siRNA targeting Rnd3 (siRnd3#1 and siRnd3#3) or control siRNA (siCtrl) and transcriptome analysis was performed by RNAseq. The graph represents the normalized expression of the *LAMP1* gene (normalized read counts divided by median of transcripts length in kb). Significance was determined with the Mann Whitney U test.

### Supplemental tables

**Supplemental Table 1:** Results of the mass spectrometry analysis. List of proteins overrepresented in the entotic fraction.

| GN | accession | description | Ratio n1 |
| --- | --- | --- | --- |
| H4C1 | P62805 | Histone H4 OS=Homo sapiens OX=9606 GN=H | 332,16 |
| ARPC4-TTLL3 | F8WCF6 | Actin-related protein 2/3 complex subunit 4 C | 7,18 |
| ACTN1 | P12814 | Alpha-actinin-1 OS=Homo sapiens OX=9606 C | 3,95 |
| RPS19 | P39019 | 40S ribosomal protein S19 OS=Homo sapiens | 29,34 |
| PFN1 | P07737 | Profilin-1 OS=Homo sapiens OX=9606 GN=PF | 25,12 |
| RALY | Q5QPL9 | RNA-binding protein Raly (Fragment) OS=Hor | 12,27 |
| PPIA | P62937 | Peptidyl-prolyl cis-trans isomerase A OS=Hor | 24,05 |
| APRT | H3BQF1 | Adenine phosphoribosyltransferase (Fragmer | 38,12 |
| RPS13 | P62277 | 40S ribosomal protein S13 OS=Homo sapiens | 10,90 |
| CFL1 | P23528 | Cofilin-1 OS=Homo sapiens OX=9606 GN=CFL | 27,69 |
| RPS25 | P62851 | 40S ribosomal protein S25 OS=Homo sapiens | 7,04 |
| SRP14 | P37108 | Signal recognition particle 14 kDa protein OS= | 18,11 |
| RPS11 | P62280 | 40S ribosomal protein S11 OS=Homo sapiens | 9,40 |
| PSMA2 | A0A024RA52 | Proteasome subunit alpha type OS=Homo sa | 14,54 |
| LAMP1 | P11279 | Lysosome-associated membrane glycoprotein | 2,90 |
| PRDX2 | P32119 | Peroxiredoxin-2 OS=Homo sapiens OX=9606 C | 22,87 |
| PLIN2 | Q99541 | Perilipin-2 OS=Homo sapiens OX=9606 GN=Pl | 10,95 |
| GLO1 | Q04760 | Lactoylglutathione lyase OS=Homo sapiens O | 16,59 |
| PSMB6 | P28072 | Proteasome subunit beta type-6 OS=Homo sa | 10,76 |
| TAGLN2 | P37802 | Transgelin-2 OS=Homo sapiens OX=9606 GN= | 21,78 |
| RPS5 | M0R0F0 | 40S ribosomal protein S5 (Fragment) OS=Hon | 12,41 |
| ANP32A | H0YN26 | Acidic leucine-rich nuclear phosphoprotein 3: | 4,97 |
| PSMB5 | P28074 | Proteasome subunit beta type-5 OS=Homo sa | 13,21 |
| TUBB3 | Q13509 | Tubulin beta-3 chain OS=Homo sapiens OX=9 | 13,57 |
| RPLP0 | P05388 | 60S acidic ribosomal protein P0 OS=Homo sa | 7,21 |
| PSMA1 | P25786 | Proteasome subunit alpha type-1 OS=Homo s | 6,64 |
| PSMA6 | G3V5Z7 | Proteasome subunit alpha type OS=Homo sa | 5,05 |
| PDIA6 | Q15084 | Protein disulfide-isomerase A6 OS=Homo sap | 8,20 |
| ILF2 | B4DY09 | Interleukin enhancer-binding factor 2 OS=Ho | 4,01 |
| EIF4B | E7EX17 | Eukaryotic translation initiation factor 4B OS= | 4,42 |
| ALDH1A1 | P00352 | Retinal dehydrogenase 1 OS=Homo sapiens C | 13,98 |
| LMAN2 | Q12907 | Vesicular integral-membrane protein VIP36 C | 3,26 |
| MARCKS | P29966 | Myristoylated alanine-rich C-kinase substrate | 4,50 |
| IDH1 | O75874 | Isocitrate dehydrogenase [NADP] cytoplasmic | 15,00 |
| P4HB | H0Y3Z3 | Protein disulfide-isomerase (Fragment) OS=H | 8,67 |
| QPRT | Q15274 | Nicotinate-nucleotide pyrophosphorylase [ca | 8,26 |
| TUBB | P07437 | Tubulin beta chain OS=Homo sapiens OX=960 | 8,06 |
| CLIC1 | O00299 | Chloride intracellular channel protein 1 OS=H | 10,49 |
| P4HB | P07237 | Protein disulfide-isomerase OS=Homo sapien | 6,19 |
| PLD3 | Q8IV08 | 5'-3' exonuclease PLD3 OS=Homo sapiens OX | 8,44 |
| DECR1 | Q16698 | 2,4-dienoyl-CoA reductase, mitochondrial OS | 2,27 |
| DTYMK | H7C312 | Thymidylate kinase (Fragment) OS=Homo sa | 7,14 |
| TALDO1 | P37837 | Transaldolase OS=Homo sapiens OX=9606 GN | 9,40 |
| IGF2BP1 | Q9NZI8 | Insulin-like growth factor 2 mRNA-binding pr | 3,43 |
| AIMP2 | Q13155 | Aminoacyl tRNA synthase complex-interactin | 3,36 |
| MDH1 | A0A5K1VW95 | Malate dehydrogenase OS=Homo sapiens OX | 10,68 |
| PEBP1 | P30086 | Phosphatidylethanolamine-binding protein 1 | 2,32 |
| RPS7 | P62081 | 40S ribosomal protein S7 OS=Homo sapiens C | 10,63 |
| ANXA4 | P09525 | Annexin A4 OS=Homo sapiens OX=9606 GN= | 8,18 |
| RAN | P62826 | GTP-binding nuclear protein Ran OS=Homo si | 5,59 |
| HSPA5 | P11021 | Endoplasmic reticulum chaperone BiP OS=Ho | 6,90 |
| PARK7 | Q99497 | Parkinson disease protein 7 OS=Homo sapien | 2,41 |
| ANP32B | Q92688 | Acidic leucine-rich nuclear phosphoprotein 3 | 3,03 |
| PAICS | P22234 | Multifunctional protein ADE2 OS=Homo sapien | 3,08 |
| CD2AP | Q9Y5K6 | CD2-associated protein OS=Homo sapiens OX | 2,11 |

| Ratio n2 | Ratio n3 | Moy | t test |
| --- | --- | --- | --- |
| 72,04 | 467,43 | 290,543087 | 0,129985805 |
| 4,18 | 82,28 | 31,2128474 | 0,358505567 |
| 7,01 | 75,51 | 28,8196271 | 0,355868217 |
| 2,02 | 49,78 | 27,0471302 | 0,200474156 |
| 2,53 | 50,14 | 25,9306694 | 0,211483978 |
| 3,56 | 60,41 | 25,4142334 | 0,301340861 |
| 2,28 | 45,93 | 24,0862614 | 0,208397746 |
| 5,46 | 19,01 | 20,8623478 | 0,170992574 |
| 3,13 | 41,32 | 18,4528109 | 0,272907397 |
| 3,36 | 17,63 | 16,2230582 | 0,16377564 |
| 2,42 | 37,43 | 15,6334345 | 0,314194996 |
| 3,18 | 22,32 | 14,5372264 | 0,145083833 |
| 2,70 | 28,14 | 13,4122854 | 0,244688667 |
| 3,46 | 20,38 | 12,7919581 | 0,140667211 |
| 2,64 | 32,21 | 12,5840349 | 0,359246583 |
| 7,17 | 7,53 | 12,5266296 | 0,155759823 |
| 2,95 | 21,62 | 11,838019 | 0,182938249 |
| 6,44 | 10,59 | 11,2052748 | 0,07413427 |
| 6,35 | 16,28 | 11,1317365 | 0,071775691 |
| 2,91 | 8,24 | 10,9767484 | 0,21762748 |
| 4,10 | 16,18 | 10,8944562 | 0,109134647 |
| 4,67 | 21,04 | 10,2262595 | 0,230019666 |
| 5,41 | 11,08 | 9,9024543 | 0,06215833 |
| 9,87 | 5,85 | 9,76409455 | 0,059119032 |
| 2,30 | 19,74 | 9,75061551 | 0,234081186 |
| 2,84 | 19,60 | 9,6928751 | 0,228715094 |
| 2,22 | 21,30 | 9,52175149 | 0,288017705 |
| 2,55 | 17,09 | 9,28276901 | 0,189358334 |
| 2,46 | 20,41 | 8,95990608 | 0,299953559 |
| 3,39 | 17,86 | 8,55570285 | 0,246501168 |
| 2,97 | 8,57 | 8,50694117 | 0,142103167 |
| 2,03 | 18,27 | 7,85436072 | 0,319673648 |
| 16,60 | 2,28 | 7,79352329 | 0,266455364 |
| 2,84 | 5,35 | 7,7276005 | 0,211196383 |
| 2,55 | 11,55 | 7,58824753 | 0,131145972 |
| 3,63 | 9,70 | 7,19427661 | 0,077409488 |
| 2,27 | 11,22 | 7,18158088 | 0,14219254 |
| 3,63 | 6,95 | 7,02185533 | 0,093285417 |
| 2,55 | 11,55 | 6,76173071 | 0,158438577 |
| 2,48 | 8,92 | 6,6121763 | 0,11359837 |
| 3,20 | 13,93 | 6,46669141 | 0,281256065 |
| 5,90 | 6,10 | 6,37819292 | 0,005060971 |
| 2,08 | 7,63 | 6,37120803 | 0,135178276 |
| 2,27 | 12,94 | 6,21650321 | 0,262592507 |
| 2,51 | 12,60 | 6,15475924 | 0,251529096 |
| 3,97 | 3,49 | 6,04767308 | 0,161754901 |
| 2,09 | 11,96 | 5,45557411 | 0,304122486 |
| 2,00 | 3,67 | 5,43414481 | 0,235382761 |
| 2,63 | 5,27 | 5,3603105 | 0,112922835 |
| 2,35 | 6,91 | 4,94909382 | 0,100244817 |
| 2,27 | 5,51 | 4,89576222 | 0,104856841 |
| 2,01 | 9,82 | 4,74685491 | 0,278072416 |
| 3,29 | 7,02 | 4,44669684 | 0,115680659 |
| 2,45 | 3,82 | 3,11557482 | 0,033637494 |
| 2,59 | 2,89 | 2,52632942 | 0,02144601 |

**Supplemental Table 2:**
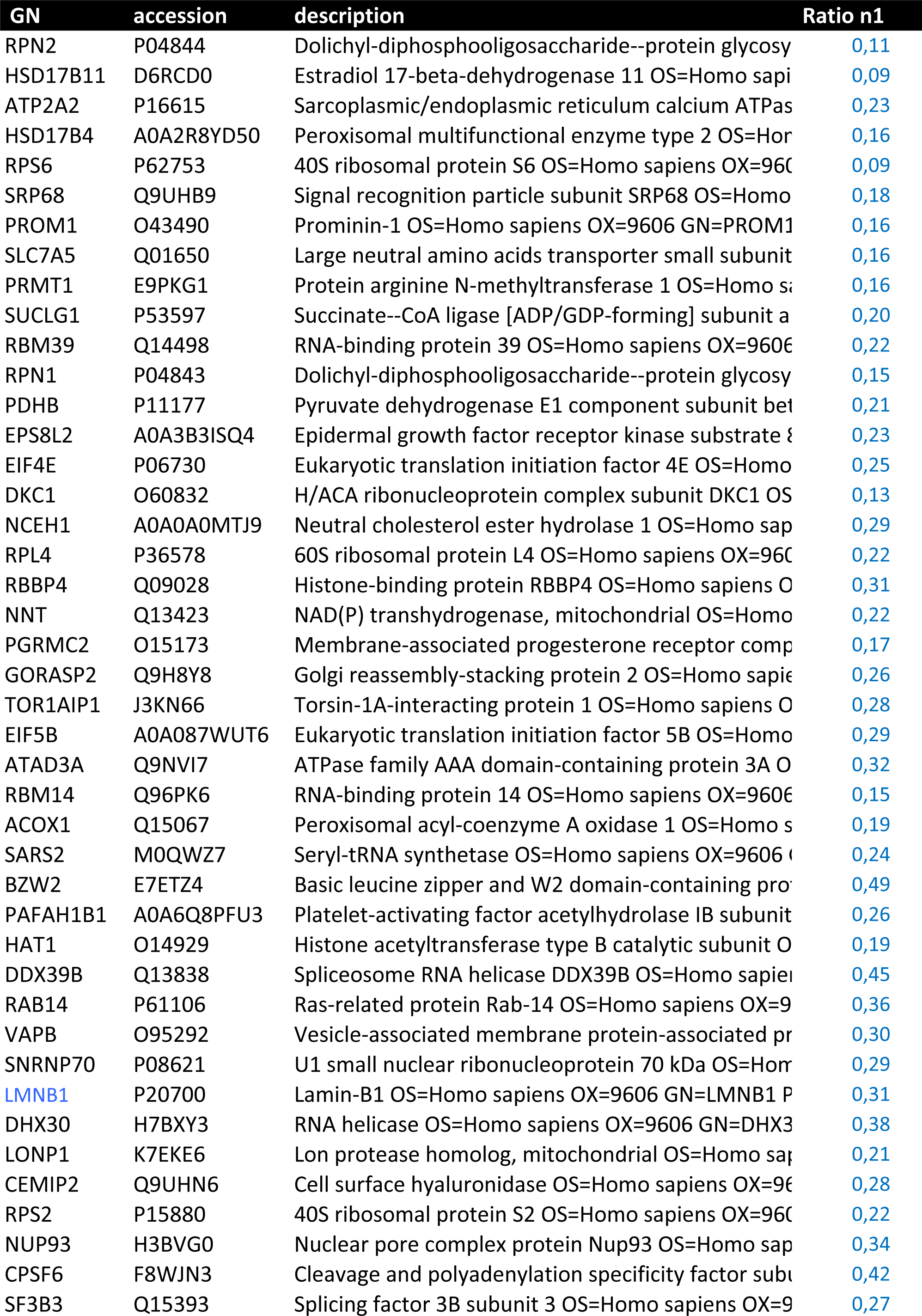
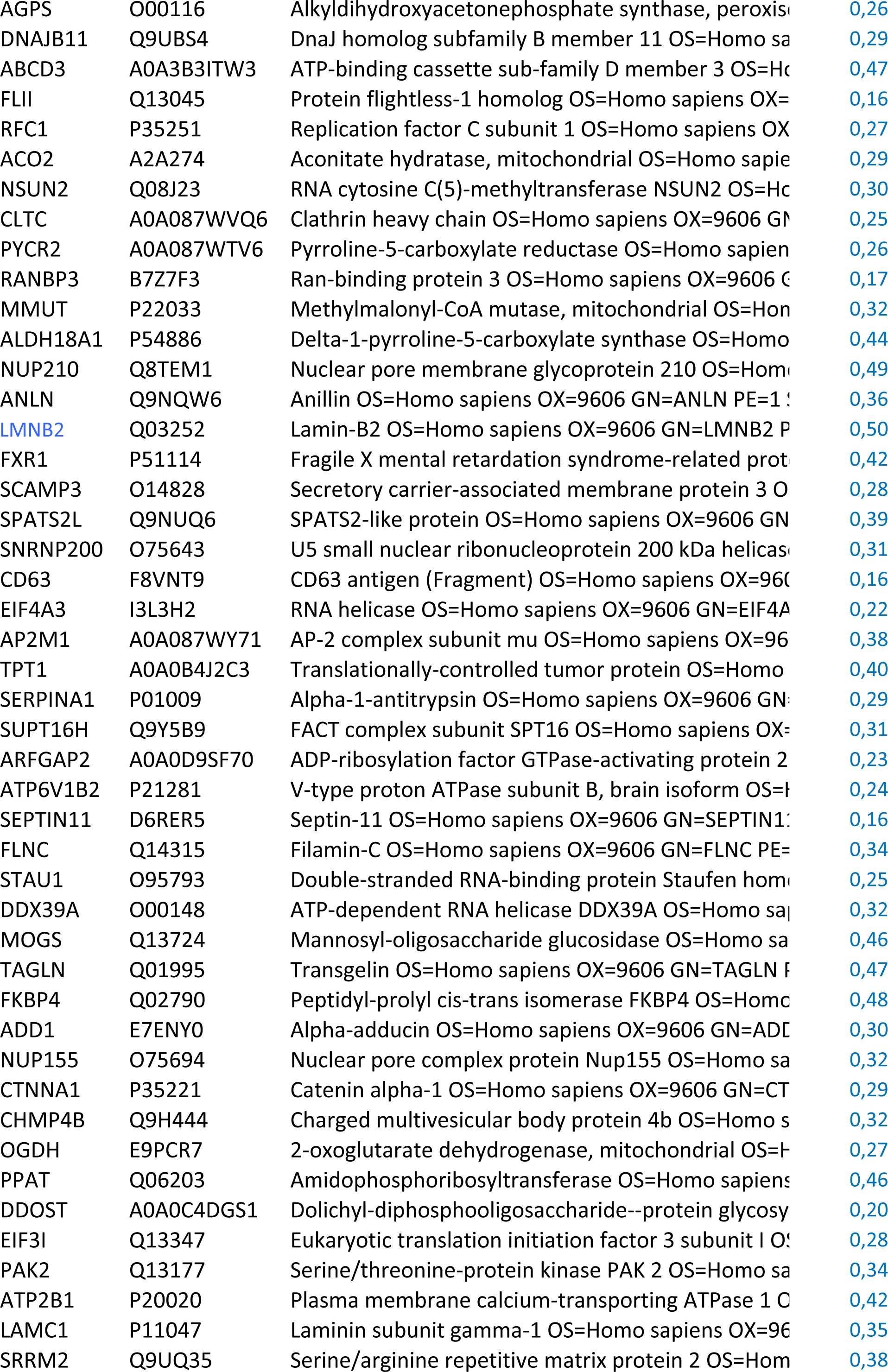
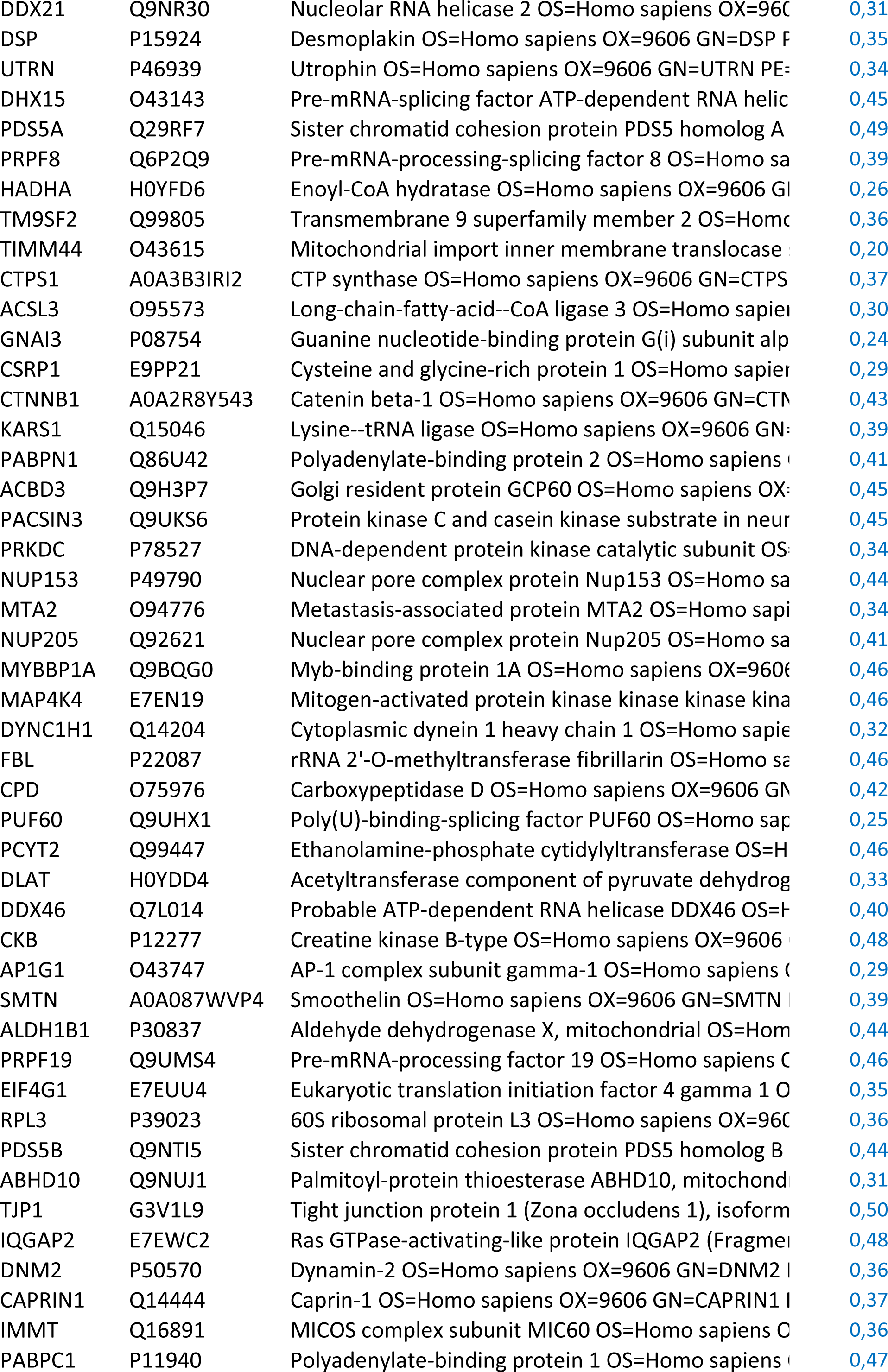
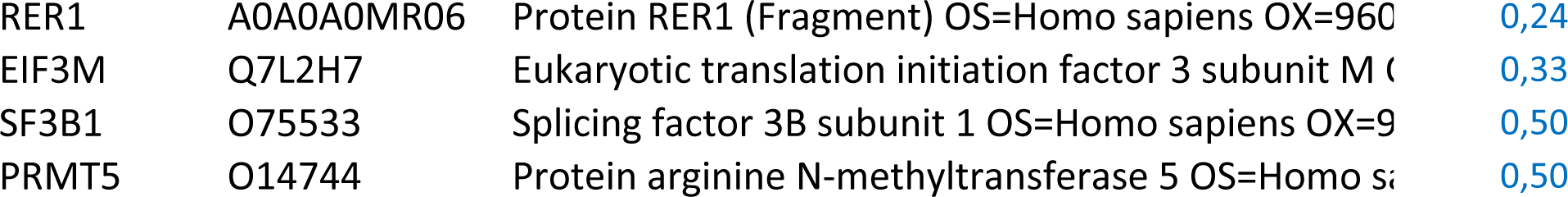
Results of the mass spectrometry analysis. List of proteins underrepresented in the entotic fraction.

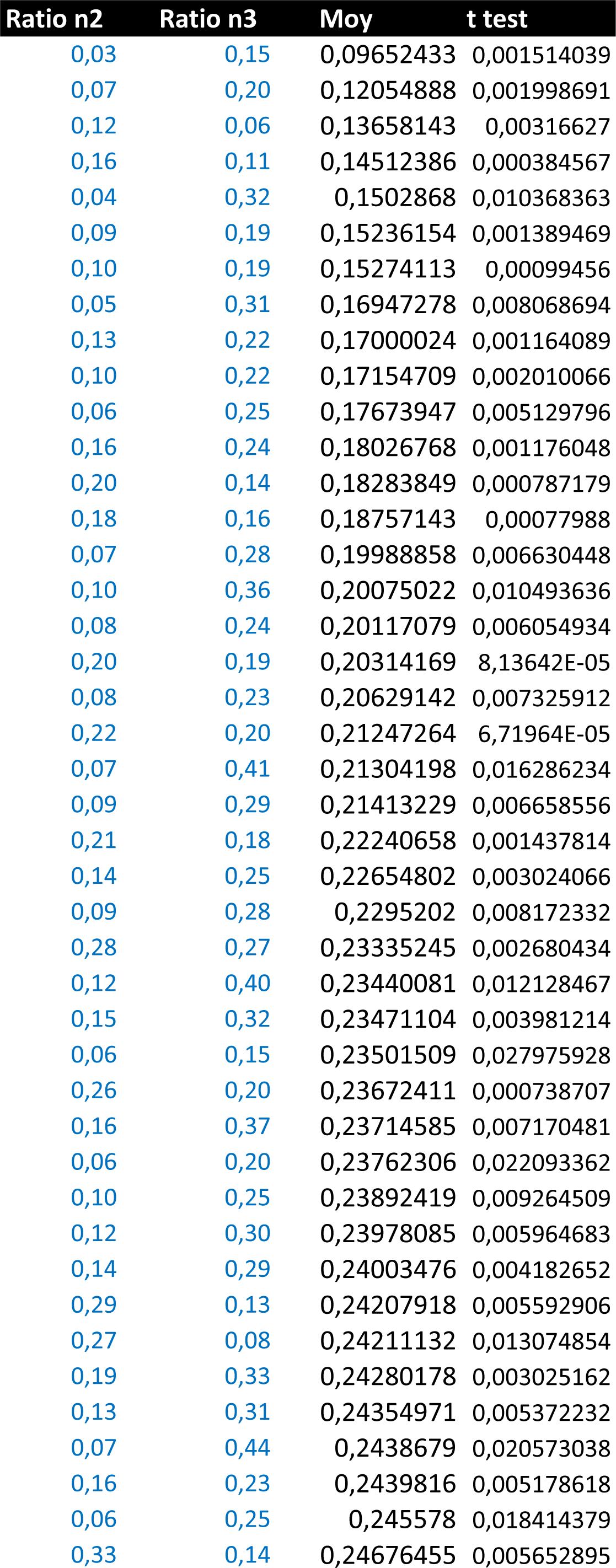
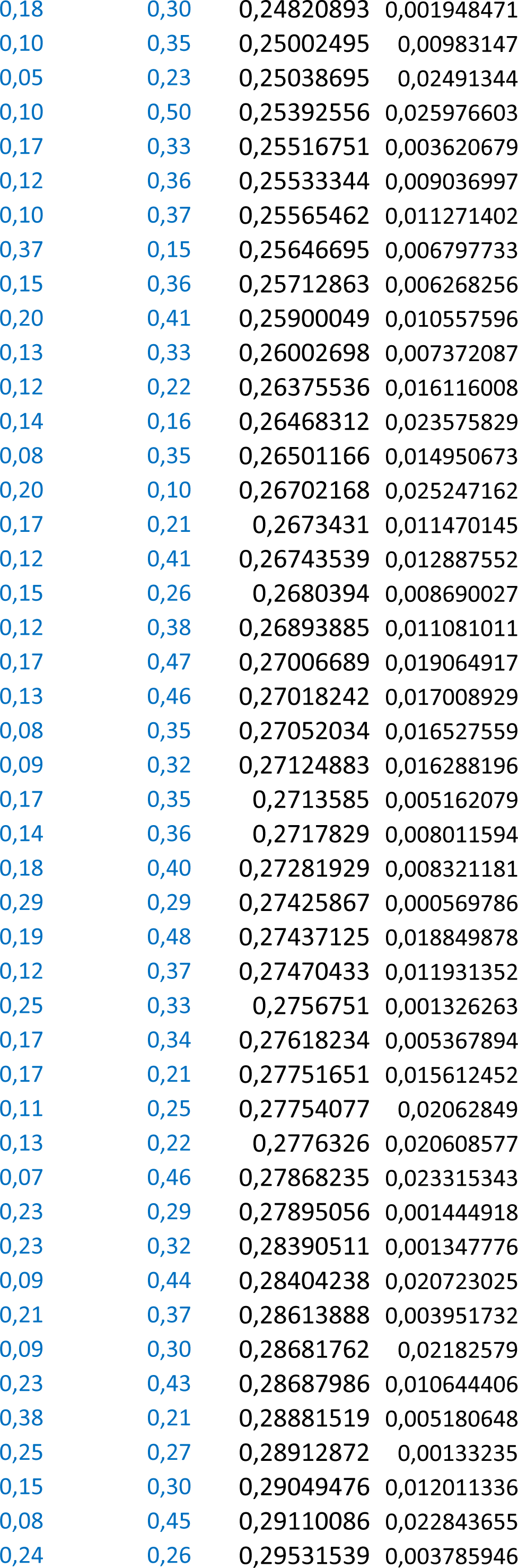
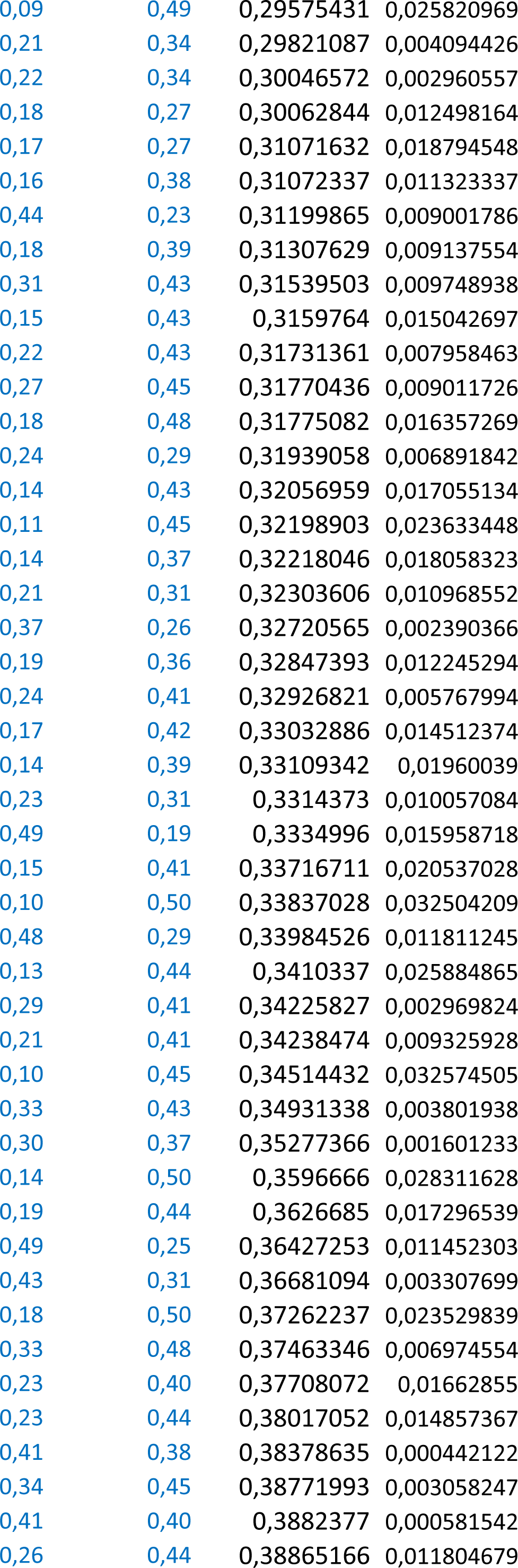
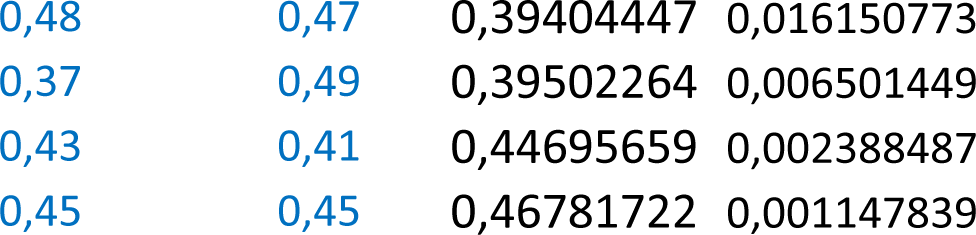

**Supplemental Table 3:**
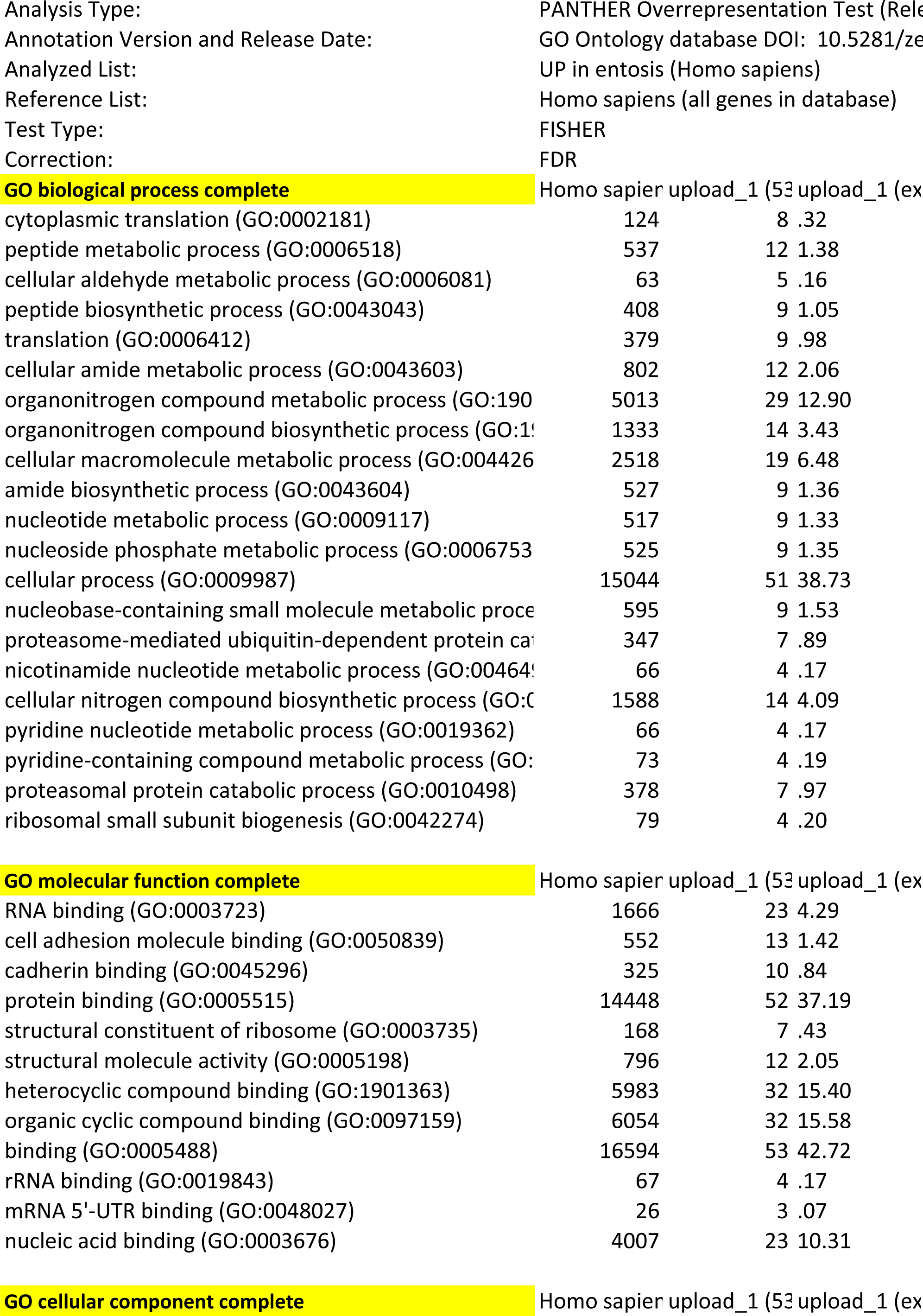
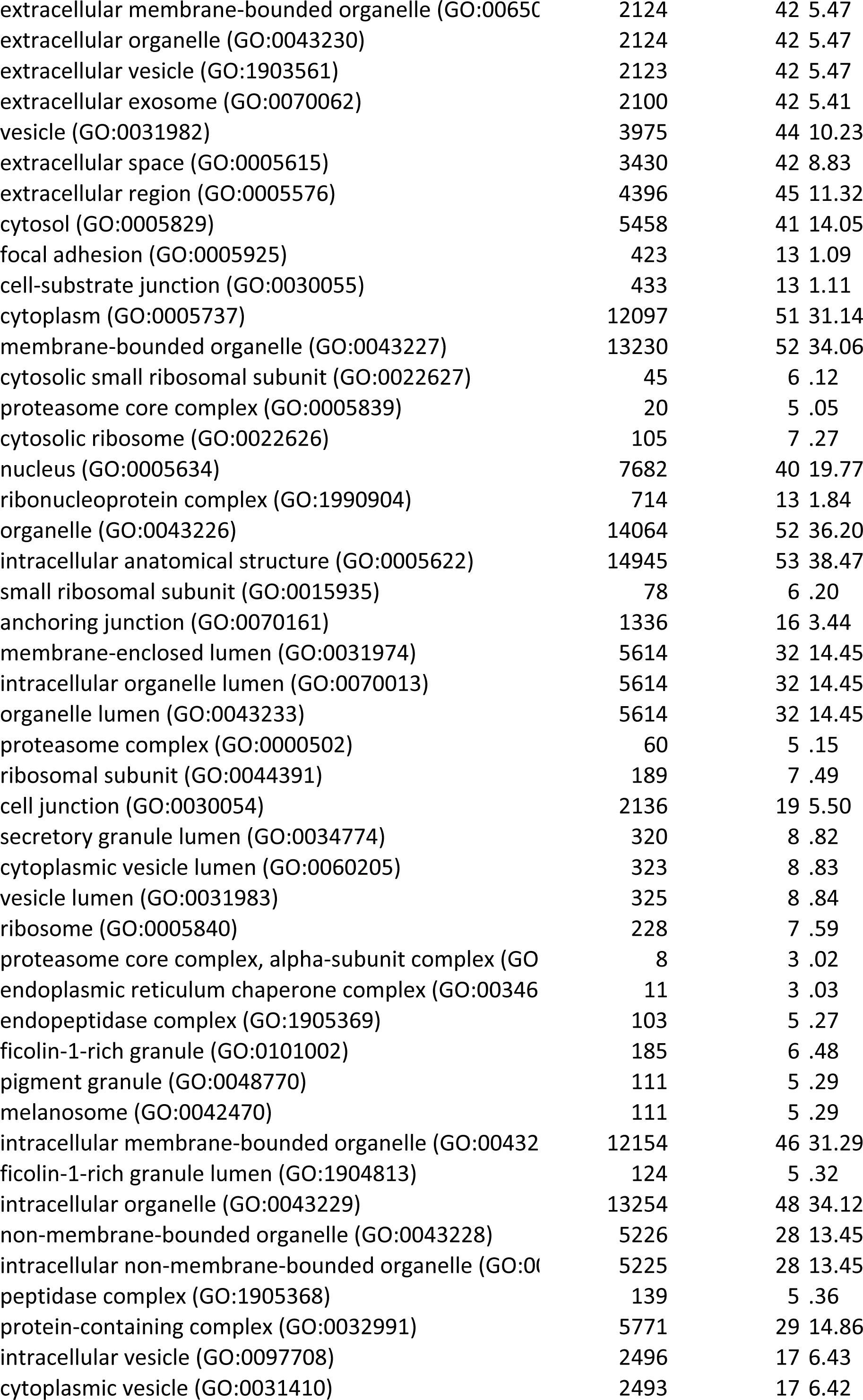
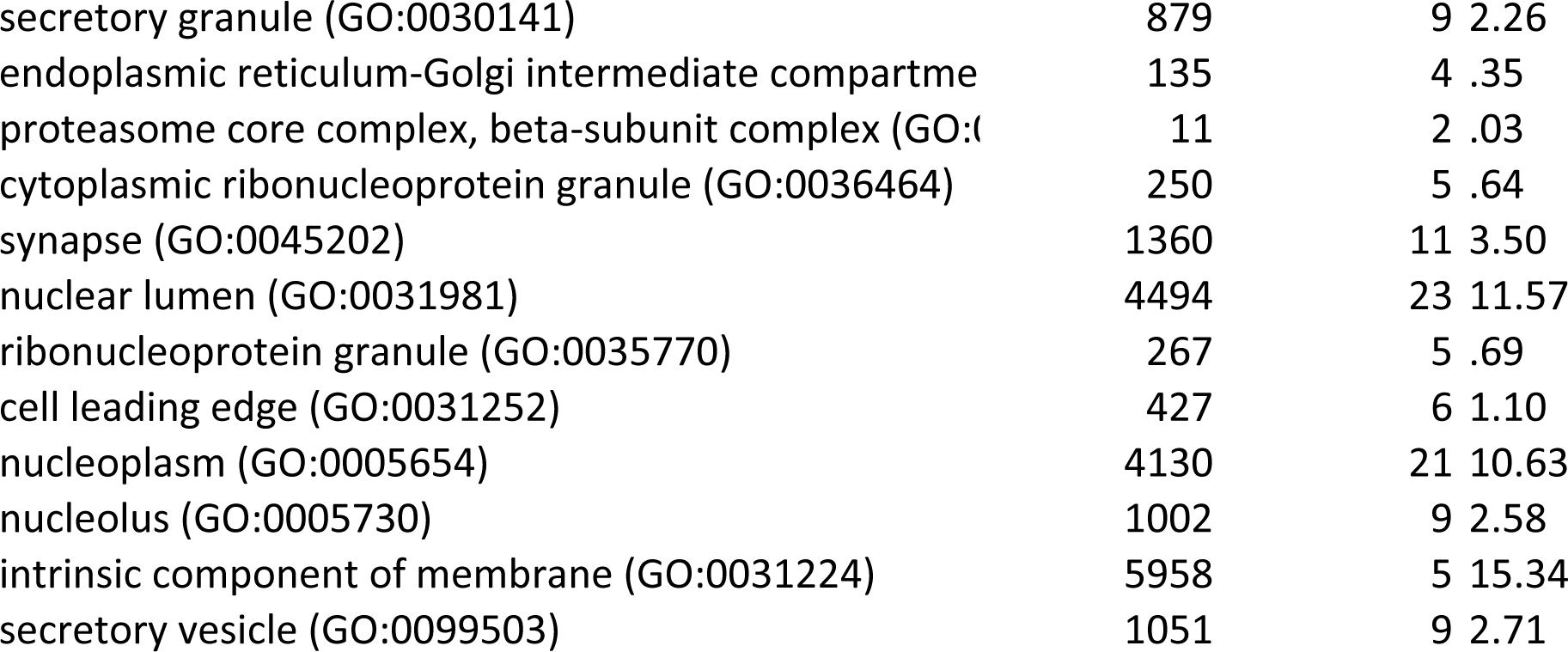
Results of the mass spectrometry analysis. GO analysis on proteins found overrepresented in the entotic fraction.

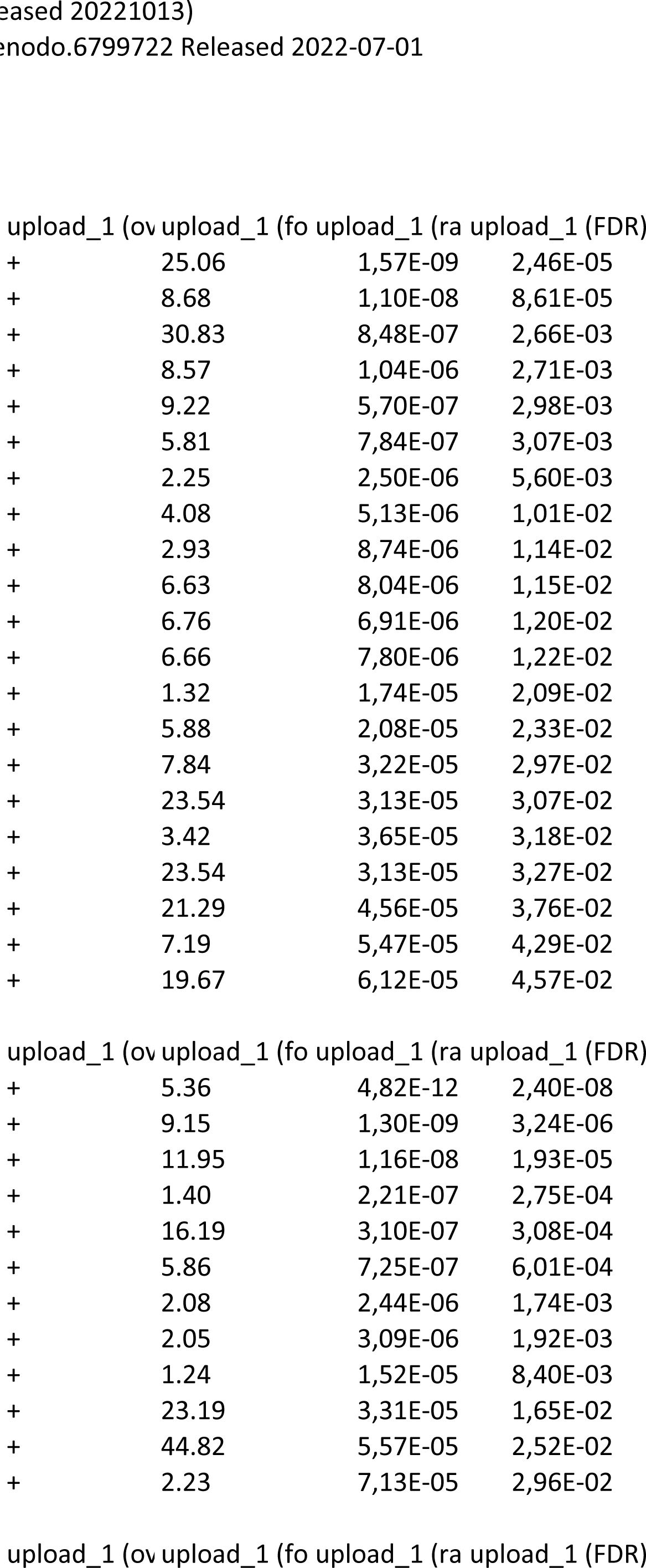
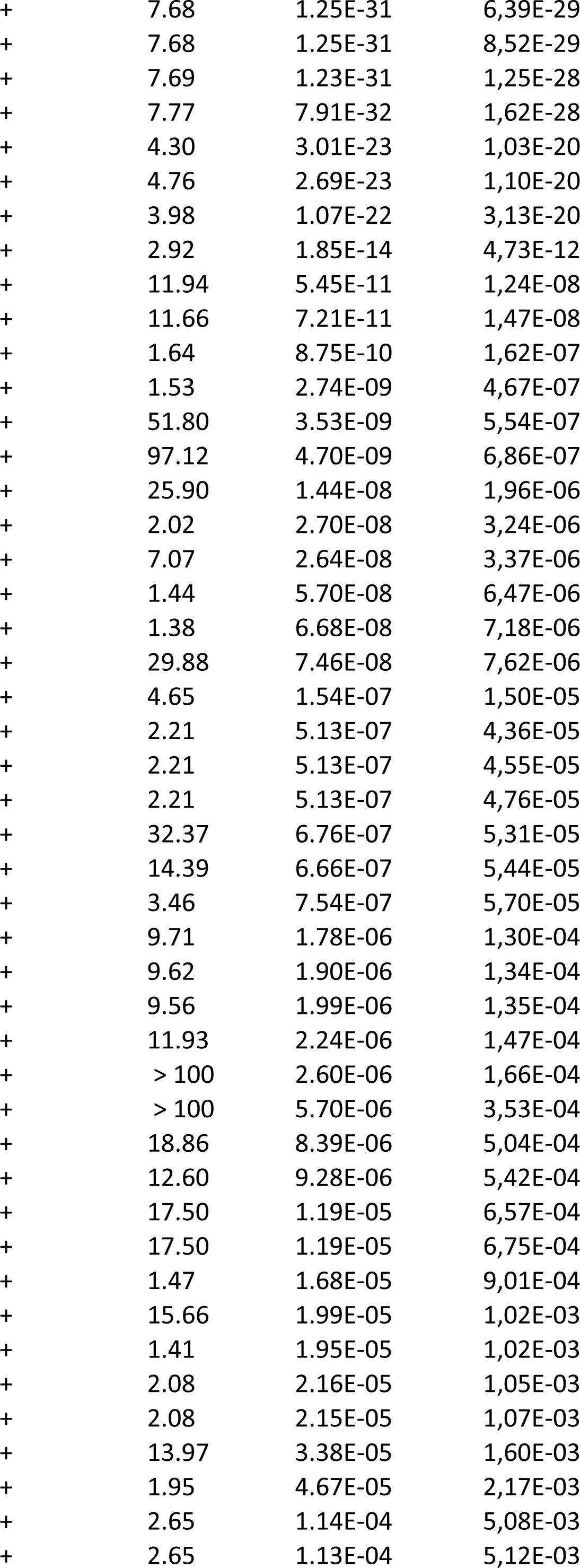
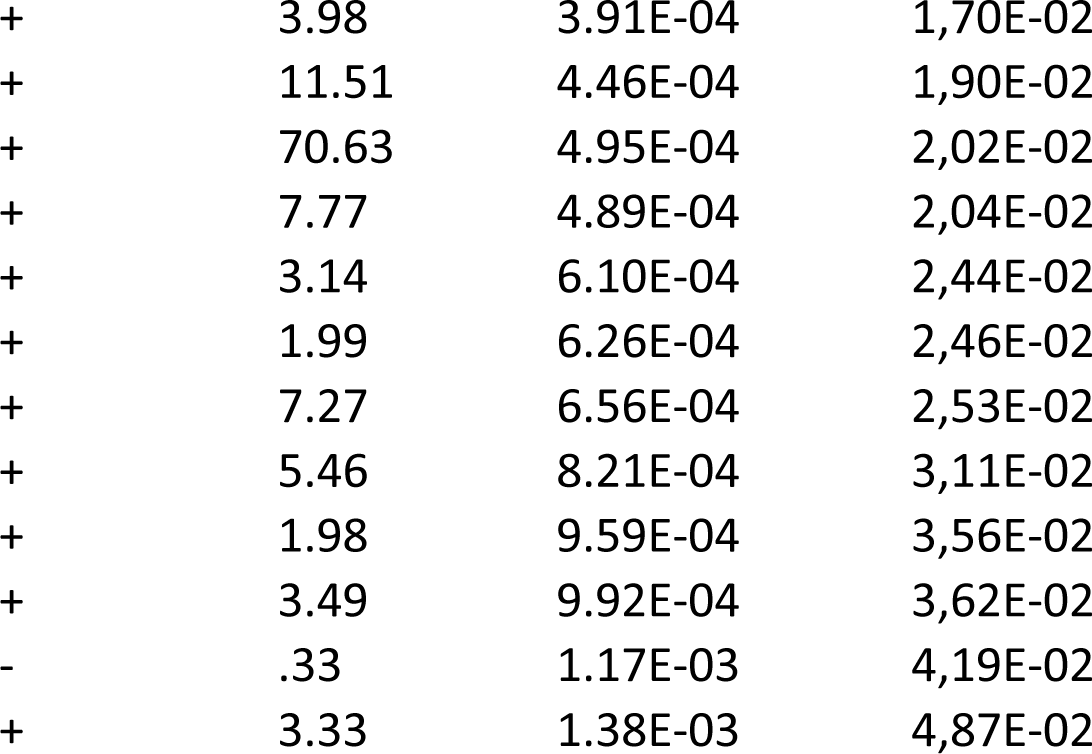

|  |  |  |  |
| --- | --- | --- | --- |
| + | 25.06 | 1,57E-09 | 2,46E-05 |
| + | 8.68 | 1,10E-08 | 8,61E-05 |
| + | 30.83 | 8,48E-07 | 2,66E-03 |
| + | 8.57 | 1,04E-06 | 2,71E-03 |
| + | 9.22 | 5,70E-07 | 2,98E-03 |
| + | 5.81 | 7,84E-07 | 3,07E-03 |
| + | 2.25 | 2,50E-06 | 5,60E-03 |
| + | 4.08 | 5,13E-06 | 1,01E-02 |
| + | 2.93 | 8,74E-06 | 1,14E-02 |
| + | 6.63 | 8,04E-06 | 1,15E-02 |
| + | 6.76 | 6,91E-06 | 1,20E-02 |
| + | 6.66 | 7,80E-06 | 1,22E-02 |
| + | 1.32 | 1,74E-05 | 2,09E-02 |
| + | 5.88 | 2,08E-05 | 2,33E-02 |
| + | 7.84 | 3,22E-05 | 2,97E-02 |
| + | 23.54 | 3,13E-05 | 3,07E-02 |
| + | 3.42 | 3,65E-05 | 3,18E-02 |
| + | 23.54 | 3,13E-05 | 3,27E-02 |
| + | 21.29 | 4,56E-05 | 3,76E-02 |
| + | 7.19 | 5,47E-05 | 4,29E-02 |
| + | 19.67 | 6,12E-05 | 4,57E-02 |
| + | 5.36 | 4,82E-12 | 2,40E-08 |
| + | 9.15 | 1,30E-09 | 3,24E-06 |
| + | 11.95 | 1,16E-08 | 1,93E-05 |
| + | 1.40 | 2,21E-07 | 2,75E-04 |
| + | 16.19 | 3,10E-07 | 3,08E-04 |
| + | 5.86 | 7,25E-07 | 6,01E-04 |
| + | 2.08 | 2,44E-06 | 1,74E-03 |
| + | 2.05 | 3,09E-06 | 1,92E-03 |
| + | 1.24 | 1,52E-05 | 8,40E-03 |
| + | 23.19 | 3,31E-05 | 1,65E-02 |
| + | 44.82 | 5,57E-05 | 2,52E-02 |
| + | 2.23 | 7,13E-05 | 2,96E-02 |
| + | 7.68 | 1.25E-31 | 6,39E-29 |
| + | 7.68 | 1.25E-31 | 8,52E-29 |
| + | 7.69 | 1.23E-31 | 1,25E-28 |
| + | 7.77 | 7.91E-32 | 1,62E-28 |
| + | 4.30 | 3.01E-23 | 1,03E-20 |
| + | 4.76 | 2.69E-23 | 1,10E-20 |
| + | 3.98 | 1.07E-22 | 3,13E-20 |
| + | 2.92 | 1.85E-14 | 4,73E-12 |
| + | 11.94 | 5.45E-11 | 1,24E-08 |
| + | 11.66 | 7.21E-11 | 1,47E-08 |
| + | 1.64 | 8.75E-10 | 1,62E-07 |
| + | 1.53 | 2.74E-09 | 4,67E-07 |
| + | 51.80 | 3.53E-09 | 5,54E-07 |
| + | 97.12 | 4.70E-09 | 6,86E-07 |
| + | 25.90 | 1.44E-08 | 1,96E-06 |
| + | 2.02 | 2.70E-08 | 3,24E-06 |
| + | 7.07 | 2.64E-08 | 3,37E-06 |
| + | 1.44 | 5.70E-08 | 6,47E-06 |
| + | 1.38 | 6.68E-08 | 7,18E-06 |
| + | 29.88 | 7.46E-08 | 7,62E-06 |
| + | 4.65 | 1.54E-07 | 1,50E-05 |
| + | 2.21 | 5.13E-07 | 4,36E-05 |
| + | 2.21 | 5.13E-07 | 4,55E-05 |
| + | 2.21 | 5.13E-07 | 4,76E-05 |
| + | 32.37 | 6.76E-07 | 5,31E-05 |
| + | 14.39 | 6.66E-07 | 5,44E-05 |
| + | 3.46 | 7.54E-07 | 5,70E-05 |
| + | 9.71 | 1.78E-06 | 1,30E-04 |
| + | 9.62 | 1.90E-06 | 1,34E-04 |
| + | 9.56 | 1.99E-06 | 1,35E-04 |
| + | 11.93 | 2.24E-06 | 1,47E-04 |
| + | > 100 | 2.60E-06 | 1,66E-04 |
| + | > 100 | 5.70E-06 | 3,53E-04 |
| + | 18.86 | 8.39E-06 | 5,04E-04 |
| + | 12.60 | 9.28E-06 | 5,42E-04 |
| + | 17.50 | 1.19E-05 | 6,57E-04 |
| + | 17.50 | 1.19E-05 | 6,75E-04 |
| + | 1.47 | 1.68E-05 | 9,01E-04 |
| + | 15.66 | 1.99E-05 | 1,02E-03 |
| + | 1.41 | 1.95E-05 | 1,02E-03 |
| + | 2.08 | 2.16E-05 | 1,05E-03 |
| + | 2.08 | 2.15E-05 | 1,07E-03 |
| + | 13.97 | 3.38E-05 | 1,60E-03 |
| + | 1.95 | 4.67E-05 | 2,17E-03 |
| + | 2.65 | 1.14E-04 | 5,08E-03 |
| + | 2.65 | 1.13E-04 | 5,12E-03 |
| + | 3.98 | 3.91E-04 | 1,70E-02 |
| + | 11.51 | 4.46E-04 | 1,90E-02 |
| + | 70.63 | 4.95E-04 | 2,02E-02 |
| + | 7.77 | 4.89E-04 | 2,04E-02 |
| + | 3.14 | 6.10E-04 | 2,44E-02 |
| + | 1.99 | 6.26E-04 | 2,46E-02 |
| + | 7.27 | 6.56E-04 | 2,53E-02 |
| + | 5.46 | 8.21E-04 | 3,11E-02 |
| + | 1.98 | 9.59E-04 | 3,56E-02 |
| + | 3.49 | 9.92E-04 | 3,62E-02 |
| - | .33 | 1.17E-03 | 4,19E-02 |
| + | 3.33 | 1.38E-03 | 4,87E-02 |

**Supplemental Table 4:**
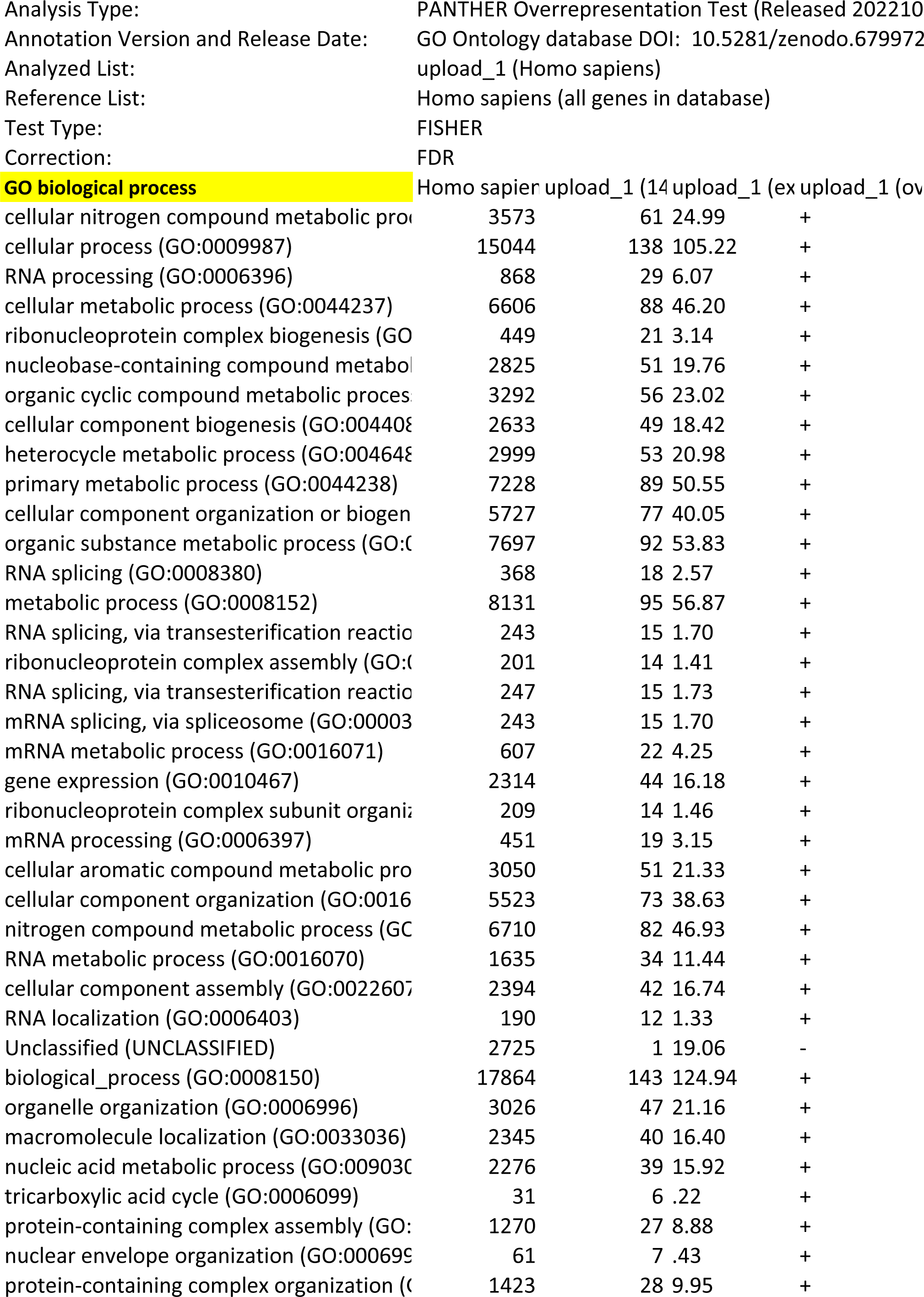
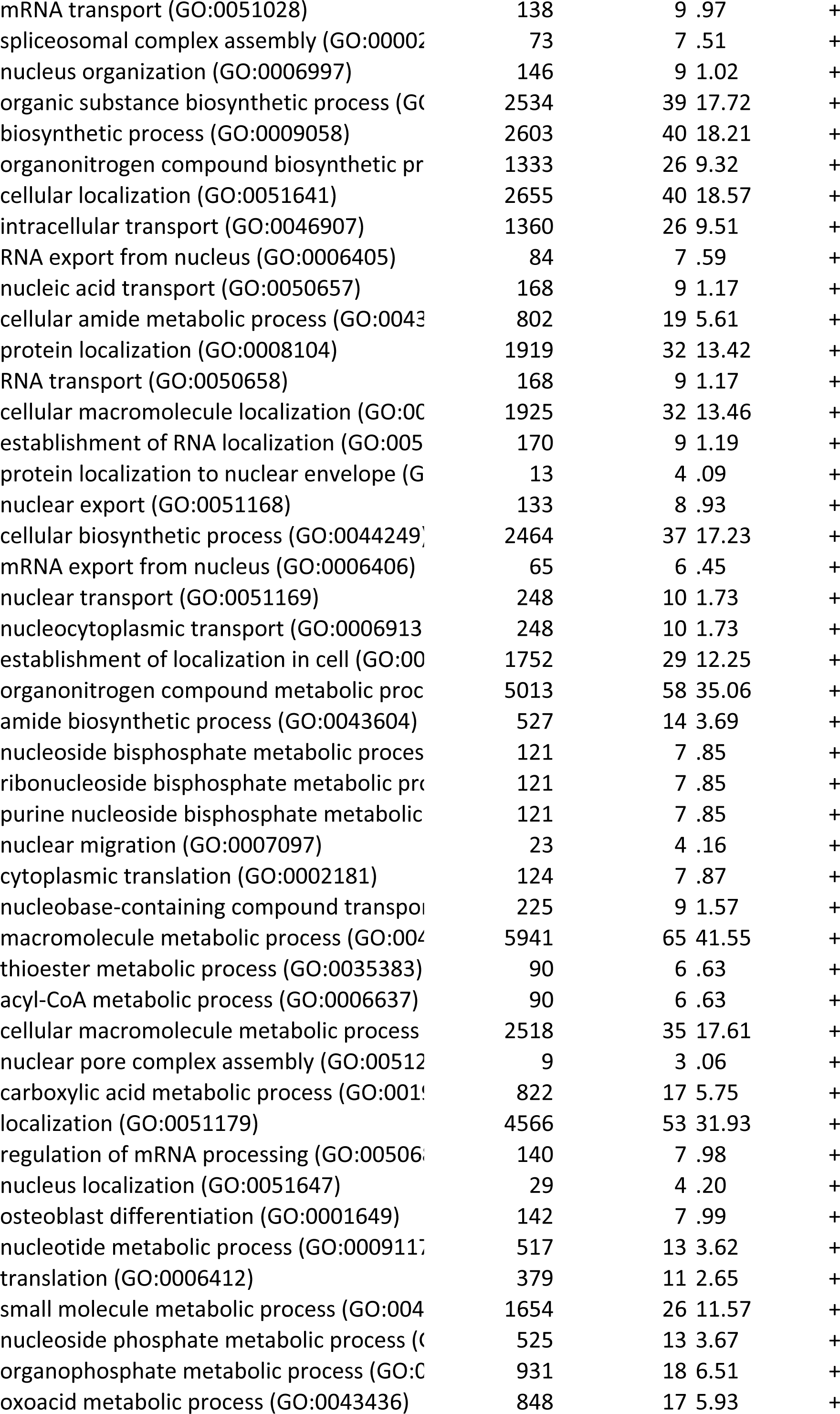
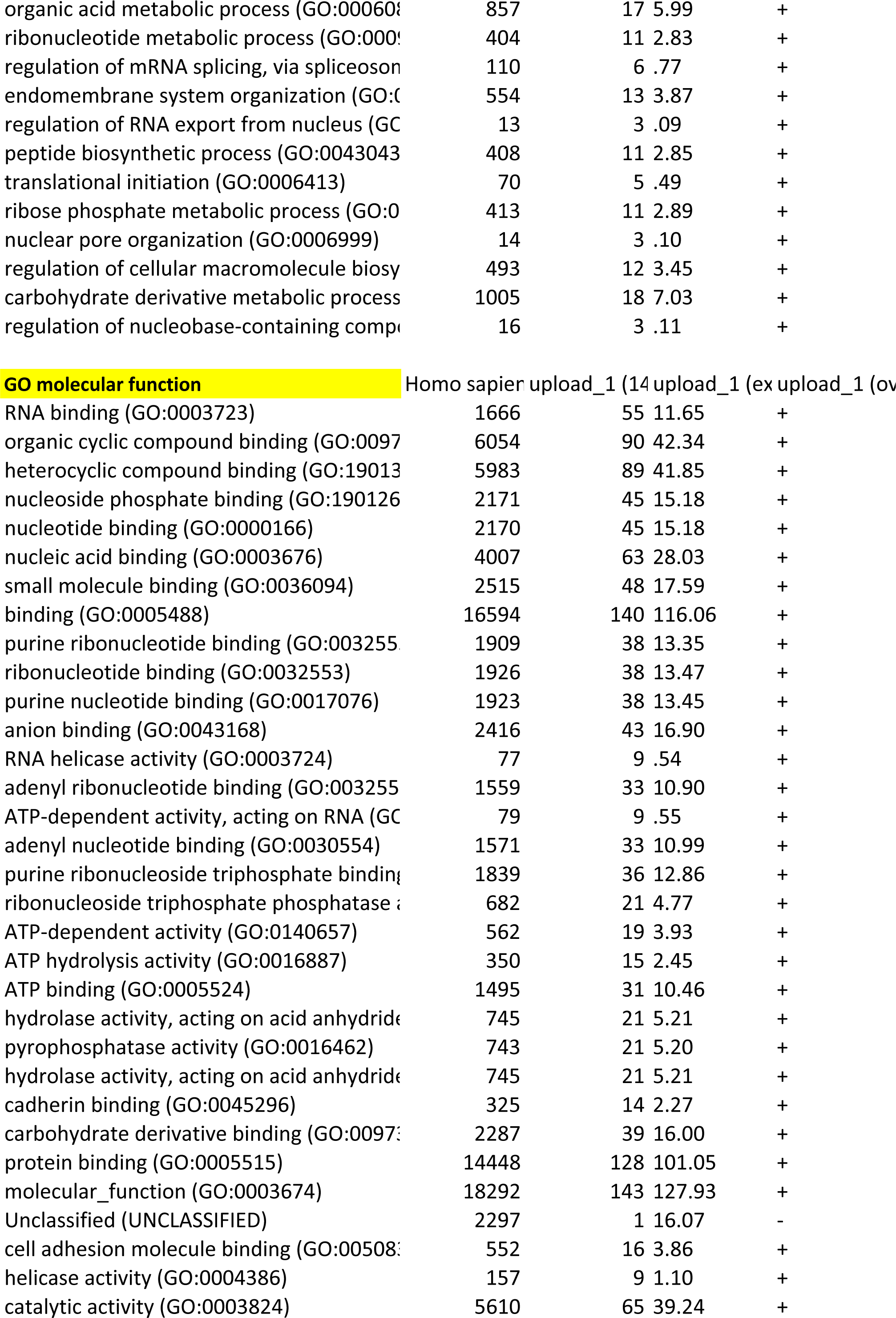
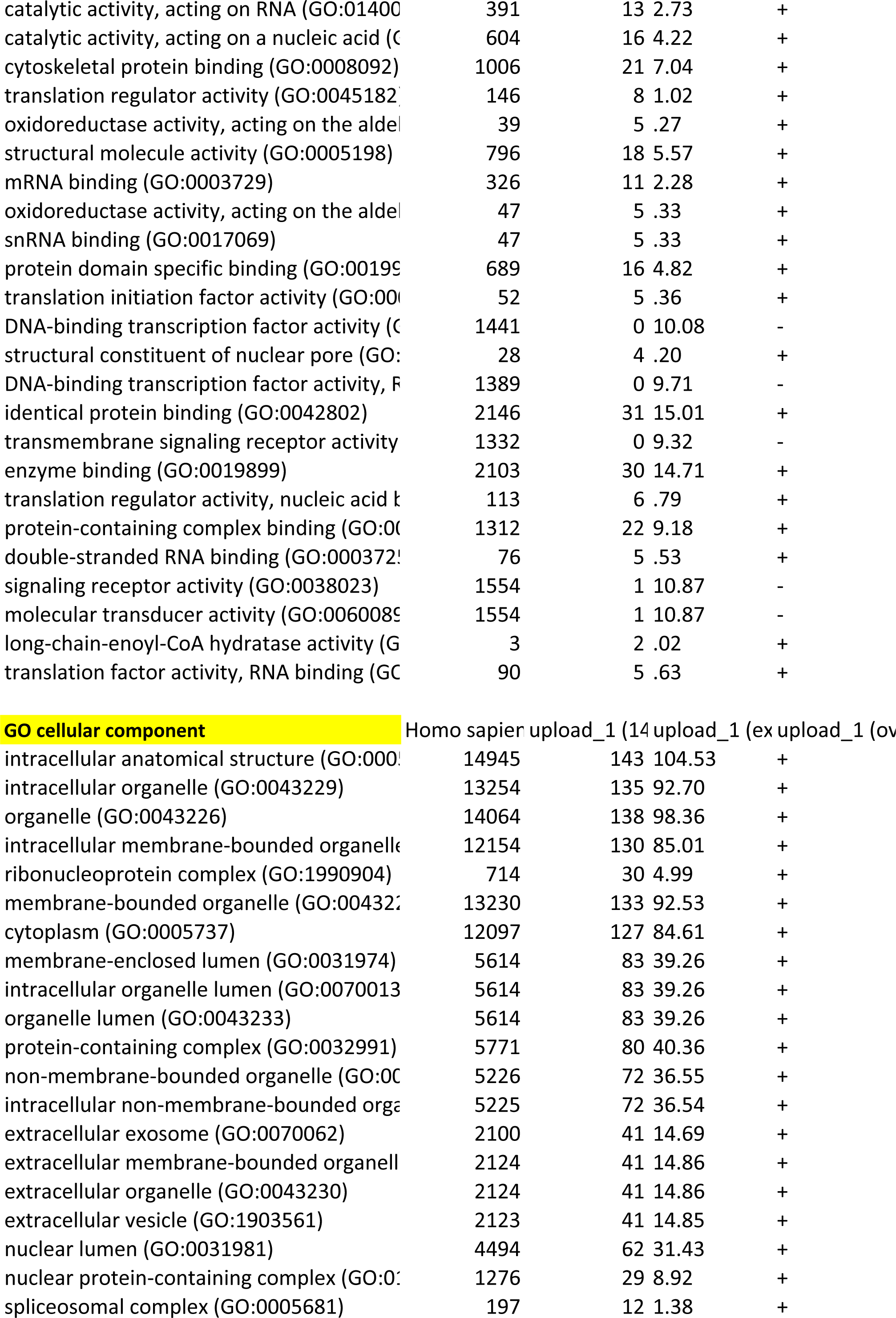
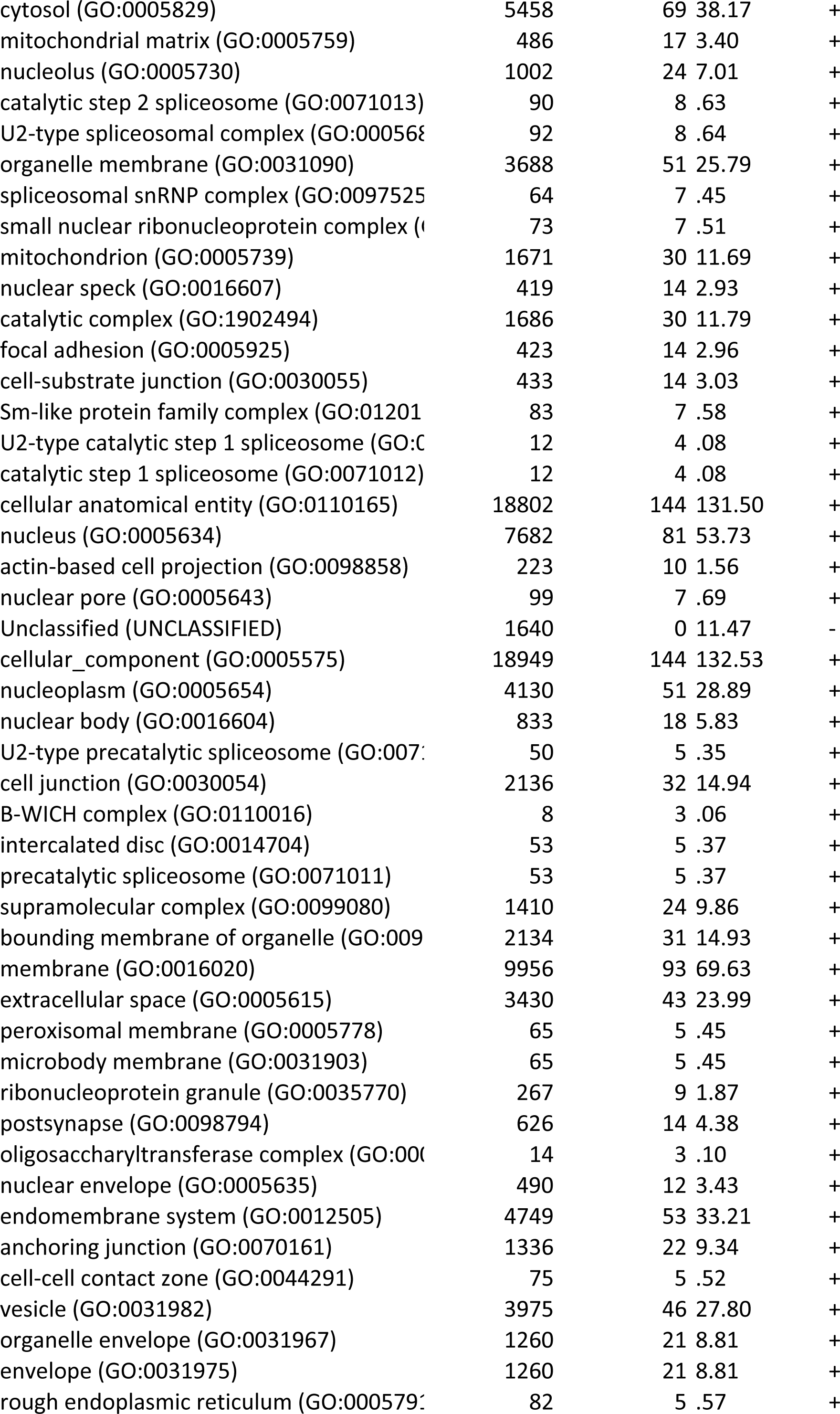
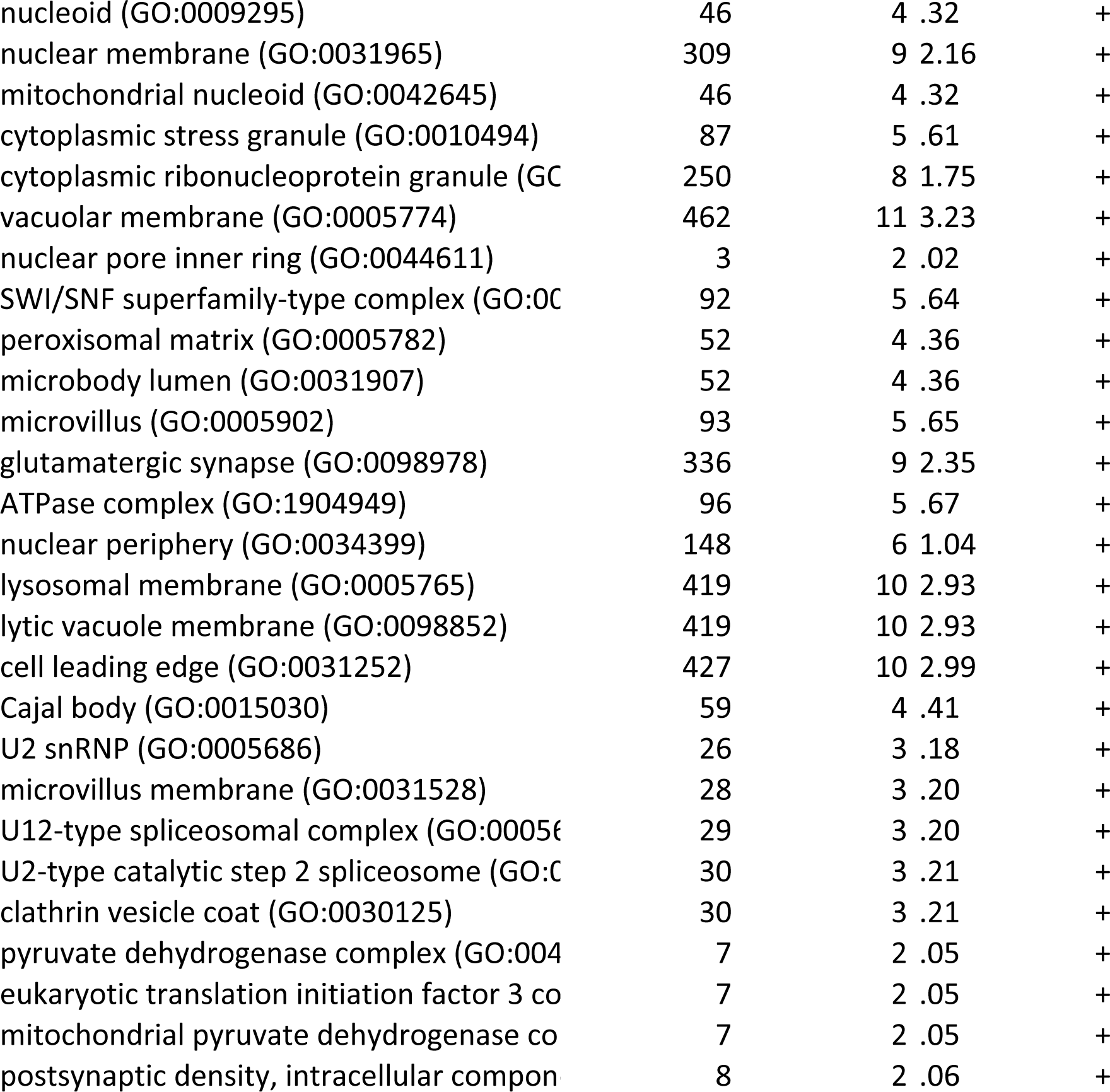
Results of the mass spectrometry analysis. GO analysis on proteins found underrepresented in the entotic fraction.

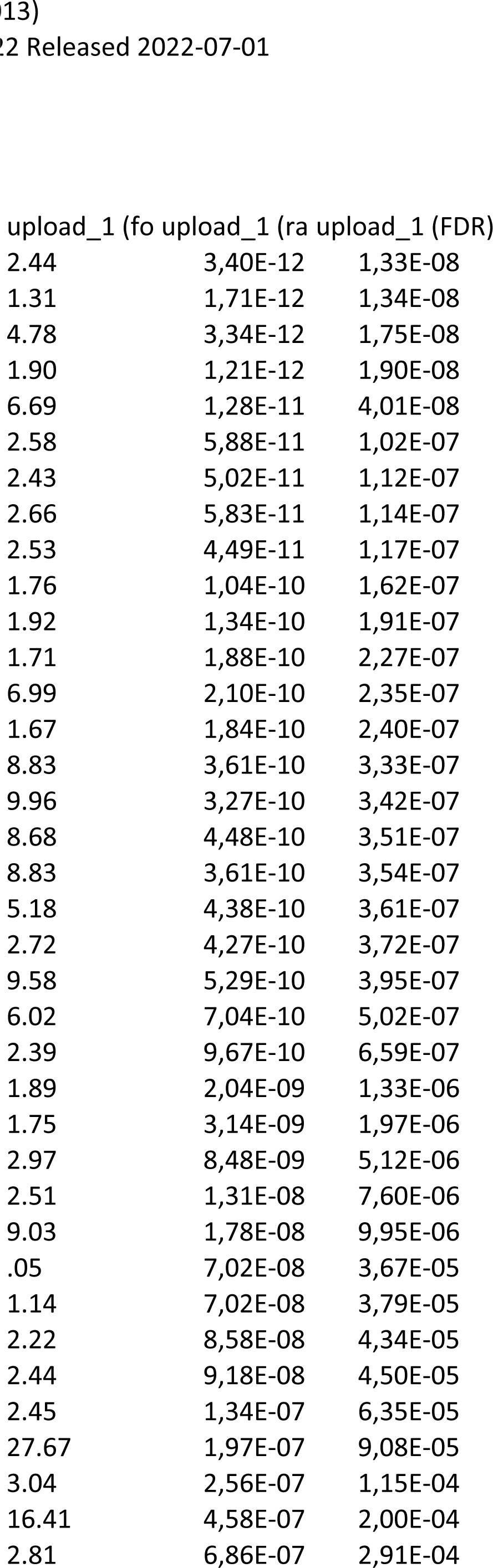
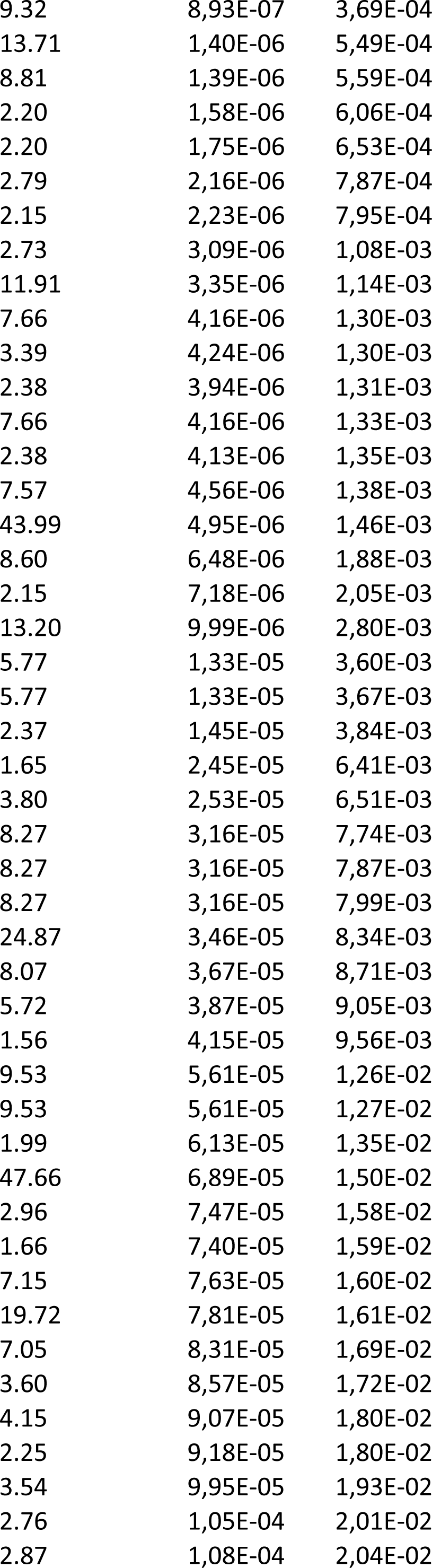
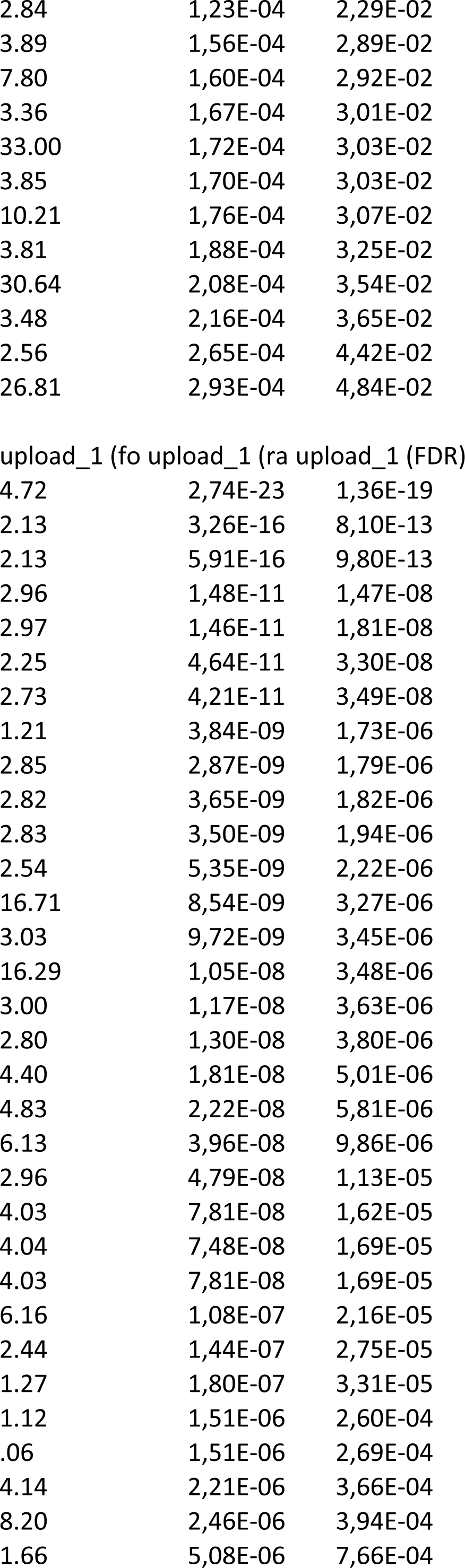
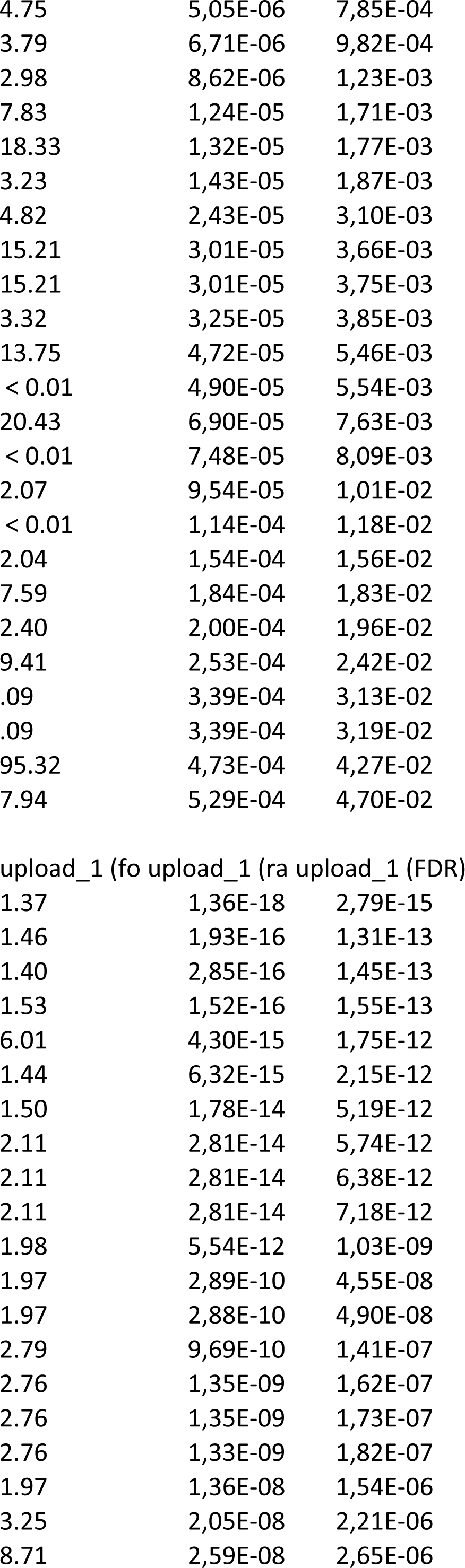
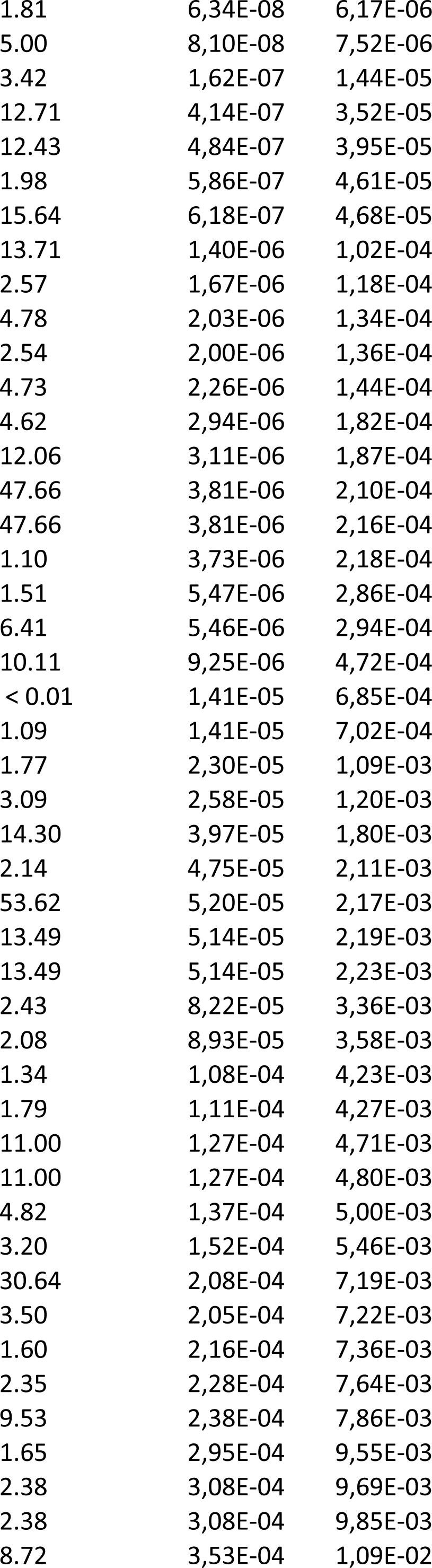
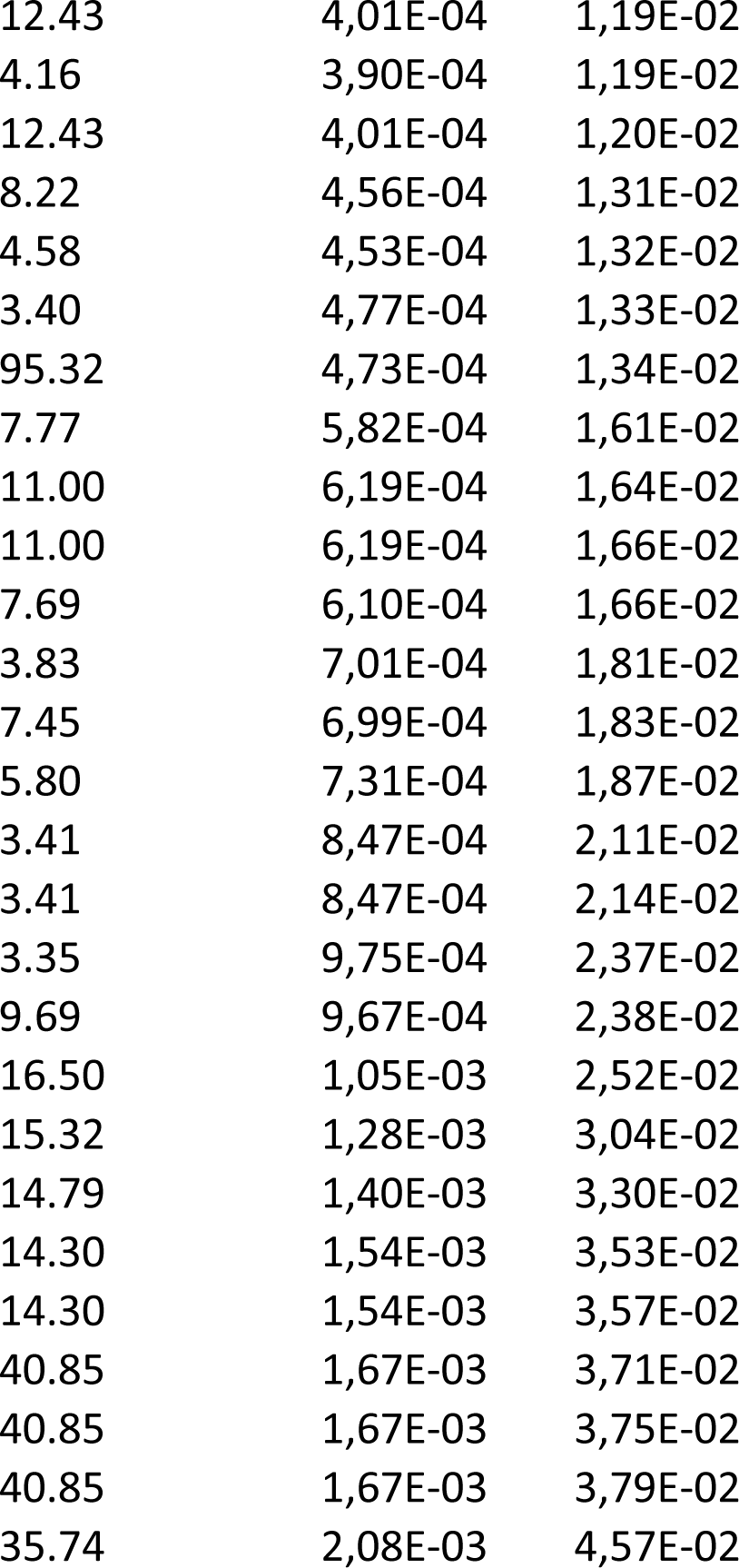

|  |  |  |
| --- | --- | --- |
| 2.44 | 3,40E-12 | 1,33E-08 |
| 1.31 | 1,71E-12 | 1,34E-08 |
| 4.78 | 3,34E-12 | 1,75E-08 |
| 1.90 | 1,21E-12 | 1,90E-08 |
| 6.69 | 1,28E-11 | 4,01E-08 |
| 2.58 | 5,88E-11 | 1,02E-07 |
| 2.43 | 5,02E-11 | 1,12E-07 |
| 2.66 | 5,83E-11 | 1,14E-07 |
| 2.53 | 4,49E-11 | 1,17E-07 |
| 1.76 | 1,04E-10 | 1,62E-07 |
| 1.92 | 1,34E-10 | 1,91E-07 |
| 1.71 | 1,88E-10 | 2,27E-07 |
| 6.99 | 2,10E-10 | 2,35E-07 |
| 1.67 | 1,84E-10 | 2,40E-07 |
| 8.83 | 3,61E-10 | 3,33E-07 |
| 9.96 | 3,27E-10 | 3,42E-07 |
| 8.68 | 4,48E-10 | 3,51E-07 |
| 8.83 | 3,61E-10 | 3,54E-07 |
| 5.18 | 4,38E-10 | 3,61E-07 |
| 2.72 | 4,27E-10 | 3,72E-07 |
| 9.58 | 5,29E-10 | 3,95E-07 |
| 6.02 | 7,04E-10 | 5,02E-07 |
| 2.39 | 9,67E-10 | 6,59E-07 |
| 1.89 | 2,04E-09 | 1,33E-06 |
| 1.75 | 3,14E-09 | 1,97E-06 |
| 2.97 | 8,48E-09 | 5,12E-06 |
| 2.51 | 1,31E-08 | 7,60E-06 |
| 9.03 | 1,78E-08 | 9,95E-06 |
| .05 | 7,02E-08 | 3,67E-05 |
| 1.14 | 7,02E-08 | 3,79E-05 |
| 2.22 | 8,58E-08 | 4,34E-05 |
| 2.44 | 9,18E-08 | 4,50E-05 |
| 2.45 | 1,34E-07 | 6,35E-05 |
| 27.67 | 1,97E-07 | 9,08E-05 |
| 3.04 | 2,56E-07 | 1,15E-04 |
| 16.41 | 4,58E-07 | 2,00E-04 |
| 2.81 | 6,86E-07 | 2,91E-04 |
| 9.32 | 8,93E-07 | 3,69E-04 |
| 13.71 | 1,40E-06 | 5,49E-04 |
| 8.81 | 1,39E-06 | 5,59E-04 |
| 2.20 | 1,58E-06 | 6,06E-04 |
| 2.20 | 1,75E-06 | 6,53E-04 |
| 2.79 | 2,16E-06 | 7,87E-04 |
| 2.15 | 2,23E-06 | 7,95E-04 |
| 2.73 | 3,09E-06 | 1,08E-03 |
| 11.91 | 3,35E-06 | 1,14E-03 |
| 7.66 | 4,16E-06 | 1,30E-03 |
| 3.39 | 4,24E-06 | 1,30E-03 |
| 2.38 | 3,94E-06 | 1,31E-03 |
| 7.66 | 4,16E-06 | 1,33E-03 |
| 2.38 | 4,13E-06 | 1,35E-03 |
| 7.57 | 4,56E-06 | 1,38E-03 |
| 43.99 | 4,95E-06 | 1,46E-03 |
| 8.60 | 6,48E-06 | 1,88E-03 |
| 2.15 | 7,18E-06 | 2,05E-03 |
| 13.20 | 9,99E-06 | 2,80E-03 |
| 5.77 | 1,33E-05 | 3,60E-03 |
| 5.77 | 1,33E-05 | 3,67E-03 |
| 2.37 | 1,45E-05 | 3,84E-03 |
| 1.65 | 2,45E-05 | 6,41E-03 |
| 3.80 | 2,53E-05 | 6,51E-03 |
| 8.27 | 3,16E-05 | 7,74E-03 |
| 8.27 | 3,16E-05 | 7,87E-03 |
| 8.27 | 3,16E-05 | 7,99E-03 |
| 24.87 | 3,46E-05 | 8,34E-03 |
| 8.07 | 3,67E-05 | 8,71E-03 |
| 5.72 | 3,87E-05 | 9,05E-03 |
| 1.56 | 4,15E-05 | 9,56E-03 |
| 9.53 | 5,61E-05 | 1,26E-02 |
| 9.53 | 5,61E-05 | 1,27E-02 |
| 1.99 | 6,13E-05 | 1,35E-02 |
| 47.66 | 6,89E-05 | 1,50E-02 |
| 2.96 | 7,47E-05 | 1,58E-02 |
| 1.66 | 7,40E-05 | 1,59E-02 |
| 7.15 | 7,63E-05 | 1,60E-02 |
| 19.72 | 7,81E-05 | 1,61E-02 |
| 7.05 | 8,31E-05 | 1,69E-02 |
| 3.60 | 8,57E-05 | 1,72E-02 |
| 4.15 | 9,07E-05 | 1,80E-02 |
| 2.25 | 9,18E-05 | 1,80E-02 |
| 3.54 | 9,95E-05 | 1,93E-02 |
| 2.76 | 1,05E-04 | 2,01E-02 |
| 2.87 | 1,08E-04 | 2,04E-02 |
| 2.84 | 1,23E-04 | 2,29E-02 |
| 3.89 | 1,56E-04 | 2,89E-02 |
| 7.80 | 1,60E-04 | 2,92E-02 |
| 3.36 | 1,67E-04 | 3,01E-02 |
| 33.00 | 1,72E-04 | 3,03E-02 |
| 3.85 | 1,70E-04 | 3,03E-02 |
| 10.21 | 1,76E-04 | 3,07E-02 |
| 3.81 | 1,88E-04 | 3,25E-02 |
| 30.64 | 2,08E-04 | 3,54E-02 |
| 3.48 | 2,16E-04 | 3,65E-02 |
| 2.56 | 2,65E-04 | 4,42E-02 |
| 26.81 | 2,93E-04 | 4,84E-02 |
| 4.72 | 2,74E-23 | 1,36E-19 |
| 2.13 | 3,26E-16 | 8,10E-13 |
| 2.13 | 5,91E-16 | 9,80E-13 |
| 2.96 | 1,48E-11 | 1,47E-08 |
| 2.97 | 1,46E-11 | 1,81E-08 |
| 2.25 | 4,64E-11 | 3,30E-08 |
| 2.73 | 4,21E-11 | 3,49E-08 |
| 1.21 | 3,84E-09 | 1,73E-06 |
| 2.85 | 2,87E-09 | 1,79E-06 |
| 2.82 | 3,65E-09 | 1,82E-06 |
| 2.83 | 3,50E-09 | 1,94E-06 |
| 2.54 | 5,35E-09 | 2,22E-06 |
| 16.71 | 8,54E-09 | 3,27E-06 |
| 3.03 | 9,72E-09 | 3,45E-06 |
| 16.29 | 1,05E-08 | 3,48E-06 |
| 3.00 | 1,17E-08 | 3,63E-06 |
| 2.80 | 1,30E-08 | 3,80E-06 |
| 4.40 | 1,81E-08 | 5,01E-06 |
| 4.83 | 2,22E-08 | 5,81E-06 |
| 6.13 | 3,96E-08 | 9,86E-06 |
| 2.96 | 4,79E-08 | 1,13E-05 |
| 4.03 | 7,81E-08 | 1,62E-05 |
| 4.04 | 7,48E-08 | 1,69E-05 |
| 4.03 | 7,81E-08 | 1,69E-05 |
| 6.16 | 1,08E-07 | 2,16E-05 |
| 2.44 | 1,44E-07 | 2,75E-05 |
| 1.27 | 1,80E-07 | 3,31E-05 |
| 1.12 | 1,51E-06 | 2,60E-04 |
| .06 | 1,51E-06 | 2,69E-04 |
| 4.14 | 2,21E-06 | 3,66E-04 |
| 8.20 | 2,46E-06 | 3,94E-04 |
| 1.66 | 5,08E-06 | 7,66E-04 |
| 4.75 | 5,05E-06 | 7,85E-04 |
| 3.79 | 6,71E-06 | 9,82E-04 |
| 2.98 | 8,62E-06 | 1,23E-03 |
| 7.83 | 1,24E-05 | 1,71E-03 |
| 18.33 | 1,32E-05 | 1,77E-03 |
| 3.23 | 1,43E-05 | 1,87E-03 |
| 4.82 | 2,43E-05 | 3,10E-03 |
| 15.21 | 3,01E-05 | 3,66E-03 |
| 15.21 | 3,01E-05 | 3,75E-03 |
| 3.32 | 3,25E-05 | 3,85E-03 |
| 13.75 | 4,72E-05 | 5,46E-03 |
| < 0.01 | 4,90E-05 | 5,54E-03 |
| 20.43 | 6,90E-05 | 7,63E-03 |
| < 0.01 | 7,48E-05 | 8,09E-03 |
| 2.07 | 9,54E-05 | 1,01E-02 |
| < 0.01 | 1,14E-04 | 1,18E-02 |
| 2.04 | 1,54E-04 | 1,56E-02 |
| 7.59 | 1,84E-04 | 1,83E-02 |
| 2.40 | 2,00E-04 | 1,96E-02 |
| 9.41 | 2,53E-04 | 2,42E-02 |
| .09 | 3,39E-04 | 3,13E-02 |
| .09 | 3,39E-04 | 3,19E-02 |
| 95.32 | 4,73E-04 | 4,27E-02 |
| 7.94 | 5,29E-04 | 4,70E-02 |
| 1.37 | 1,36E-18 | 2,79E-15 |
| 1.46 | 1,93E-16 | 1,31E-13 |
| 1.40 | 2,85E-16 | 1,45E-13 |
| 1.53 | 1,52E-16 | 1,55E-13 |
| 6.01 | 4,30E-15 | 1,75E-12 |
| 1.44 | 6,32E-15 | 2,15E-12 |
| 1.50 | 1,78E-14 | 5,19E-12 |
| 2.11 | 2,81E-14 | 5,74E-12 |
| 2.11 | 2,81E-14 | 6,38E-12 |
| 2.11 | 2,81E-14 | 7,18E-12 |
| 1.98 | 5,54E-12 | 1,03E-09 |
| 1.97 | 2,89E-10 | 4,55E-08 |
| 1.97 | 2,88E-10 | 4,90E-08 |
| 2.79 | 9,69E-10 | 1,41E-07 |
| 2.76 | 1,35E-09 | 1,62E-07 |
| 2.76 | 1,35E-09 | 1,73E-07 |
| 2.76 | 1,33E-09 | 1,82E-07 |
| 1.97 | 1,36E-08 | 1,54E-06 |
| 3.25 | 2,05E-08 | 2,21E-06 |
| 8.71 | 2,59E-08 | 2,65E-06 |
| 1.81 | 6,34E-08 | 6,17E-06 |
| 5.00 | 8,10E-08 | 7,52E-06 |
| 3.42 | 1,62E-07 | 1,44E-05 |
| 12.71 | 4,14E-07 | 3,52E-05 |
| 12.43 | 4,84E-07 | 3,95E-05 |
| 1.98 | 5,86E-07 | 4,61E-05 |
| 15.64 | 6,18E-07 | 4,68E-05 |
| 13.71 | 1,40E-06 | 1,02E-04 |
| 2.57 | 1,67E-06 | 1,18E-04 |
| 4.78 | 2,03E-06 | 1,34E-04 |
| 2.54 | 2,00E-06 | 1,36E-04 |
| 4.73 | 2,26E-06 | 1,44E-04 |
| 4.62 | 2,94E-06 | 1,82E-04 |
| 12.06 | 3,11E-06 | 1,87E-04 |
| 47.66 | 3,81E-06 | 2,10E-04 |
| 47.66 | 3,81E-06 | 2,16E-04 |
| 1.10 | 3,73E-06 | 2,18E-04 |
| 1.51 | 5,47E-06 | 2,86E-04 |
| 6.41 | 5,46E-06 | 2,94E-04 |
| 10.11 | 9,25E-06 | 4,72E-04 |
| < 0.01 | 1,41E-05 | 6,85E-04 |
| 1.09 | 1,41E-05 | 7,02E-04 |
| 1.77 | 2,30E-05 | 1,09E-03 |
| 3.09 | 2,58E-05 | 1,20E-03 |
| 14.30 | 3,97E-05 | 1,80E-03 |
| 2.14 | 4,75E-05 | 2,11E-03 |
| 53.62 | 5,20E-05 | 2,17E-03 |
| 13.49 | 5,14E-05 | 2,19E-03 |
| 13.49 | 5,14E-05 | 2,23E-03 |
| 2.43 | 8,22E-05 | 3,36E-03 |
| 2.08 | 8,93E-05 | 3,58E-03 |
| 1.34 | 1,08E-04 | 4,23E-03 |
| 1.79 | 1,11E-04 | 4,27E-03 |
| 11.00 | 1,27E-04 | 4,71E-03 |
| 11.00 | 1,27E-04 | 4,80E-03 |
| 4.82 | 1,37E-04 | 5,00E-03 |
| 3.20 | 1,52E-04 | 5,46E-03 |
| 30.64 | 2,08E-04 | 7,19E-03 |
| 3.50 | 2,05E-04 | 7,22E-03 |
| 1.60 | 2,16E-04 | 7,36E-03 |
| 2.35 | 2,28E-04 | 7,64E-03 |
| 9.53 | 2,38E-04 | 7,86E-03 |
| 1.65 | 2,95E-04 | 9,55E-03 |
| 2.38 | 3,08E-04 | 9,69E-03 |
| 2.38 | 3,08E-04 | 9,85E-03 |
| 8.72 | 3,53E-04 | 1,09E-02 |
| 12.43 | 4,01E-04 | 1,19E-02 |
| 4.16 | 3,90E-04 | 1,19E-02 |
| 12.43 | 4,01E-04 | 1,20E-02 |
| 8.22 | 4,56E-04 | 1,31E-02 |
| 4.58 | 4,53E-04 | 1,32E-02 |
| 3.40 | 4,77E-04 | 1,33E-02 |
| 95.32 | 4,73E-04 | 1,34E-02 |
| 7.77 | 5,82E-04 | 1,61E-02 |
| 11.00 | 6,19E-04 | 1,64E-02 |
| 11.00 | 6,19E-04 | 1,66E-02 |
| 7.69 | 6,10E-04 | 1,66E-02 |
| 3.83 | 7,01E-04 | 1,81E-02 |
| 7.45 | 6,99E-04 | 1,83E-02 |
| 5.80 | 7,31E-04 | 1,87E-02 |
| 3.41 | 8,47E-04 | 2,11E-02 |
| 3.41 | 8,47E-04 | 2,14E-02 |
| 3.35 | 9,75E-04 | 2,37E-02 |
| 9.69 | 9,67E-04 | 2,38E-02 |
| 16.50 | 1,05E-03 | 2,52E-02 |
| 15.32 | 1,28E-03 | 3,04E-02 |
| 14.79 | 1,40E-03 | 3,30E-02 |
| 14.30 | 1,54E-03 | 3,53E-02 |
| 14.30 | 1,54E-03 | 3,57E-02 |
| 40.85 | 1,67E-03 | 3,71E-02 |
| 40.85 | 1,67E-03 | 3,75E-02 |
| 40.85 | 1,67E-03 | 3,79E-02 |
| 35.74 | 2,08E-03 | 4,57E-02 |

**Supplemental Table 5:**
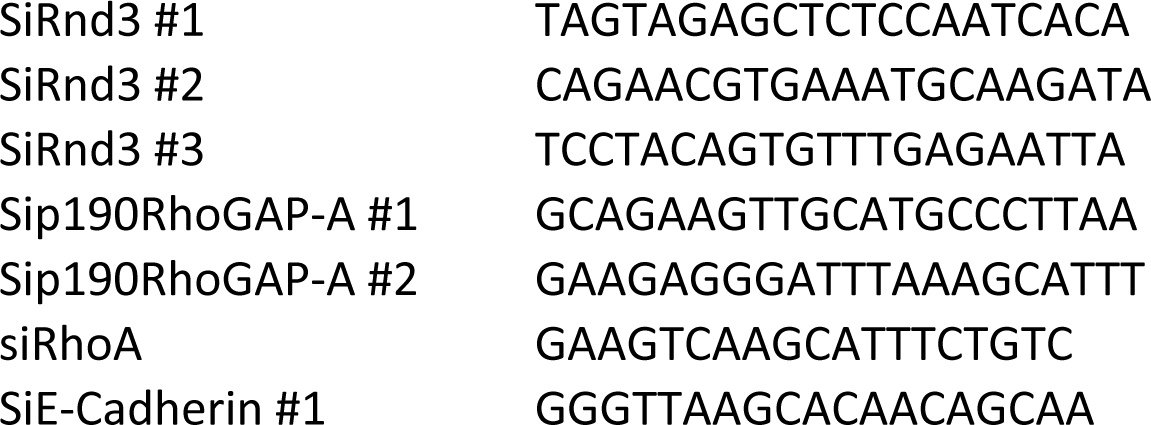
Sequences of siRNA used in this study.

### Videos

**Video 1**: The video shows the entosis mechanism upon Rnd3 silencing in Hep3B cells. Hep3B cell lines were transduced with H2B-GFP (yellow) and lifeact-mRuby (cyan) to mark the nucleus and the actin respectively. Cells were transfected with siRNA targeting Rnd3 and spinning disk microscopy analysis was done over 48 hours. Three stages were defined: i) cell contact, ii) internalization of the inner binuclear cell 21 hours after contact between cells and iii) beginning of the inner cell degradation 2 hours after internalization. The cell degradation takes almost 10 hours to be completed. Scale bar, 28 µm.

## REFERENCES

1. M. Schwegler, A. M. Wirsing, H. M. Schenker, L. Ott, J. M. Ries, M. Buttner-Herold, R. Fietkau, F. Putz, L. V. Distel, Prognostic Value of Homotypic Cell Internalization by Nonprofessional Phagocytic Cancer Cells. Biomed Res Int 2015, 359392 (2015).

2. H. Huang, M. He, Y. Zhang, B. Zhang, Z. Niu, Y. Zheng, W. Li, P. Cui, X. Wang, Q. Sun, Identification and validation of heterotypic cell-in-cell structure as an adverse prognostic predictor for young patients of resectable pancreatic ductal adenocarcinoma. Signal Transduct Target Ther 5, 246 (2020).

3. X. Zhang, Z. Niu, H. Qin, J. Fan, M. Wang, B. Zhang, Y. Zheng, L. Gao, Z. Chen, Y. Tai, M. Yang, H. Huang, Q. Sun, Subtype-Based Prognostic Analysis of Cell-in-Cell Structures in Early Breast Cancer. Front Oncol 9, 895 (2019).

4. B. Ruan, Z. Niu, X. Jiang, Z. Li, Y. Tai, H. Huang, Q. Sun, High Frequency of Cell-in-Cell Formation in Heterogeneous Human Breast Cancer Tissue in a Patient With Poor Prognosis: A Case Report and Literature Review. Front Oncol 9, 1444 (2019).

5. J. Fan, Q. Fang, Y. Yang, M. Cui, M. Zhao, J. Qi, R. Luo, W. Du, S. Liu, Q. Sun, Role of Heterotypic Neutrophil-in-Tumor Structure in the Prognosis of Patients With Buccal Mucosa Squamous Cell Carcinoma. Front Oncol 10, 541878 (2020).

6. A. Hayashi, A. Yavas, C. A. McIntyre, Y.-J. Ho, A. Erakky, W. Wong, A. M. Varghese, J. P. Melchor, M. Overholtzer, E. M. O’Reilly, D. S. Klimstra, O. Basturk, C. A. Iacobuzio-Donahue, Genetic and clinical correlates of entosis in pancreatic ductal adenocarcinoma. Mod Pathol 33, 1822– 1831 (2020).

7. Y. Wang, Z. Niu, L. Zhou, Y. Zhou, Q. Ma, Y. Zhu, M. Liu, Y. Shi, Y. Tai, Q. Shao, J. Ge, J. Hua, L. Gao, H. Huang, H. Jiang, Q. Sun, Subtype-Based Analysis of Cell-in-Cell Structures in Esophageal Squamous Cell Carcinoma. Front Oncol 11, 670051 (2021).

8. K. Borensztejn, P. Tyrna, A. M. Gaweł, I. Dziuba, C. Wojcik, L. P. Bialy, I. Mlynarczuk-Bialy, Classification of Cell-in-Cell Structures: Different Phenomena with Similar Appearance. Cells 10, 2569 (2021).

9. S. Fais, M. Overholtzer, Cell-in-cell phenomena in cancer. Nat Rev Cancer 18, 758–766 (2018).

10. A. S. Garanina, O. P. Kisurina-Evgenieva, M. V. Erokhina, E. A. Smirnova, V. M. Factor, G. E. Onishchenko, Consecutive entosis stages in human substrate-dependent cultured cells. Sci Rep 7, 12555 (2017).

11. M. Overholtzer, A. A. Mailleux, G. Mouneimne, G. Normand, S. J. Schnitt, R. W. King, E. S. Cibas, J. S. Brugge, A nonapoptotic cell death process, entosis, that occurs by cell-in-cell invasion. Cell 131, 966–79 (2007).

12. J. C. Hamann, A. Surcel, R. Chen, C. Teragawa, J. G. Albeck, D. N. Robinson, M. Overholtzer, Entosis Is Induced by Glucose Starvation. Cell Rep 20, 201–210 (2017).

13. R. Chen, A. Ram, J. G. Albeck, M. Overholtzer, Entosis is induced by ultraviolet radiation. iScience 24, 102902 (2021).

14. J. Durgan, Y. Y. Tseng, J. C. Hamann, M. C. Domart, L. Collinson, A. Hall, M. Overholtzer, O. Florey, Mitosis can drive cell cannibalism through entosis. Elife 6 (2017), doi:10.7554/eLife.27134.

15. Q. Sun, T. Luo, Y. Ren, O. Florey, S. Shirasawa, T. Sasazuki, D. N. Robinson, M. Overholtzer, Competition between human cells by entosis. Cell Res 24, 1299–310 (2014).

16. L. Paysan, L. Piquet, F. Saltel, V. Moreau, Rnd3 in Cancer: A Review of the Evidence for Tumor Promoter or Suppressor. Mol. Cancer Res. 14, 1033–1044 (2016).

17. S. Basbous, R. Azzarelli, E. Pacary, V. Moreau, Pathophysiological functions of Rnd proteins. Small GTPases, 1–22 (2020).

18. F. Grise, S. Sena, A. Bidaud-Meynard, J. Baud, J. B. Hiriart, K. Makki, N. Dugot-Senant, C. Staedel, P. Bioulac-Sage, J. Zucman-Rossi, J. Rosenbaum, V. Moreau, Rnd3/RhoE Is down-regulated in hepatocellular carcinoma and controls cellular invasion. Hepatology 55, 1766–75 (2012).

19. W. Ma, C. C. Wong, E. K. Tung, C. M. Wong, I. O. Ng, RhoE is frequently down-regulated in hepatocellular carcinoma (HCC) and suppresses HCC invasion through antagonizing the Rho/Rho-kinase/myosin phosphatase target pathway. Hepatology 57, 152–61 (2013).

20. S. Basbous, L. Paysan, S. Sena, N. Allain, J.-B. Hiriart, N. Dugot-Senant, B. Rousseau, E. Chevret, V. Lagrée, V. Moreau, Silencing of RND3/RHOE inhibits the growth of human hepatocellular carcinoma and is associated with reversible senescence. Cancer Gene Ther 29, 437–444 (2022).

21. G. Loor, P. T. Schumacker, Role of hypoxia-inducible factor in cell survival during myocardial ischemia-reperfusion. Cell Death Differ 15, 686–690 (2008).

22. Q. Sun, E. S. Cibas, H. Huang, L. Hodgson, M. Overholtzer, Induction of entosis by epithelial cadherin expression. Cell Res 24, 1288–98 (2014).

23. Z. Ezzoukhry, E. Henriet, F. P. Cordelières, J.-W. Dupuy, M. Maître, N. Gay, S. Di-Tommaso, L. Mercier, J. G. Goetz, M. Peter, F. Bard, V. Moreau, A.-A. Raymond, F. Saltel, Combining laser capture microdissection and proteomics reveals an active translation machinery controlling invadosome formation. Nat Commun 9, 2031 (2018).

24. S. Krishna, M. Overholtzer, Mechanisms and consequences of entosis. Cell Mol Life Sci 73, 2379– 86 (2016).

25. H. Luo, Z. Dong, J. Zou, Q. Zeng, D. Wu, L. Liu, Down-regulation of RhoE is associated with progression and poor prognosis in hepatocellular carcinoma. Journal of surgical oncology 105, 699– 704 (2012).

26. W. Ma, K. M. Sze, L. K. Chan, J. M. Lee, L. L. Wei, C. M. Wong, T. K. Lee, C. C. Wong, I. O. Ng, RhoE/ROCK2 regulates chemoresistance through NF-kappaB/IL-6/ STAT3 signaling in hepatocellular carcinoma. Oncotarget 7, 41445–41459 (2016).

27. J. Liang, Z. Niu, B. Zhang, X. Yu, Y. Zheng, C. Wang, H. Ren, M. Wang, B. Ruan, H. Qin, X. Zhang, S. Gu, X. Sai, Y. Tai, L. Gao, L. Ma, Z. Chen, H. Huang, X. Wang, Q. Sun, p53-dependent elimination of aneuploid mitotic offspring by entosis. Cell Death Differ 28, 799–813 (2021).

28. O. Florey, S. E. Kim, C. P. Sandoval, C. M. Haynes, M. Overholtzer, Autophagy machinery mediates macroendocytic processing and entotic cell death by targeting single membranes. Nat Cell Biol 13, 1335–1343 (2011).

29. A. K. Agarwal, N. Srinivasan, R. Godbole, S. K. More, S. Budnar, R. P. Gude, R. D. Kalraiya, Role of tumor cell surface lysosome-associated membrane protein-1 (LAMP1) and its associated carbohydrates in lung metastasis. J Cancer Res Clin Oncol 141, 1563–1574 (2015).

30. M. Krajcovic, N. B. Johnson, Q. Sun, G. Normand, N. Hoover, E. Yao, A. L. Richardson, R. W. King, E. S. Cibas, S. J. Schnitt, J. S. Brugge, M. Overholtzer, A non-genetic route to aneuploidy in human cancers. Nat Cell Biol 13, 324–30 (2011).

31. L. Piquet, T. Robbe, V. Neaud, S. Basbous, S. Rosciglione, F. Saltel, V. Moreau, Rnd3/RhoE expression is regulated by G-actin through MKL1-SRF signaling pathway. Exp. Cell Res. 370, 227– 236 (2018).

32. E. Henriet, A. Abou Hammoud, J.-W. Dupuy, B. Dartigues, Z. Ezzoukry, N. Dugot-Senant, T. Leste-Lasserre, N. Pallares-Lupon, M. Nikolski, B. Le Bail, J.-F. Blanc, C. Balabaud, P. Bioulac-Sage, A.-A. Raymond, F. Saltel, Argininosuccinate synthase 1 (ASS1): A marker of unclassified hepatocellular adenoma and high bleeding risk. Hepatology 66, 2016–2028 (2017).

33. E. W. Deutsch, N. Bandeira, Y. Perez-Riverol, V. Sharma, J. J. Carver, L. Mendoza, D. J. Kundu, S. Wang, C. Bandla, S. Kamatchinathan, S. Hewapathirana, B. S. Pullman, J. Wertz, Z. Sun, S. Kawano, S. Okuda, Y. Watanabe, B. MacLean, M. J. MacCoss, Y. Zhu, Y. Ishihama, J. A. Vizcaíno, The ProteomeXchange consortium at 10 years: 2023 update. Nucleic Acids Res 51, D1539–D1548 (2023).

34. M. Bou-Nader, S. Caruso, R. Donne, S. Celton-Morizur, J. Calderaro, G. Gentric, M. Cadoux, A. L’Hermitte, C. Klein, T. Guilbert, M. Albuquerque, G. Couchy, V. Paradis, J.-P. Couty, J. Zucman-Rossi, C. Desdouets, Polyploidy spectrum: a new marker in HCC classification. Gut 69, 355–364 (2020).

